# A contextualised protein language model reveals the functional syntax of bacterial evolution

**DOI:** 10.1101/2025.07.20.665723

**Authors:** Maciej Wiatrak, David C. Abelson, Roman Mikhaylenko, Ramon Viñas Torné, Maria Ntemourtsidou, Adam Dinan, Divya Arora, Samuel T. Horsfield, John Lees, Maria Brbić, Aaron Weimann, R. Andres Floto

## Abstract

Bacteria have evolved a vast diversity of functions and behaviours that are currently incompletely understood and poorly predicted from DNA sequence alone. To understand the syntax of bacterial evolution and discover genome-to-phenotype relationships, we curated over 1.3 million genomes spanning bacterial phylogenetic space, represented each as an ordered sequence of proteins, and used these sequences to train a transformer-based, contextualised protein language model, *Bacformer*. By pretraining on genome-wide evolutionary patterns, *Bacformer* captures the compositional and positional relationships of proteins and thereby provides a whole-genome framework for linking genomic organisation and content to measurable bacterial traits. We demonstrate the ability of *Bacformer* to accurately predict protein–protein interactions; uncover operon structure, which we validated experimentally; infer important phenotypic traits, including antimicrobial resistance, while revealing likely causal genes; and design template synthetic proteomes with desirable properties. Thus, *Bacformer* establishes a genomic foundation model that reveals the evolutionary rules governing bacterial gene organisation, function, and phenotype, opening a route to systematic whole-genome engineering.

## INTRODUCTION

Bacteria are arguably the most successful organisms on Earth, having evolved over millions of years to inhabit diverse geographic environments [1], biological niches [2, 3], and ecosystems [4]; acquired a vast metabolic repertoire to exploit diverse substrates, generate energy, and create biomass [5]; and developed complex behavioural traits such as virulence [6], immune evasion [7], and antibiotic resistance [8].

Bacterial genomes are known to encode these adaptations in a highly structured manner. For example: functionally related genes are usually found contiguously in operons [9]; transcription factors mostly regulate proximal rather than distant genes [10, 11]; and the genomic distance from the origin of replication generally predicts gene identity [12, 13]. Thus, the architecture of genomes may provide a framework to create informative representations of bacteria that could be used to understand the basis of evolution and enable generalisable functional predictions across taxa.

Previous large-scale analyses of bacterial genomes have focused on specific applications, such as biosynthetic gene cluster discovery [14, 15] or predicting antimicrobial resistance [16, 17, 18], and have been hampered by biases in public genome collections towards a few pathogenic bacteria [19, 20]. Moreover, existing methods such as DNA and protein language models (LMs) incorporate limited genomic context, ignoring the network of functional interactions encoded across entire bacterial genomes and their overall genomic organisation [21, 22, 23, 24, 25, 26].

We therefore sought to develop a foundation model of bacteria that captures the gene structure and function of entire genomes across the full spectrum of bacterial diversity. To achieve this, we leveraged and curated existing collections of over 1.3M bacterial metagenome-assembled genomes (MAGs) [27, 28], derived from environmental sequencing across diverse habitats to maximise phylogenetic and ecological diversity and encoding almost three billion proteins, to create a foundation model of bacterial life.

We represent individual bacterial genomes as ordered sequences of proteins and use these as inputs to train a transformer-based, contextualised protein language model (pLM), called *Bacformer*, which learns the functional organisation of bacterial genomes. By pretraining on genome-wide evolutionary patterns, *Bacformer* learns the compositional and positional relationships between proteins as emergent properties and performs strongly on core functional genomic tasks, including protein– protein interaction prediction and the inference of phenotypes such as antimicrobial resistance and carbon source utilisation. *Bacformer* can accurately reconstruct genomic organisation, including operon structure, and can also be used for generative modelling to design template synthetic genomes with specified properties. By modelling the genome as a sequence of protein embeddings, we create a whole-genome model that overcomes the context-length limits of DNA-based models and supports unified prediction and interpretation of genome-scale traits.

## RESULTS

### Training a genome-contextualised protein language model

To develop a foundation model that captures evolutionary processes across bacteria, we first curated a broad taxonomic corpus of over 1.3 million metagenome-assembled genomes (MAGs) from over 25, 000 species, spanning almost 1, 500 orders and more than 70 biomes (***Figure 1A***). To ensure a high-quality and representative training corpus, we applied quality-control filters to remove contaminated and highly fragmented genomes, while preserving broad taxonomic and environmental diversity (***Figure S1***; see *Supplementary Methods*).

**Figure 1.**
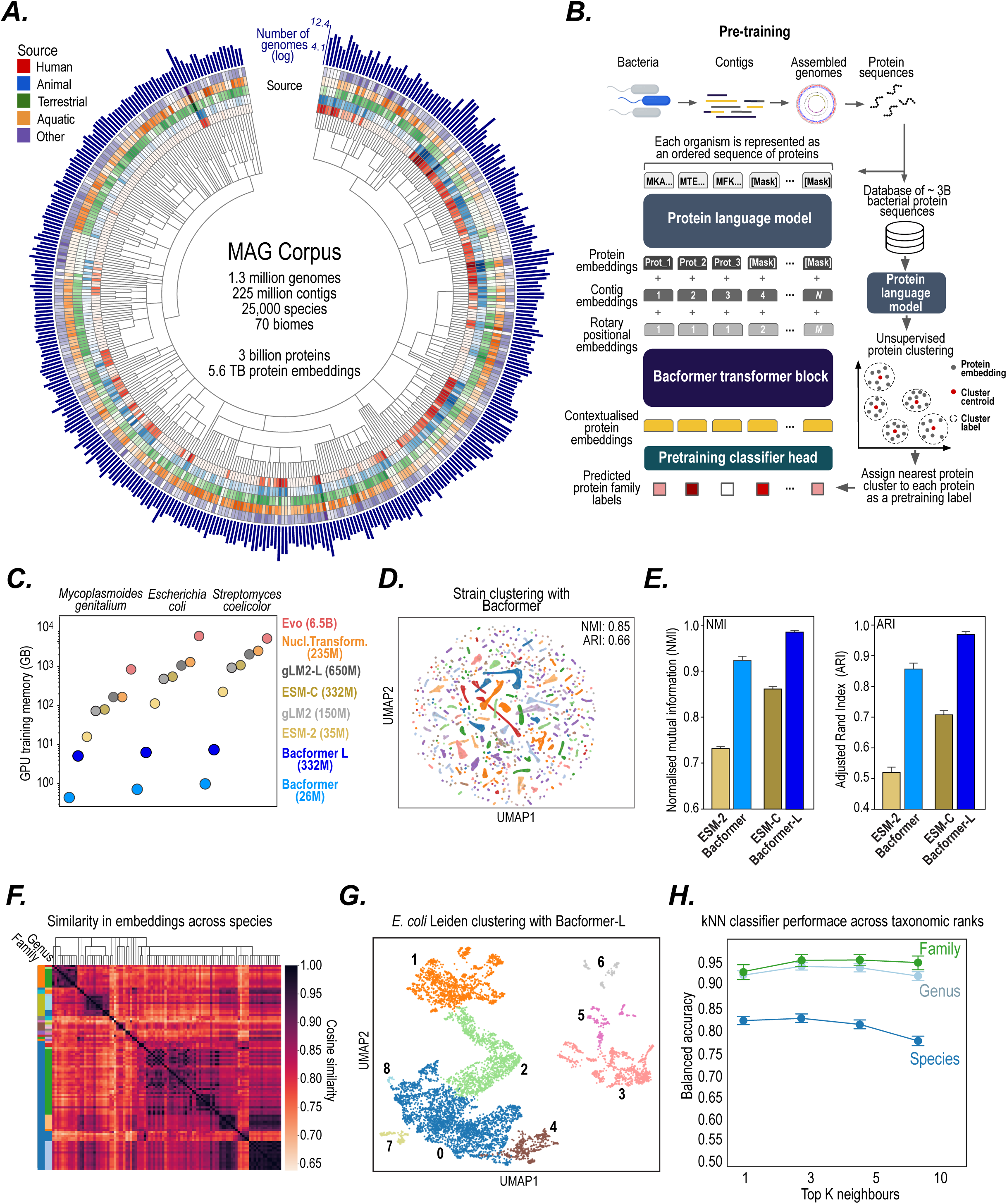
*Bacformer*, a bacterial foundation model, represents bacterial genomes as a sequence of contextualised protein embeddings. **(A)** To pretrain *Bacformer*, we constructed a metagenome-assembled genome (MAG) corpus, a dataset of more than 1.3 million genomes, 3 billion proteins, and 225 million contigs collected across 25, 000 species and 70 distinct environments. For visualisation, the taxonomy tree represents the 200 orders with the highest number of genomes in the MAG corpus, annotated by number of genomes (log; dark blue outer circle) and distribution across biospheres. **(B)** *Bacformer* model architecture and pretraining scheme. *Bacformer* takes as input the set of protein sequences present in a bacterial genome, ordered by genome location. Protein sequences are then embedded with a pretrained protein language model [21, 29] and combined with rotary positional embeddings [31] as well as learnable contig representations. The resulting protein representations are then used as input to the transformer, which computes contextualised protein embeddings using other proteins present in the genome. During pretraining, the model predicts the protein family of the masked protein based on the genomic context. The discrete vocabulary of protein-family labels necessary for pretraining was established by embedding non-contextual protein sequences with a protein language model and performing unsupervised clustering. **(C)** GPU training memory required to model 3 representative genomes using different genomic LMs. The required memory was calculated by generating the cost of the forward and backward pass with a batch size of 1 on distinct genomes. For all models, we used Flash attention and `bfloat16` precision on an NVIDIA A100 GPU (see *Supplementary Methods*). (D) *Bacformer* accurately clusters MAGs across diverse genera in an unsupervised manner. UMAP plot showing strain clustering using *Bacformer* embeddings of assemblies from the 10 most frequent genera in the MAG corpus, amounting to 314, 148 unique strains (clustering metrics: Adjusted Rand Index (ARI), Normalised Mutual Information (NMI); all metrics vary from 0 to 1, higher is better). **(E)** Performance of *Bacformer* and *Bacformer-L* compared to their base pLMs, *ESM-2* and *ESM-C*, respectively, on a species strain clustering benchmark task consisting of 6, 071 MAGs across 25 species. Metrics vary from 0 to 1, higher is better. Error bars denote 95% confidence intervals estimated from 10 independent bootstrap resamples, each using 80% of the data. (F) *Bacformer* representations of MAGs maintain evolutionary relationships. Cosine similarity heatmap for the 100 most common species of *Bacteroidales* showing higher similarity between species in the same family and genus. (G) *Bacformer-L* reveals subpopulations within a single species. UMAP plot of *Escherichia coli* MAGs from the *MGnify* dataset [27] using genome embeddings from *Bacformer-L* annotated with *Leiden* clustering labels (colours). (H) *k*NN classifier results across taxonomic ranks (family, green; genus, light blue; species, dark blue) demonstrate that *Bacformer* accurately identifies similar bacterial isolates. Error bars represent the 95% confidence interval across genomes.

Every individual assembled genome was then modelled as an ordered sequence of proteins, each of which is represented using existing protein language model (pLM) embeddings from the ESM family [21, 29] that capture evolutionary and functional properties of proteins [22, 21, 30, 29], supplemented with rotary positional embeddings [31]; an arrangement that preserves the genomic ordering of proteins (***Figure 1B***). To account for mobile elements, contig boundaries, and multiple chromosomes, we included special tokens that mark discontinuities in genomic sequence and preserve information about higher-order genome structure. *Bacformer*then utilises these enhanced embeddings of the entire proteome of an organism as inputs to a 12-layer (*Bacformer*) or a larger 30-layer (*Bacformer-L*) transformer model [32], treating each protein sequence as a single token and processing the genome in a single pass (see *Supplementary Methods*).

Unlike DNA- or protein-based LMs that rely on pre-defined and constrained token dictionaries [33, 25, 23, 21, 22], *Bacformer* accommodates the vast diversity of possible proteins by clustering them within embedding space into protein families and using these labels as a discrete vocabulary for the autoregressive or masked pretraining tasks, where the model learns to predict the family of masked proteins based on the context of other proteins in the genome (***Figure 1B***). To ensure that *Bacformer*’s protein family clusters used for pretraining reflect evolutionary relationships, we benchmarked them against established orthology databases (*eggNOG* [34]) and clustering methods (*MMseqs2* [35]), obtaining normalised mutual information (NMI) scores of 74% and 88%, respectively, indicating high agreement. Compared with KEGG functional annotations, *Bacformer* protein family clusters were highly functionally coherent (95.2% purity), with more than 99% of the clusters significantly enriched for specific KEGG categories after multiple-testing correction (BH-FDR, *q* < 0.05), indicating that the clusters capture biologically meaningful protein functions (see *Supplementary Methods*). Protein family clusters therefore provide a generalisable discrete vocabulary of functionally related proteins across taxa for pretraining. Importantly, the protein family clusters retained high within-genome diversity: the median ratio of unique clusters to proteins per genome was 0.91 ± 0.05, indicating that most proteins in each genome were assigned to distinct clusters (***Figure S1***). This allows *Bacformer* to learn relationships among many distinct functional units within each genome while representing each protein as a single token. *Bacformer*is designed to take the entire proteome as input, natively supporting finetuning for whole-genome tasks. In contrast, existing DNA- and protein-based LMs can only model full genomes by splitting them into segments, which requires orders of magnitude more GPU memory for finetuning, making this approach intractable to most users (***Figure 1C***).

During pretraining, *Bacformer* showed robust scaling behaviour: perplexity decreased as the training dataset grew (***Figure S2***), and validation loss improved with model size (***Figure S3***), demonstrating consistent trends observed in scaling transformer architectures [36, 37]. *Bacformer* also generalised to genomes not seen during training across multiple tasks and across a range of genome completeness bins (***Figure S2***). We further refined *Bacformer* by extending pretraining to include 48, 932 complete genomes [38], providing important additional contextual information that complements the broader MAG corpus. Having access to complete genomes boosted both zero-shot (i.e. no further training) and finetuned performance on tasks including protein–protein interaction and phenotypic trait prediction, highlighting the value of uninterrupted genomic context for the model (***Figure S3***).

### Genome embeddings uncover evolutionary relationships across bacteria

We next explored how the unified representation of bacterial genomes learned by *Bacformer* might provide insights into strain-level diversity and reveal subpopulations with distinct functional or evolutionary characteristics.

Existing clustering methods for bacteria operate only on closely related strains, rely on a limited set of marker genes and thus fail to consider gene content, or require computationally expensive pairwise sequence comparison [39, 40, 41, 42, 43, 44]. In contrast, *Bacformer* represents individual genomes across species in a shared embedding space (by averaging their contextual protein embeddings across the genome), enabling more robust strain-level comparisons based on the organisation of proteins within each isolate, and learning evolutionary relationships in an unsupervised way.

When applied to different sets of species, genera, and families (see *Supplementary Methods*), *Bacformer* accurately clustered genomes according to their taxonomic labels (based on Adjusted Rand Index (ARI), Adjusted Silhouette Width (ASW), and normalised mutual information (NMI)), considerably outperforming non-contextual pLM embeddings (*ESM-2* for *Bacformer* and *ESM-C* for *Bacformer-L*), other genomic LMs, and traditional clustering methods (AAI and ANI [44]), with *Bacformer-L* achieving mean scores of over 0.98 ±0.01 and 0.97 ±0.02 on NMI and ARI, respectively, compared to 0.86 ± 0.01 and 0.71 ± 0.02 achieved by the second-best-performing *ESM-C* (***Figure 1D,E***; ***Figure S4***), while also generalising to taxa not seen during training. Moreover, even in this fully unsupervised manner, *Bacformer* learned phylogenetic relationships. For example, for the *Bacteroidales* order, *Bacformer* embeddings produced average cosine similarities among species belonging to the same genus or to the same family of 0.96 ± 0.02 and 0.93 ± 0.05, respectively (***Figure 1F***).

To evaluate whether *Bacformer* could resolve fine-scale diversity within a single species and identify meaningful evolutionary differences, we examined the *Escherichia coli* sample collection from the *MGnify* dataset [27]. We extracted *Bacformer-L* embeddings for each MAG and applied *Leiden* clustering [45], which clearly identified nine distinct subpopulations, not driven by sampling variability that were not well delineated when we used *ESM-C* embeddings (***Figure 1G***; ***Figure S5***). We identified several core and accessory genes that were significantly cluster-specific (adjusted *p* < 0.05, Fisher’s exact test), including: *orgA* (in cluster 0) that is associated with type III secretion and host-cell invasion [46, 47], *fldP* (in cluster 1) that shields cells from oxidative stress and aids survival in low-oxygen environments [48, 49], and *gltB* (in cluster 2) that encodes the large subunit of glutamate synthase, supporting nitrogen assimilation [50] (***Figure S5***). *Bacformer-L* achieved high concordance with the established clustering methods average nucleotide identity (ANI) and graphical pangenome analysis *Panaroo* [51] (NMI > 80%), confirming that its embeddings recover fine-scale population structure captured by both sequence similarity and gene content-based approaches.

*Bacformer* embeddings can also be used to rapidly search for related genomes and infer evolutionary context, ecological niche and bacterial traits. We linked the 1.3M *Bacformer* genome embeddings to associated biological and clinical metadata to allow users to input an unannotated genome and retrieve similar genomes and associated traits from our database. We validated the effectiveness of this method by applying a *k*-nearest neighbour (*k*NN) classifier to a held-out subset of bacterial genomes, demonstrating highly accurate retrieval across taxonomic ranks (***Figure 1H***), with the method achieving almost 95% balanced accuracy across diverse genera.

Thus, *Bacformer*’s learnt representations of complete contextualised genomes can reveal important evolutionary signals at both broad phylogenetic and localised strain-specific scales, and can be used to infer bacterial properties of unseen genomes.

### Bacformer reveals functional organisation of proteins

Since *Bacformer* was trained to predict proteins from their genomic context, we anticipated that it would learn the evolutionary relationships between proteins and hence their organisation into functional networks.

To explore this potential emergent property, we first examined whether *Bacformer* implicitly learns genome-wide protein–protein interactions and hence would be able to predict functional protein networks. We evaluated zero-shot performance across a diverse set of interaction types and strains (2, 088) from *STRING* DB [52] by computing pairwise cosine similarities between protein embeddings present in the genome. Unlike traditional co-occurrence or phylogenetic profiling approaches, our strategy avoids multiple-sequence alignments and ad hoc distance cut-offs, yielding a single scalable similarity measure across proteins and species. *Bacformer* considerably outperformed its non-contextual protein language model base (*ESM-2*) across all tasks (***Figure 2A***; ***Figure S6***) and was further significantly improved by finetuning (*p*< 0.01, *t*-test; see *Supplementary Methods*). To assess the performance against other deep-learning methods, we benchmarked *Bacformer* models on a set of held-out strains. To test generalisation to phylogenetically distinct bacteria, we withheld entire genera during evaluation. *Bacformer* and *Bacformer-L* substantially outperformed other methods (***Figure 2B***), with *Bacformer-L* improving the zero-shot AUROC by 14 percentage points compared to its base pLM *ESM-C*, showing the importance of incorporating genomic organisation into the model architecture. To quantify the effect of genomic organisation, we compared PPI performance with retained versus randomly shu#ed gene order, and found that the AUROC dropped by 13 percentage points after shu#ing (*p* < 10^−5^; Wilcoxon signed-rank test), indicating that *Bacformer* heavily utilises genomic organisation across bacterial species (***Figure S7***).

**Figure 2.**
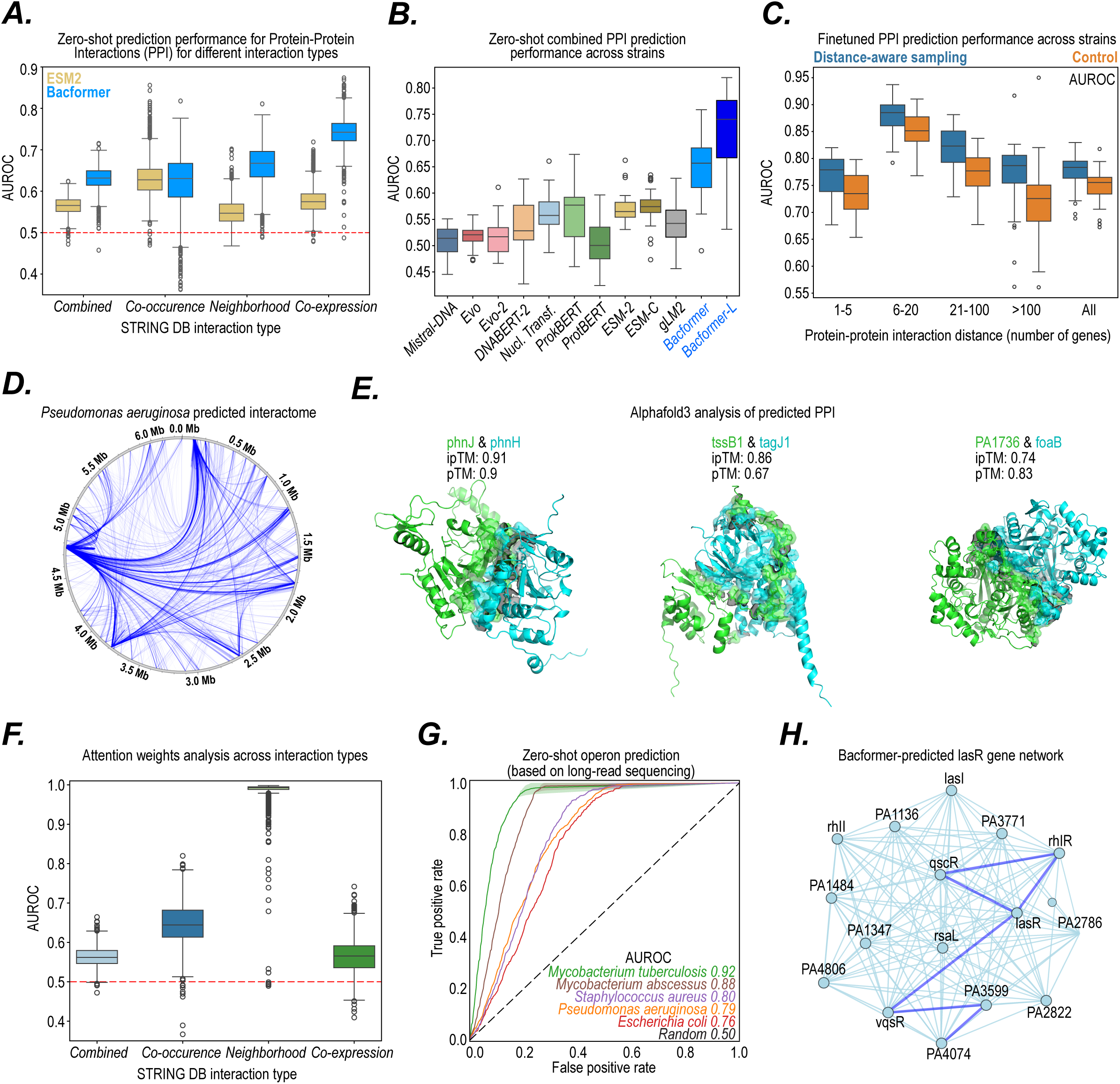
*Bacformer* identifies protein–protein interactions across bacterial genomes. **(A)** *Bacformer* learns protein–protein interactions in a zero-shot manner. Performance across different interaction types from *STRING DB* [52], outperforming its non-contextual protein embeddings (*ESM-2*, yellow) on held-out genomes (*N* = 2, 088). Lines and boxes represent median and interquartile range (IQR), respectively; whiskers extend 1.5⌐ IQR beyond Q1 and Q3; random classifier AUROC is shown as a red dashed line. **(B)** Benchmarking zero-shot performance across different genomic LMs on a smaller randomly sampled set of strains (*N* = 54) from *STRING DB*. Interaction scores were calculated using cosine similarity between protein representations on held-out genomes (see *Supplementary Methods*). Lines and boxes represent median and interquartile range (IQR), respectively; whiskers extend 1.5⌐ IQR beyond Q1 and Q3. **(C)** Performance on the protein–protein interaction task across distance bins (measured by number of genes) using distance-aware sampling versus control finetuned *Bacformer-L* across a distinct set of strains (*N* = 54). To test generalisation to phylogenetically distinct bacteria, we withheld entire genera during evaluation. Lines and boxes represent median and interquartile range (IQR), respectively; whiskers extend 1.5⌐ IQR beyond Q1 and Q3. **(D)** Proteome-wide interaction scores for *Pseudomonas aeruginosa* PAO1 predicted from finetuned *Bacformer-L* (line thickness indicates prediction strength; only predictions with probability > 0.5 were visualised). **(E)** Predicted structures obtained from *AlphaFold3* [53] for top-scoring protein complexes from *P. aeruginosa* PAO1 predicted by *Bacformer* (*ipTM* score indicates the probability of the two proteins binding together and *pTM* score is a measure of overall predicted structure confidence; range for both 0–1, higher is better). **(F)** Zero-shot protein-protein interaction prediction results using attention weights (last layer, average of all attention heads) as interaction scores across interaction types from STRING DB across *N* = 2, 088 genomes using area under the receiver operating characteristic (AUROC). Line and boxes represent median and interquartile range (IQR) respectively; whiskers extend 1.5 × IQR beyond Q1 and Q3; random classifier AUROC (red dashed line). **(G)** *Bacformer* zero-shot operon predictions (assessed by AUROC) for five bacterial species where we generated experimental ground truth data from long-read RNA sequencing. **(H)** Protein–protein interaction network extracted using *Leiden* clustering (resolution = 40) [45] on *Bacformer* contextualised protein embeddings. Edge weights with high importance (cosine similarity > 0.8) are highlighted in dark blue.

Bacterial PPIs are unevenly distributed across genomes. Interacting proteins are disproportionately encoded by neighbouring genes, reflecting evolutionary processes (***Figure S7***), causing a proximity bias that is also present in reference interaction datasets and could lead models to rely on genomic distance rather than interaction-specific biological signals. To account for this, we developed a distance-aware sampling strategy that bins protein pairs by genomic separation and samples equal numbers of interacting and non-interacting pairs from each bin during training. We then finetuned *Bacformer-L* trained with this balanced sampling strategy, which outperformed an unbalanced control across bins (*p* < 10^−10^; Wilcoxon signed-rank test, ***Figure 2C***), highlighting the importance of explicitly accounting for genomic-distance bias when evaluating PPI prediction.

When applied to *P. aeruginosa*, the finetuned model uncovered previously known, as well as novel, high-confidence protein–protein interactions (***Figure 2D***). We then used *AlphaFold3* [53] to validate novel physical interactions by predicting the structure of protein complexes (***Figure 2E***). Protein complexes predicted by *Bacformer-L* had significantly higher *ipTM* scores (a metric that evaluates the accuracy of predicted protein–protein interfaces [54]) than complexes based on distance-matched random pairs (*p* < 0.004; one-sided distance-stratified permutation test; ***Figure S7***). Thus, *Bacformer-L*-predicted interactions may reflect direct physical binding, although further experimental validation is required.

To better understand how *Bacformer* supports these interaction predictions, we examined the model’s attention weights across thousands of genes in diverse genomes and found that *Bacformer* focuses on neighbouring genes. While early layers distribute attention evenly, middle layers concentrate attention strongly on the closest neighbouring genes, whereas later layers spread attention further along the chromosome (***Figure S8***). We then examined whether the attention weights capture known protein–protein interactions (PPI) on a set of 2, 088 bacterial genomes across 1, 573 species using different types of interaction score (combined, co-occurrence, neighbourhood, co-expression) from the *STRING* database [52] as binary labels (***Figure 2F***). *Bacformer* attention weights recovered known PPIs, performing particularly well for neighbourhood (0.99 ± 0.03) and co-occurrence (0.64 ± 0.05) median AUROC scores, indicating that the model captures both conserved local gene organisation and broader evolutionary co-retention of functionally linked proteins.

We therefore tested whether *Bacformer* could predict the presence of operons (physically adjacent, co-transcribed genes that represent the fundamental regulatory units of bacterial genomics [9]). Since current operon annotations are incomplete and potentially inaccurate, we performed long-read RNA sequencing of strains from five bacterial species (*Mycobacterium tuberculosis*, *Mycobacterium abscessus*, *Staphylococcus aureus*, *Pseudomonas aeruginosa*, and *Escherichia coli*; see *Supplementary Methods*). We then established a method for obtaining zero-shot operon predictions that leverages representations from the *Bacformer* model and genomic features such as strand orientation. *Bacformer*achieved zero-shot AUROC of 0.76–0.92 when evaluated against experimentally validated operons across strains (***Figure 2G***), demonstrating how the model captures genome organisation without task-specific training. When compared with other methods, *Bacformer* representations outperformed them, illustrating the benefits of incorporating genomic organisation into the model architecture (*Tables S1,S2*).

*Bacformer* was also able to identify broader regulatory networks using its embedding similarity matrix, even without direct input of transcriptional information. For example, in *P. aeruginosa*, *Bacformer* was able to predict clusters of co-expressed genes more accurately than *ESM-2* (***Figure S7***) and identify networks mediating canonical quorum-sensing [55], biofilm formation [56], and motility [57] (***Figure 2H***; ***Figure S7***), showcasing how *Bacformer* embeddings can reveal co-evolved, co-regulated, and interacting protein families.

### Predicting protein essentiality and functions

To explore how kingdom-wide evolutionary signals and their genome-wide context learnt by *Bacformer* might reveal protein characteristics, we first examined its ability to predict protein essentiality. To create a ground truth dataset, we compiled genomes from 55 bacterial genomes across 37 species (from the *DEG* database [58]), providing a total of 193, 213 genes labelled as essential or non-essential. We then examined the performance of *Bacformer* and *Bacformer-L* in predicting protein essentiality using a phylogeny-aware split in which entire genera were held out for testing, allowing us to assess generalisation to unseen taxa. A linear model trained on top of the *Bacformer* embeddings was able to clearly separate essential from non-essential genes based on predicted probability distributions, achieving 0.81 ± 0.07 median AUROC across diverse genomes (***Figure S9***), and was able to identify many genes not previously annotated as essential (which we scored as false positives) but with orthologs in other species that are known to be essential, such as ribosomal proteins [59, 60] and tRNA synthetases [61, 62] (*Tables S3,S4*), indicating that our model issued biologically plausible predictions.

Importantly, we found that *Bacformer* was still able to make confident predictions on gene essentiality even when we masked the query protein itself (0.64 median AUROC across held-out genomes, ***Figure S9***), indicating the importance of learned contextual information for model predictions.

We then evaluated the ability of *Bacformer* and *Bacformer-L* to predict protein function. We leveraged KEGG [63] to extract protein function labels across diverse genomes (see *Supplementary Methods*), creating a dataset of 1, 126, 510 proteins across 300 genomes and 9, 059 unique protein function labels (KEGG KO orthology groups). Using the dataset, we trained a linear layer on top of the protein embeddings to predict the function of each protein. *Bacformer*and *Bacformer-L* achieved balanced accuracy of over 94% across held-out test strains, increasing to over 98.5% after finetuning the entire model (***Figure S10***). To demonstrate the functionality we then annotated hypothetical as well as other unannotated proteins, allowing us to confidently annotate 98, 821 proteins across 60 held-out strains, demonstrating how the *Bacformer* models can be used for protein function inference (***Figure S10***).

### Genome-scale inference of bacterial traits and ecological niche

By modelling entire bacterial genomes, *Bacformer* links genome-wide organisation to bacterial traits and ecological niches. We found that *Bacformer* genome embeddings intrinsically reflect phenotypic traits. For example, isolate embeddings clustered by Gram stain (***Figure S12***) and by host environment, even when comparing closely related species, for example within the *Bifidobacterium* genus (***Figure 3A***). We therefore assessed whether *Bacformer*, even without finetuning, could predict selected bacterial traits such as metabolism, carbon source utilisation or oxygen requirement.

**Figure 3.**
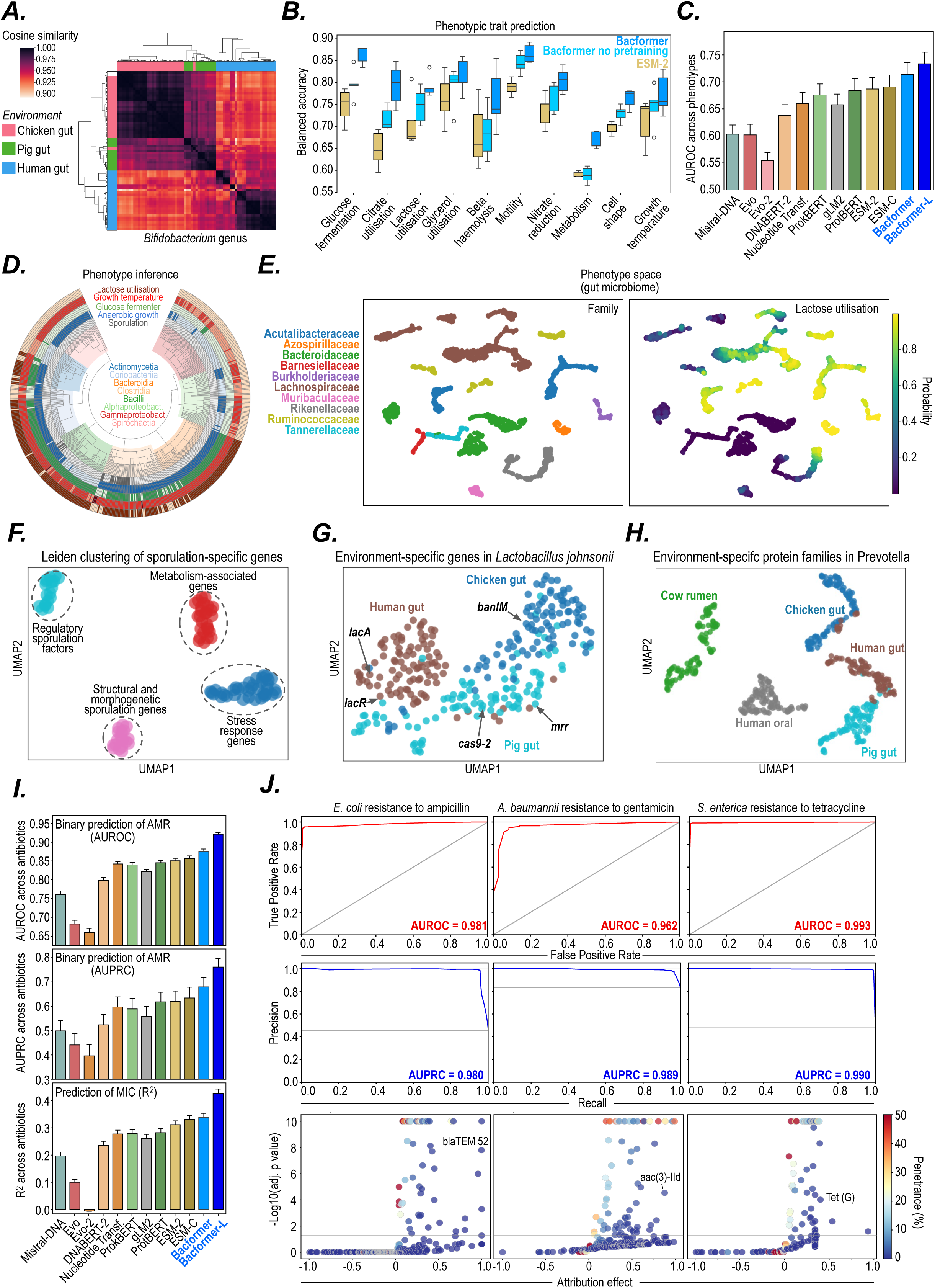
*Bacformer* enables prediction of selected phenotypic traits and prioritisation of trait-associated genes. **(A)** *Bacformer* genome embeddings capture environmental niche in an unsupervised manner. Cosine similarity heatmap between strains from the *Bifidobacterium* genus showing higher similarity between strains from the same animal niche (chicken gut, pink; pig gut, green; human gut, blue). The dendrogram was created by performing hierarchical clustering on *Bacformer* genome embeddings. **(B)** Selected trait prediction performance of pretrained *Bacformer* (dark blue), *Bacformer* without pretraining (light blue), and non-contextual *ESM-2* (yellow) shown for a range of curated traits. Lines and boxes represent median and interquartile range (IQR), respectively; whiskers extend 1.5⌐ IQR beyond Q1 and Q3. **(C)** Benchmarking performance across methods on 138 distinct traits across random seeds as evaluated by the area under the receiver operating characteristic (AUROC). For each method-trait combination, we trained a linear layer on top of the genome embedding, splitting the data by genus to ensure generalisation across phylogenetically different bacteria (see *Supplementary Methods*). Error bars represent 95% confidence intervals across traits and random seeds. **(D)** Example of selected trait inference across bacterial classes. Inner circle: cladogram of a selection of bacterial genomes (sampled from the MAG corpus), colour-coded by class. Outer circles: a subset of traits (from a total of 138) inferred using trait-specific models trained using *Bacformer* genome embeddings. **(E)** UMAP representation of the microbial community from a human gut sample created using predicted trait scores. Each point is a MAG. MAGs belonging to species represented by at least 50 genomes were extracted from a human gut metagenomic sample, resulting in 2, 132 unique MAGs. The MAGs were then assigned *Bacformer*-predicted scores for 138 phenotypic traits. The UMAP was computed by performing PCA followed by UMAP on the matrix of MAGs by predicted trait scores and coloured by family and lactose utilisation. **(F)** UMAP plot of the 100 genes with the highest importance scores for the sporulation trait across genomes (using their embeddings extracted from the last layer of the *Bacformer* model and averaged across genomes) and annotated by *Leiden* clustering [45] label and function (function labels were established by analysing the genes within each Leiden cluster). **(G)** The top 100 genes from *Lactobacillus johnsonii* genomes with the highest importance scores for each environmental niche (human gut, brown; chicken gut, dark blue; pig gut, light blue; i.e. isolation source) and plotted as a UMAP (using gene embeddings extracted from the last layer of the *Bacformer* model and averaged across genomes). Some genes with known association to the environmental niche are labelled. Genomes and environmental niche labels provided by the *MGnify* dataset [27]. **(H)** UMAP plot of the top 100 protein families (i.e. clusters) with the highest importance scores for each environmental niche (cow rumen, green; human oral, grey; chicken gut, dark blue; human gut, brown; pig gut, light blue) across *Prevotella* genus genomes using protein family embeddings extracted from the last layer of the *Bacformer* model and averaged across genomes. Genomes and environmental niche labels provided by the *MGnify* dataset [27]. **(I)** Benchmarking AMR prediction performance across methods on distinct drugs across random seeds. For each method-drug combination, we trained a linear layer on top of the genome embedding. We evaluated performance in the binary (top and middle; predicting susceptible/resistant labels) and regression setting (bottom; predicting minimum inhibitory concentration; see *Supplementary Methods*). **(J)** Performance of finetuned *Bacformer* model on 3 distinct species-drug combinations on the area under the receiver operating characteristic curve (AUROC; top) and precision-recall (AUPRC, middle) curves. Error bars represent 95% confidence intervals across drugs and random seeds. Volcano plot demonstrating gene element importance discovered by *Bacformer*, annotated by *AMR Finder Plus* [17]. The *x*-axis shows the model attribution (rank-biserial statistic derived from *Mann–Whitney* U rank-sum test with higher values indicating greater positive attribution compared to background). The *y*-axis displays statistical significance (− log_10_ adjusted *p*-value; one-sided test for element attribution > background). Colours represent the prevalence of each annotated element across all genomes in the species. Details of each gene score are provided in *Tables S6, S7, S8*.

We compiled a dataset of 24, 462 genomes from 15, 477 species that had been annotated with 138 distinct phenotypic traits [64, 65, 66] and trained separate classifiers to predict each trait from *Bacformer*genome-level embeddings (***Figure 3B***) computed by averaging all of the contextual protein embeddings present in the genome. For benchmarking, we used phylogeny-aware genus-level splits, holding out entire genera during testing to assess generalisation to unseen taxa. *Bacformer* and *Bacformer-L* predicted a range of specific traits, including carbon source utilisation, optimal growth temperature, motility, and toxin production, outperforming state-of-the-art deep-learning methods, including their non-contextual ESM base models. Moreover, *Bacformer* without pretraining also outperformed its base *ESM-2* model, demonstrating the benefits of incorporating genomic organisation with positional embeddings (***Figure 3B***). These comparisons demonstrate that both genome-wide context and pretraining substantially improve phenotype prediction. We then employed those phenotype-specific models with high predictive performance (balanced accuracy > 85%) to infer annotations for the entire 1.3M genomes within the MAG corpus across measured traits spanning metabolic, ecological, and physiological characteristics (***Figure 3D***; see *Supplementary Methods*). We provide these models, trait annotations, and accompanying metadata as a large-scale resource for inference of these bacterial phenotypes, which researchers can explore, filter, or extend for their own studies.

Given the breadth of trait information that *Bacformer*can infer across a large phylogenetic spread of bacteria, we asked whether these predictions could help characterise the functional composition of polymicrobial communities. As an exemplar, we annotated a single human gut microbiome sample containing 5, 811 MAGs with predictions for all 138 traits and used these scores to compute a UMAP plot, labelling genomes by both bacterial family and predicted lactose utilisation (***Figure 3E***). This analysis highlights predicted trait variability within families and suggests potential functional differences among community members.

The accuracy of *Bacformer*in predicting phenotypes motivated us to further leverage its contextual genomic representations to prioritise genes putatively associated with those traits by employing a gradient-based attribution method [67] that assigns importance scores to genes in a genome. Uniquely, *Bacformer* enables finetuning using whole-proteome context as input, allowing us to identify candidate phenotype-associated genes across diverse genomes.

We first examined the ability of this method to prioritise genes related to sporulation, motility, and anaerobic growth (three traits with some pre-existing knowledge of associated genes [68, 69, 70, 63]), and found that known trait-associated genes were assigned higher importance scores than other genes (*p* < 10^−10^, Mann–Whitney test; ***Figure S12***).

We then examined in detail the sporulation-associated genes identified by *Bacformer* and found that: the canonical sporulation genes, *spoIIIAG* and *spoIIAF* [71, 72], had been ascribed the highest importance scores; and that clustering the top 100 sporulation-associated genes revealed functionally distinct groups including known regulatory factors (*sigG*, *spoIIAB* [73, 74]) and structural genes (*spoVE*, *spoVB* [75, 76]), as well as plausible candidate genes, such as *katB* [77, 78] and *galE* [79] (***Figure 3F***).

We next tested whether we could use *Bacformer* to identify niche-specific genes. Within *Lactobacillus johnsonii*, a gut bacterium found in several animal species [80, 81], *Bacformer* was able to identify known genes involved in host-specific adaptation, such as *lacA* and *lacR* (human, [80]), *cas9* genes in livestock [82, 83], type III RM gene (*mmr*) (pig [82]), and *banIM* (chicken, [84]), as well as a putative set of additional novel genes (***Figure 3G***; *Table S5*). Finally, applying a similar approach to the *Prevotella*, a genus for which gene annotations and experimental validations are sparse, *Bacformer* was able to identify novel protein families specifically associated with gastrointestinal environments in different animal species (***Figure 3H***).

Thus, by combining large-scale prediction of traits with high-resolution gene-level attribution, *Bacformer* can not only assign specific traits across bacteria but can also discover potentially causally associated genes.

### Predicting antimicrobial resistance (AMR) and AMR-associated genes

We next evaluated whether *Bacformer* could predict antimicrobial resistance (AMR). Using an existing dataset of isolate genome sequences from 37 pathogenic bacterial species and their response to 54 different antibiotics (NCBI AST Browser [85]), we trained a separate linear classifier and regression model for each antibiotic using the genome-level embeddings of each isolate (see *Supplementary Methods*). *Bacformer* and *Bacformer-L* markedly outperformed other DNA and protein language models on both binary classification and regression tasks (***Figure 3I***). *Bacformer* models also substantially outperformed their base pLMs, with *Bacformer-L* improving AUPRC by over 12 percentage points across drugs, highlighting the benefit of modelling contextual protein interactions rather than using a bag-of-proteins representation. Consistently, randomising protein order reduced performance by over 12 percentage points in AUPRC (*p* < 10^−10^; Wilcoxon signedrank test; ***Figure S11***). Thus, gene order, rather than purely gene presence or absence, is important for the AMR prediction task.

In contrast to other genomic language models which only consider short context windows [25, 33, 21, 23, 26, 24], *Bacformer* is specifically designed to model whole genomes and consequently can be readily finetuned for organism-level phenotypic trait prediction tasks. To illustrate this property, we finetuned *Bacformer* to predict resistance for three distinct combinations of bacterial species and classes of antibiotics, achieving high AUROC and AUPRC performance (***Figure 3J***), even with training sets of less than 800 isolates. Because *Bacformer* predicts AMR from whole-genome context rather than testing resistance genes in isolation, attribution can highlight known determinants as well as accessory, regulatory, or epistatic loci, enabling prioritisation of AMR-associated candidate genes (***Figure 3J***; *Tables S6,S7,S8*).

### Generating de novo bacterial proteomes

Since *Bacformer* has learnt genomic representations of bacteria that encompass evolutionary relationships, reflect protein functional organisation, and support inference of bacterial phenotypes, we next looked to evaluate its potential for synthesising bacterial proteomes *de novo*.

We first evaluated *Bacformer*’s generative capabilities by analysing its performance on complete genomes from taxa not seen during training (*N* = 2, 086). *Bacformer* achieved a low perplexity (a measure of uncertainty when predicting the next token; *Bacformer* median = 9.12 vs random = 50, 000) across diverse unseen taxa (***Figure S3***), suggesting it captures the genomic architecture of strains not encountered in the training set. We then prompted *Bacformer* with a short sequence of initial proteins from *Escherichia coli* and *Staphylococcus aureus* and asked it to generate a complete genome (see *Supplementary Methods*). The resulting ordered sequence of protein families within synthesised genomes was comparable in distribution and ontological diversity to those of true genomes (***Figure 4A,B***; ***Figure S13***), demonstrating *Bacformer*’s capacity to create genomes that span the full spectrum of protein functions. To ensure that the model did not simply recapitulate genomes seen during training, we generated multiple genomes from identical protein prompts while varying the temperature (a scaling factor in autoregressive sampling that controls randomness) and observed that *Bacformer* generated a variety of distinct genomes, none of which were replicas of any within the training corpus (***Figure S13***).

**Figure 4.**
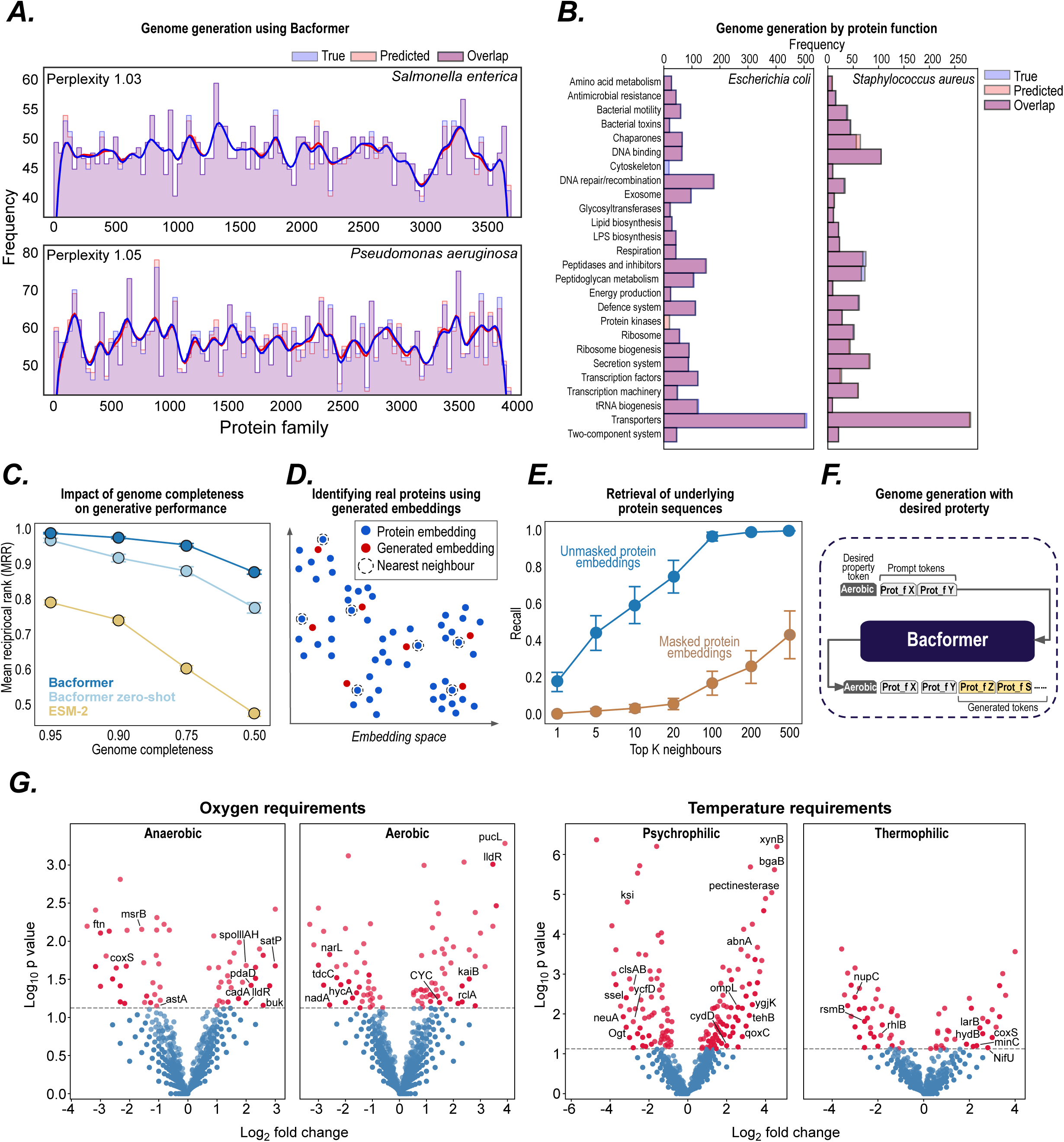
*Bacformer* generates bacterial proteome templates conditioned on specified properties. **(A)** *Bacformer* successfully generates genomes not seen during training. Next protein family prediction (orange) on *Salmonella enterica* (top) and *P. aeruginosa* (bottom) compared to true (blue) protein family distributions. **(B)** *Bacformer* generates genomes encompassing a realistic spectrum of protein functions (predicted, orange; true, blue) for *E. coli* (left) and *S. aureus* (right). We prompted *Bacformer* with a short sequence of initial proteins (*N* = 500) and evaluated the generated genomes against true labels. **(C)** *Bacformer* retrieves true complete genomes from incomplete genomes. Mean reciprocal rank (MRR; 0–1, higher is better) of genomes generated from inputs with ablated proteins by zero-shot (light blue) and non-zero-shot finetuned *Bacformer* (dark blue) and *ESM-2* (green). Error bars represent 95% confidence intervals. **(D)** Visualisation showing the process of identifying existing proteins from *Bacformer*-generated contextual protein embeddings using next-protein family prediction. **(E)** Retrieval of underlying protein sequences using unmasked (blue) and masked (brown) contextualised protein embeddings across a range of *k* neighbours. Error bars show standard error. **(F)** *Bacformer* generates genome templates (defined as sequences of protein families) conditioned on specified properties. We finetuned *Bacformer* to generate protein family sequences conditional on a property (represented as a token appended to the beginning of the sequence). The prompt includes the property token and a set of initial protein families present in the genome that are used by the model to generate the rest of the genome. **(G)** We generated genome templates conditioned on different specified phenotypic traits (exemplars shown are oxygen requirement and optimal growth temperature). We then compared sequences produced with and without the property token, calculated the log_2_ fold change for *KEGG* orthology (KO) groups [63], and assessed statistical significance with Fisher’s exact test.

We then tested whether *Bacformer* could accurately retrieve the full genome from an incomplete set of proteins, a valuable capability particularly for partial genomes typically recovered from metagenomic sequencing approaches. We therefore artificially removed a percentage of proteins from a collection of held-out complete genomes and computed their genome embeddings compared to the complete genomes, calculating the mean reciprocal rank (MRR). *Bacformer* was able to extrapolate the full genomic context even when only 50% of proteins were present, demonstrating the ability to confidently infer the whole proteome from sparse protein sets (***Figure 4C***; ***Figure S14***).

Since *Bacformer* was trained to predict protein families rather than specific proteins, we augmented its training with a contrastive loss to allow matching contextual protein embeddings from *Bacformer* to protein sequences, encouraging the model’s contextual embeddings to align closely with actual protein embeddings present in our database (see *Supplementary Methods*), thereby enabling retrieval of specific protein sequences from generated contextual protein embeddings (***Figure 4D***). To do this, we finetuned the entire *Bacformer* model on a non-causal protein prediction task (where 15% of proteins were masked), combining the masked objective with a contrastive loss. The resulting embeddings from *Bacformer* were significantly more similar to their true protein counterparts than to other proteins (*p* < 10^−10^; ***Figure S14***), demonstrating that the model can retrieve the underlying protein sequence from contextualised protein representations, including when the protein identity must be inferred from genomic context alone (***Figure 4E***).

Finally, we explored how *Bacformer* could be used to generate bacterial genome templates conditioned on specified traits, a core ambition of synthetic biology [86, 87, 88]. Leveraging our phenotypic trait analysis (***Figure 3***), we introduced a property token at the start of each protein-family sequence (***Figure 4F***), and then finetuned *Bacformer* to generate protein-family sequences conditional on a specified trait, selecting oxygen requirement and optimal growth temperature to evaluate its generative capacity.

We then used each trait-conditioned model to generate genome templates consisting of protein family tokens from an initial set of protein families drawn from held-out species and observed functional enrichment of proteins with biological roles linked to the specified trait (***Figure 4G***). For example: conditioning on aerobic growth produced significant positive enrichment of urate oxidase (*pucL*) and cytochrome c family genes (*CYC*), both of which catalyse key oxidative steps in purine catabolism and electron transport during aerobic respiration [89, 90], and significant depletion of proteins that promote anaerobic metabolism (such as the nitrate-responsive regulator *narL* and formate-hydrogen lyase component *hycA* [91, 92]). Conditioning on psychrophilic growth yielded significant enrichment of cold-active xylanases (*xynB*) and ω-galactosidases (*bgaB*), enzymes commonly isolated from polar and deep-sea bacteria where they facilitate polysaccharide degradation at low temperature [93, 94], and significant depletion of genes (such as steroid ε-isomerase (*ksi*) [95]). These findings indicate the potential utility of *Bacformer* for rational genome design.

## Discussion

*Bacformer*, a genomic foundation model trained on millions of diverse bacterial strains, offers a unified representation of genomes, modelled as an ordered sequence of interacting proteins, and generalises across species to incorporate the constraints and relationships that shape proteomes during evolution.

During pretraining, *Bacformer* generates informative whole-genome embeddings that identify biologically important clusters of bacteria and their associated gene content, and can be used to rapidly annotate unknown genomes by relating them to existing isolates with known traits. Importantly, *Bacformer* exhibits emergent properties by learning both protein–protein interactions and protein functional organisation, enabling accurate prediction of operons, protein annotation, strain clustering, and the syntactic rubric underpinning bacterial evolution and functional diversification, thereby supporting biological discovery.

Leveraging these emergent properties, we have trained *Bacformer* to predict bacterial phenotypes, including antimicrobial resistance, and identify putative causal gene modules associated with them, thereby revealing important biological processes and permitting phenotype inference within or across species at microbiome and ecological scale. *Bacformer* can also be used for generative tasks, inferring complete genomes from incomplete protein sets and creating a roadmap for *de novo* genome-template synthesis conditioned on specified properties, with potential applications to synthetic biology and bioengineering.

In summary, *Bacformer* demonstrates that bacterial genomes can be modelled as structured, genome-scale systems in which protein content, order, and neighbourhood encode evolutionary and functional information, providing a framework for whole-genome representation learning that supports genome comparison, functional annotation, phenotype prediction, candidate gene prioritisation, and genome-template design across diverse bacterial taxa.

## Acknowledgements

We thank Vitor Mendes, Chris Ruis, Stephane d’Ascoli, Aïcha Bentaieb, Alma Andersson, Maxwell Berk Monconis, Kristina Kordova, Charlie Harris, Oliver Hernandez, Timon Schneider, Paulina Kulyte, Chris Brown, Audra Devoto and Daniela Goltsman for helpful discussions on the study design and manuscript. We thank Severine Habert, Nikolai Solmsdorf and Anja Hagting for assistance with computational infrastructure. We thank Julia Staniszewska for help with visualisations. We thank the *SPIRE* authors for help in accessing the genomic sequences.

## Author Contributions

M.W., A.W. and R.A.F. conceived the project. R.A.F., A.W., M.B. and J.L. supervised the project. M.W. designed the model architecture. M.W., A.W. and M.N. curated and preprocessed pretraining datasets. M.W., A.W., R.V.T. and D.C.A. curated and preprocessed datasets for finetuning and downstream task evaluation. M.W. implemented models and infrastructure for pretraining. M.W. evaluated the pretraining model and conducted the finetuning. M.W., D.C.A., R.M. and A.W. conducted experiments for downstream task analysis. D.A. prepared the samples for long-read RNA sequencing. A.W., M.W. and D.A. preprocessed and analysed the long-read RNA sequencing results. R.V.T. conducted experiments on phenotypic trait prediction. D.C.A. conducted experiments on antimicrobial resistance prediction. A.D. contributed to protein–protein interaction prediction analysis. S.H. and J.L. contributed to the modelling infrastructure. M.W., R.A.F., A.W., M.B., S.H. and J.L. contributed to study design and analysis. M.W. implemented the public codebase. R.A.F., M.W. and A.W. wrote the first draft of the manuscript. All authors wrote the final draft of the manuscript.

## Code and data availability

Code and models related to this study, together with tutorials, are publicly available at https://github.com/macwiatrak/Bacformer under a permissive *Apache 2.0* license. Code for benchmarking and using different genomic LMs is available at https://github.com/macwiatrak/BacBench.

The genomic sequences and assemblies analysed in this study are publicly available from *MGnify* [27], *SPIRE* [28], and NCBI RefSeq [38]. The long-read RNA sequencing operon annotations are available at https://huggingface.co/datasets/macwiatrak/operon-identification-long-read-rna-sequencing. Downstream-task datasets used for evaluation and benchmarking are available at https://huggingface.co/datasets/macwiatrak/datasets.

Additional details on the datasets used in this study and their processing are provided in Methods.

## Supplementary Materials

### METHODS

#### Model architecture

The *Bacformer* architecture is designed to mirror how bacterial genomes are structured. It represents complete genomes consisting of different replicons, i.e. the primary chromosome alongside any chromids or plasmids, as well as fragmented isolate genomes and metagenome-assembled genomes (MAGs) distributed across multiple contigs, while explicitly capturing the relationships between neighbouring genes.

*Bacformer* utilises a pretrained protein language model (pLM) to generate embeddings for each protein present in the bacterial genome. These protein-level vectors are then passed to a genomescale transformer that contextualises every representation across the bacterial genome.

##### Input embeddings

Every input sample to *Bacformer* is an ordered set of proteins from a bacterial genome (an ordered sequence of genes on one or multiple contigs, where each contig can correspond to a chromosome, a plasmid, or an assembled contig).

*Bacformer* represents each bacterial genome as a set of protein sequences in the context of their ordered locations on the genome. To do so, each protein is treated as a token, and tokens combine information from (1) a protein embedding, (2) a contig (also referred to as chromosome or plasmid) embedding, and (3) a relative position embedding to encode relative distance between proteins, using rotary positional embeddings [1].

To create these, each genome *G* is first converted into a list of proteins *P_i_* ordered by their coordinates on the bacterial chromosome, plasmid(s) or contigs, so that *G* = (*P*_1_, *P*_2_, *…*, *P_N_*) for a genome *G* with *N* proteins.

###### Protein embeddings

A pretrained pLM maps every protein sequence *P_i_* to a matrix of residue embeddings; to get a single vector for a protein we average the residue embeddings,

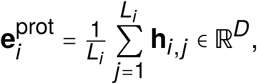

yielding a fixed-length vector of dimension *D*. We use protein language models from the *ESM* family, specifically *ESM-2* (12 layers, 35M parameters, *D* = 480) [2] and *ESM-C* (30 layers, 332M parameters, *D* = 960) [3] as the base protein language models for *Bacformer* and *Bacformer Large* respectively.

###### Contig embeddings

To encode the order of proteins and their location on the genome, we represented contigs using a *contig embedding* 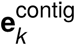 for each of the *K* contigs that are part of the bacterial genome. As each contig embedding (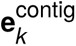) is the same for all proteins on the contig, the model can therefore distinguish, for instance, plasmid-specific gene clusters.

For a protein *P_i_* residing on contig *c_i_* we form a token vector

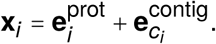

Special learnable tokens are appended to mark global structure: `[CLS]` at the beginning, `[SEP]` between contigs, and at the genome terminus. The ordered sequence

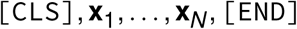

preserves native synteny, allowing the transformer to represent functional dependencies such as operons [4].

###### Rotary positional embeddings

The rotary position embeddings [1] are incorporated in the attention mechanism to encode relative distances between proteins, enabling the model to capture genomic context. This is essential in bacteria, where genes are often organised in operons—compact, co-transcribed clusters whose function and regulation depend on precise ordering and spacing. Rotary embeddings are used instead of sinusoidal schemes to avoid length limitations.

Below we show examples of ordering in genomic sequences for complete genomes and MAGs. `chr`, `plasm` and `contig` refer to chromosome, plasmid, and contig, respectively.

**Complete genome:**

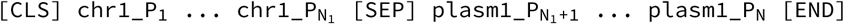

**MAG:**

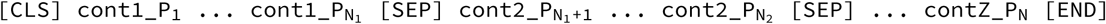

During pretraining on MAGs, we randomised the order of contigs within every bacterial genome for every epoch to prevent the model from learning spurious signals from the arbitrary contig ordering produced by the metagenomic assembly.

Finally, each token is processed by a shared linear layer, followed by layer normalisation and dropout (with a rate of 0.2). The resulting sequence of embeddings for each bacterial genome is then fed back into the transformer, which updates each protein representation by attending to all other proteins in the genome, thereby capturing their genomic context and relationships.

##### Transformer block

Each bacterial genome, consisting of *N* proteins, is fed into a transformer to learn protein embeddings conditioned on all other proteins in the genome, yielding contextualised representations. The rationale is that a genome can be viewed as an ordered set of proteins whose interactions ultimately determine the organism’s phenotype.

We employ rotary positional embeddings (RoPE) [1] within the self-attention mechanism and use `FlashAttention` [5] in `PyTorch` for memory-efficient training and inference. The transformer is trained from scratch. For downstream tasks, we use the final hidden state of each protein, while the mean of all token representations provides a fixed-length embedding of the whole genome.

##### Model sizes (Bacformer and Bacformer Large)

We trained two versions of *Bacformer*, *Bacformer* and *Bacformer Large (Bacformer-L)*, with 26M and 332M encoder parameters, respectively. The 26M *Bacformer*model consists of 12 transformer layers with 8 attention heads and a hidden dimension of 480. *Bacformer Large* (332M parameters) has 30 layers, 15 attention heads, and a hidden dimension of 960. The two architectures are the same, except that *Bacformer-L* uses the SwiGLU activation function [6] instead of GELU [7] and omits `[CLS]` and `[SEP]` tokens using contig embeddings for contig boundaries. This choice was motivated by empirical evaluation.

Throughout the text we refer to *Bacformer* as the 26M parameter model, and *Bacformer Large* as the 332M parameter model.

#### Pretraining

We sought to learn the relationships between proteins present in the genome, leveraging billions of years of bacterial evolution. All large language models (LMs), including protein LMs and DNA LMs, rely on a discrete vocabulary set for pretraining. In *Bacformer*, each token is a protein, and the universe of bacterial proteins is effectively unbounded. To overcome this, we constructed a finite protein family vocabulary.

##### Unsupervised protein family clustering for pretraining

We ran unsupervised clustering on millions of proteins across organisms using *k*-means clustering [8]. We leveraged *ESM-2* [2] and *ESM-C* [3] protein embeddings to represent the proteins and, after experimenting with various values of *k*, set it to 50, 000 centroids based on the small scale pretraining experiments where we monitored the validation loss. Pre-training strongly relies on the unsupervised clustering producing high-quality protein family classes that reflect evolutionary processes. Therefore, after clustering we performed quality checks leveraging the *ESM-2* protein embedding clustering—evaluating distances to the centroids and the number of proteins per cluster. We removed labels for proteins that exceeded a Euclidean distance threshold of 2.0 to the closest centroid. We leveraged the *CuPy* library [9] to efficiently perform clustering on millions of proteins.

To confirm that these clusters aligned with known evolutionary relationships, we used *eggNOG* 5.0 [10], a database of protein orthology across organisms. Using *eggNOG* annotations available through *MGnify* [11] on a set of 12, 630 diverse species-representative genomes, we extracted annotations for all proteins with an *eggNOG* label and compared them to the cluster labels from the *k*-means algorithm. We found strong agreement between the *eggNOG* annotations and our clusters, with the two achieving a 74% normalised mutual information score (NMI; range 0–1, higher is better), indicating that our *k*-means clusters capture independently inferred groupings.

Moreover, we compared the *k*-means clustering with *MMseqs2*, a widely used protein clustering method [12]. We randomly sampled 12.3 million proteins from species-representative genomes, clustered them with *MMseqs2* (minimum sequence identity = 0.8 and coverage = 0.8), and observed 88% NMI concordance with our *k*-means solution.

To evaluate functional coherence of *Bacformer*protein family clusters, proteins with KEGG annotations [13] were mapped to KEGG BRITE functional categories. Cluster purity was computed from the contingency table between *Bacformer* clusters and KEGG BRITE categories. For each cluster, the majority BRITE category was identified, and purity was defined as

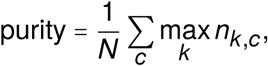

where *n_k_*_,*c*_ is the number of proteins assigned to KEGG BRITE category *k* in cluster *c*, and *N* is the total number of BRITE-annotated proteins included. Functional enrichment was assessed for each cluster–BRITE-category pair using an upper-tail hypergeometric test against the full annotated protein background, considering clusters with at least 10 annotated proteins. Resulting *p*-values were corrected using the Benjamini–Hochberg procedure, with significant enrichment defined at *q* < 0.05.

To ensure that the protein family vocabulary retained sufficient within-genome diversity for pretraining, we calculated the number of unique protein family clusters present in each genome and normalised by the total number of proteins.

The protein family label is predicted using a two-layer multilayer perceptron (MLP) that receives the output of the last hidden layer of the transformer. The MLP is trained from scratch over the course of pre-training. Importantly, we only use the protein family labels for supervision during pre-training; the input protein embeddings are constructed from amino-acid sequences, not from family labels. This design allows the model to tackle tasks that require amino-acid-level sensitivity. We also experimented with using protein family labels as token IDs learned from scratch, but amino-acid-level input yielded better performance on downstream tasks.

##### Training objective

We trained two versions of *Bacformer* with different self-supervised objectives: (1) an autoregressive next-protein family prediction (generative) objective and (2) a masked-prediction objective. In practice these approaches confer complementary strengths on downstream tasks. Each model contains transformer layers followed by a classification head that predicts the protein family label from the final layer’s output.

The masking procedure resembles the standard BERT-style scheme [14], adapted to our setting: (*i*) 15 % of the proteins are selected for masking. (*ii*) In 87.5 % of those cases the selected tokens are replaced by `[MASK]`. (*iii*) In the remaining 12.5 % the selected tokens are left unchanged.

For *Bacformer-L*, we trained only with the masked-prediction objective to prioritise computational efficiency while retaining bidirectional attention. We chose this objective because representations from the *Bacformer* model trained with a masked objective outperformed those from the autoregressive model on most downstream tasks.

##### Pretraining details

We first pretrained both models on a corpus of ∼1.3M metagenome-assembled genomes (MAGs). Training was performed on a mixture of NVIDIA A100 and Intel Ponte Vecchio (PVC) GPUs for up to 1.2M optimisation steps; running on four A100s required approximately two weeks. The effective batch size was 32. We used a linear learning-rate schedule with a 10 % warmup, ramping to a peak of 0.00015 per GPU; the rate was then scaled by the square root of the total number of GPUs. Optimisation employed Adam [15] with weight decay 0.01 and ω = (0.9, 0.999). All tensors used the `bfloat16` format. For *Bacformer-L*, we kept the same pretraining hyperparameters, except that we used a learning rate of 0.0002 per GPU and trained on NVIDIA H100 GPUs. Models were implemented in `PyTorch` [16] and trained with the `HuggingFace` `Transformers` library [17]. We did not use early stopping.

After this stage we continued pretraining on a held-out set of 48, 932 complete genomes. We separated the two stages because MAGs usually consist of multiple contigs whose order is unknown, whereas complete genomes comprise an uninterrupted chromosome (or set of replicons) with a well-defined gene order, which can be fed to the model as a single contiguous sequence. For every genome we capped the maximum number of proteins at 6, 000, a value that exceeds the protein count of more than 99% of genomes in our corpus.

All pretrained models are available for zero-shot use or finetuning at https://huggingface.co/collections/macwiatrak/bacformer. Code for pretraining is provided in our GitHub repository: https://github.com/macwiatrak/Bacformer.

##### Pretraining performance analysis

During training, we monitored the validation loss and selected the checkpoint with the lowest value. We then evaluated pretraining performance by running the model on the validation set at regular intervals, recording accuracy for the masked-prediction objective and perplexity for the autoregressive objective. In both cases performance improved steadily with additional steps, demonstrating that the models generalise to taxa not seen during training (i.e. zero-shot; ***Figure S2***).

We next examined the effect of genome completeness. MAGs are often fragmented and may lack sizeable genomic segments; therefore, the model must be robust to varying completeness levels. Pretraining results show that *Bacformer*benefits from more complete assemblies. However, it remains relatively robust until completeness drops below 75 % (***Figure S2***). The same trend appears when comparing MAGs with complete genomes (***Figure S3***): access to the full sequence and an intact chromosome without contig breaks markedly boosts accuracy.

Finally, we investigated scaling. A variant trained on only 25% of the data performed noticeably worse as measured by perplexity, indicating that *Bacformer* benefits from additional training data. Moreover, the larger *Bacformer-L* achieved substantially lower validation loss on the same data and performed much better on downstream tasks, demonstrating the benefits of scaling the model size (***Figure S3***). These results suggest that increasing both the amount of training data and the number of model parameters is likely to further improve performance.

#### Data

We collated a diverse set of bacterial genomes spanning the tree of life and a wide range of environments. Our guiding principle was to maximise phylogenetic and ecological diversity so that the pretrained model would be broadly applicable to downstream tasks in bacterial genomics. For this reason, we used MGnify and SPIRE resources, which contain large collections of diverse metagenome-assembled genomes (MAGs) from different environments.

##### MAG corpus

For pretraining we assembled the MAG corpus from two complementary resources: *MGnify* v2.0 [11] and *SPIRE* [18]. *MGnify* provides seven biome-specific genome catalogues that together contain 306, 596 non-redundant metagenome-assembled or isolate genomes representing 11, 048 species. *SPIRE* provides 99, 146 consistently processed whole-genome shotgun metagenomic samples drawn from 739 studies worldwide and 1, 008, 689 MAGs.

We downloaded the assemblies containing protein annotations and associated metadata from both resources. Protein order within each contig was preserved so that positional encodings could capture synteny. Lineage labels (species, genus, family) and contextual metadata (sample identifier, biome, etc.) were processed as provided by *MGnify* and *SPIRE*.

To ensure that our corpus retained high-quality assemblies while preserving broad taxonomic and environmental diversity, we applied a series of quality-control filters before pretraining. We kept only genomes with estimated completeness > 50% and removed highly contaminated genomes (contamination > 10%), thereby excluding assemblies likely to contain large missing regions or substantial signal from other organisms. We also removed genomes with abnormal genome lengths (< 250 kb or > 20 Mb), which may indicate severely incomplete assemblies, mis-binned contigs, or non-bacterial material, as well as highly fragmented genomes (> 1, 000 contigs), where long-range gene order is unlikely to be reliable. These filters were chosen to reduce technical artefacts while retaining the diversity of MAGs needed for robust genome-context pretraining.

The final MAG corpus after quality checks comprises 1, 314, 344 genomes, totalling 2, 700, 709, 306 proteins, 220, 207, 650 contigs, 24, 663 species and 70 distinct environments.

##### Complete-genome corpus

Many downstream applications such as comparative genomics, gene-context analysis and operon prediction benefit from access to complete isolate genomes. Unlike MAGs, which are reconstructed *in silico* from a metagenome-wide assembly and are often fragmented, incomplete or contaminated, complete genomes are obtained from clonal cultures and typically consist of a single circular chromosome (and any additional chromids or plasmids) with high continuity and accuracy [19]. To capture these characteristics, we downloaded available bacterial assemblies from NCBI RefSeq [20] and GenBank [21]. After removing duplicates, we retained 48,932 unique complete genomes, each with its translated proteome and annotation files. The metadata (i.e., assembly accessions) were stored alongside the sequences. The NCBI collection is biased towards clinically important pathogens—for example, *Salmonella enterica*, *Staphylococcus aureus* and *Mycobacterium tuberculosis*. The combined corpus, consisting of the MAG and complete-genome corpora, therefore encompasses a heterogeneous yet complementary training set that balances ecological breadth with assembly quality, positioning the model to generalise across both environmental and clinical bacterial genomics despite the currently available datasets being biased and broad sections of the global bacterial ecosystem being under-represented.

##### Embedding genomes with a protein language model

*Bacformer* represents each genome by the average of its protein embeddings computed with a pretrained protein language model (pLM). To avoid the cost of re-embedding every protein at every training epoch, we generated all embeddings once, off-line, and cached the resulting protein embeddings.

We chose the 12-layer, 35M-parameter variant of *ESM-2* [2] for *Bacformer* and 30-layer 332M-parameter variant of *ESM-C* [3] for *Bacformer-L* because it offers a good balance between accuracy and computational efficiency when compared with larger pLMs such as *ESM-2-650*M or *ProtT5-XL* [2, 22]. Embedding the entire corpus of ∼1.3 million genomes with ESM-2 (35M) required approximately 30 days on a cluster equipped with 16 Intel Ponte Vecchio GPUs, and produced ∼5.6 TB of embedding files.

##### Long-read RNA sequencing for operon identification

To test *Bacformer*’s ability to predict operons, we performed long-read RNA sequencing to annotate operons across five distinct strains, processing three independent biological replicates for each strain.

###### Bacterial strains and culture conditions

Five bacterial strains were used in this experiment: *Staphylococcus aureus RN450* (*S. aureus* RN450), *Mycobacterium abscessus* ATCC 19977 (*M. ab*), εleuD εpanCD *Mycobacterium tuberculosis* H37Rv 102J23 (εleuD εpanCD *M. tb*), *Pseudomonas aeruginosa* PAO1, and *Escherichia coli DH5*ϑ. Each strain was cultured in triplicate under nutrient-rich conditions until mid-exponential phase (*OD*_600_ = 0.4–0.6) was reached.

###### RNA isolation

Cells were harvested by centrifugation and processed for total RNA extraction using the MasterPure Complete DNA and RNA Purification Kit (Lucigen) with strain-specific modifications: For *S. aureus* RN450, cell pellets were pre-treated with lysostaphin (Tris buffer, pH 8.0) at 37 °C for 30 min to aid lysis. For the mycobacterial strains (*M. ab* and εleuD εpanCD *M. tb*), cells were mechanically disrupted by bead beating in lysis buffer, followed by extraction with the standard kit protocol. Isolated RNA was treated twice with TURBO DNase (Invitrogen) to remove residual genomic DNA and purified using RNA Clean & Concentrator columns (Zymo Research). RNA integrity was assessed on an Agilent TapeStation RNA ScreenTape system, and concentrations were measured with a Qubit fluorometer (Invitrogen).

###### Library preparation and sequencing

For each replicate, 1,000 ng of total RNA underwent rRNA depletion with riboPOOLs (siTOOLs Biotech). The depleted RNA was polyadenylated with poly(A) polymerase (PAP) in the presence of a manganese catalyst, adding 50–90 adenosines per molecule. cDNA libraries were prepared with the Nanopore cDNA-PCR kit, pooled and sequenced on a PromethION device equipped with R10 flow cells (Oxford Nanopore Technologies).

#### Downstream tasks & analysis

##### Benchmarking

To evaluate *Bacformer* against other methods, we performed rigorous benchmarking comparing *Bacformer* to other deep learning methods. We selected a diverse set of genomic LMs, including DNA language models (DNA LMs), protein language models (pLMs) and a mixed-modality genomic LM. These genomic LMs take as input DNA, proteins, or both (*gLM2* [23]) and can therefore generalise across the bacterial tree of life. Moreover, the suite spans models which leverage distinct modalities: DNA-only, single-protein, mixed DNA–protein, and multiple proteins present in the genome (i.e. Bacformer), objectives (masked vs. autoregressive), and training corpora (bacteria-specific vs. cross-kingdom), enabling a controlled comparison of how context length, modality, and pretraining data shape bacterial genome embeddings. The details of the benchmarked methods can be found in *Table S9*.

Below we outline how we represent and embed bacterial genomes using DNA LMs, pLMs, mixed-modality LMs (*gLM2*) and *Bacformer* models. We differentiate between task types that require different representations from the models; specifically: (*i*) gene- and system-level tasks, where the prediction target is a gene, protein, protein pair or local gene neighbourhood, including protein–protein interaction prediction, operon identification, gene essentiality and protein function prediction. In these tasks, we extract task-specific representations for the relevant gene(s) or protein(s), optionally retaining local genomic context when supported by the model; (*ii*) genome-level prediction, such as genome clustering, phenotypic traits prediction and antimicrobial resistance prediction (AMR), where the prediction target is the whole genome and representations are obtained by aggregating gene-, protein-, window- or contig-level embeddings into a single genome vector. We describe the setup that yielded the best representations for the downstream tasks across gene-, system- and genome-scale and different model types. All of the modelling code was implemented in `PyTorch` [16]. The benchmarking code is available at https://github.com/macwiatrak/BacBench.

###### DNA Language Models (DNA LMs)

For every model, we loaded the public checkpoint (*Table S9*) and kept the hidden states of the *last* encoder layer, which performed best across the benchmarked DNA LMs. If a DNA string has length of *G* tokens, the encoder produces **X** ∈ ℝ*^G^*^×*D*^; we collapse it with a mean-pool to obtain a vector

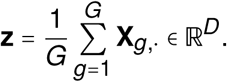

We also experimented with using max-pooling which yielded worse results across tasks.

###### Gene- & system-level

When the input gene (plus upstream promoter) is longer than the model context *C*, we slide a window of length *C* with stride *s* and embed each window. Averaging the *M* window vectors {**z**^(1)^, *…*, **z**^(^*^M^*^)^} gives the final gene representation 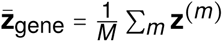.

For Evo and Evo-2, the context length often exceeds the length of the target gene. In this setting, we centre the input on the gene and add approximately equal left and right flanking context. We also define a binary mask *m* ∈ {0, 1}*^L^*, where *m_i_* = 1 if token *i* belongs to the gene and *m_i_* = 0 otherwise. Given token embeddings **X** ∈ ℝ*^L^*^×*D*^, we compute the final gene representation by mean-pooling only over masked gene tokens, i.e. 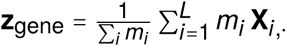. This allows the model to use flanking genomic context while ensuring that the final representation remains specific to the gene itself. Empirically, this performed better than using the same context without the gene mask.

###### Genome-level

We treat the whole genome in the same way: split it into *C*-bp windows, embed each window, and average; if several contigs are present, we first average per contig and then across contigs.

###### Protein Language Models (pLMs)

A protein of *K* amino acids yields **X** ∈ ℝ*^K^*^×*D*^ and 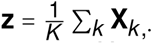. For genome-level tasks we aggregate the *M* protein vectors in that genome via 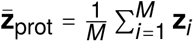. Gene- and operon-level tasks use only the proteins involved. For all models we use the representations from the last hidden layer and mean pooling.

###### *gLM2* (mixed-modality LM)

*gLM2* ingests mixed-modality genomic scaffolds in which protein-coding regions are translated to amino-acid tokens and intergenic regions remain as nucleotide tokens. We tokenize each scaffold into a single sequence up to the model context *C* (4,096 tokens); longer scaffolds are processed with a sliding window of length *C* and stride *s*, mirroring the DNA LM setup. For each window, we retain the last-layer hidden states and mean-pool to obtain a window vector **z**^(^*^m^*^)^ ∈ ℝ*^D^*; genome-level embeddings average across all windows for a genome (and across contigs when present), where *M* denotes the total number of such windows:

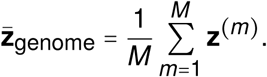

For gene essentiality and operon tasks, inputs include contextual flanking sequence to supply regulatory and positional cues, but the final gene representation is computed using the same masking scheme as above. Specifically, for a window of length *L* with hidden states **X** ∈ ℝ*^L^*^×*D*^, we define a binary mask *m* ∈ {0, 1}*^L^*, where *m_i_* = 1 if token *i* falls within the gene coordinates and *m_i_* = 0 otherwise, and compute 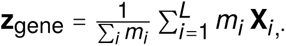,⋅. Empirically, this performed better than averaging over the entire contextual slice.

###### Bacformer

*Bacformer* models take as input *local* protein vectors **z**_1_, *…*, **z***_M_* obtained exactly as in the pLM setting. These protein vectors are computed using existing pLMs, specifically ESM-2 for Bacformer and ESM-C for Bacformer Large. Protein vectors are then **ordered by their genomic coordinates** (chromosome followed by plasmids) so that the model can incorporate genome organisation. Rotary positional embeddings [1] are added to these vectors and the ordered sequence is fed to a *genome-level* Transformer encoder [24] with *L* layers,

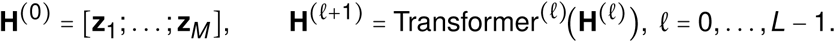

The output **H**^(^*^L^*^)^ = [**z̃**_1_; *…* ; **z̃***_M_*] contains **contextualised protein embeddings z̃***_i_* ∈ ℝ*^D^* that encode both the protein sequence and its genomic neighbourhood. During pretraining, Bacformer learns to predict which proteins co-evolved, so these embeddings capture functional coupling across the genome.

Empirically, averaging token embeddings performed slightly better than using the special [CLS] token, so mean pooling was used for every model throughout the paper.

Code for embedding genomes using the benchmarked methods can be found at https://github.com/macwiatrak/BacBench. We include details on the context length of each model (*C*) and stride used (*S*) in *Table S10*.

###### General modelling details

For all tasks and settings that required finetuning, including gene essentiality prediction and the finetuned PPI setup, we kept the backbone encoder frozen and trained only the task-specific neural network layer(s) stacked on top of the pretrained embeddings, unless explicitly stated otherwise that the encoder was finetuned. This was motivated by the computational cost required to embed all 51k genomes used for benchmarking with all models. Further details on runtime and computational cost can be found below in *Computational complexity analysis*. We used the Adam optimiser [15] in all finetuning setups. All checkpoints used were downloaded directly from HuggingFace and are specified in *Table S9*. For each task and model combination, we tuned the learning rate while keeping other parameters unchanged. Further details on hyperparameters used for each task and setup can be found in the task-specific sections below.

##### Computational complexity analysis

A typical bacterial genome contains on the order of 3, 000⌐5, 000 genes and ∼ 4⌐6 Mbp of DNA, with average protein lengths of ∼ 300 amino acids; embedding an entire genome therefore entails thousands of forward passes for protein LMs and millions of tokens for DNA LMs, making raw throughput a practical bottleneck for population-scale studies. We therefore quantified the total GPU-hours required to embed the genomic data needed for each task (*Table S11*). These estimates reveal large differences across both model families and task scales. Gene- and system-level tasks are comparatively inexpensive for most models, typically requiring well below 1 GPU-hour per task, whereas the genome-level phenotypic-trait and antibiotic-resistance tasks dominate the total cost because they require embedding of whole genomes at scale. Even relatively efficient models such as ESM-2 and Bacformer require roughly 40⌐45 GPU-hours per genome-level task, while larger DNA models such as Evo-2 require 365⌐393 GPU-hours and Evo requires 1, 426⌐1, 534 GPU-hours for the same settings. Summed across BacBench, the total embedding cost ranges from 71 GPU-hours for Mistral-DNA and 83–87 GPU-hours for ESM-2 and Bacformer to 786 GPU-hours for Evo-2 and more than 3,087 GPU-hours for Evo. These measurements show that runtime is a major practical constraint on benchmark design and model selection, especially for whole-genome evaluations at population scale.

We include the additional cost of running each experiment beyond embedding the genomes in the task-specific sections below.

###### GPU Memory Scaling for Full-Genome Finetuning

Beyond embedding costs, whole-model finetuning on genome-level tasks requires backpropagating through *all* proteins per genome (median ∼2,506 proteins across our datasets) or through multi-megabase DNA contexts, which dramatically amplifies memory and compute. ***Figure 1C*** makes this concrete by showing the peak training GPU memory for a single full-genome batch (*B* = 1) across three representative bacterial strains: (1) *Mycoplasmoides genitalium*, (2) *Escherichia coli*, and (3) *Streptomyces coelicolor*, chosen to span genomes of markedly different lengths across a representative set of genomic language models. For each model and strain, peak memory is computed as the sum of model-state memory and saved activation memory, with the number of trainable parameters obtained from the released model weights and model architecture.

To calculate the peak training GPU memory, we assume mixed precision with bfloat16 for weights and gradients and FP32 for master weights and Adam moments. Concretely, the code allocates 2 bytes/param for weights, 2 bytes/param for gradients, 4 bytes/param for FP32 master weights, and 8 bytes/param for Adam’s first/second moments, yielding 16 B/param in total. We assume FlashAttention [5] for all architectures, so the *T* × *T* attention matrices are never materialised. For a sequence length of *T* and a Transformer with hidden size *D*, feed-forward *F*, and layers *L*, the saved activations per saved layer are linear in *T* : residual states (*T* · *D*), *Q*/*K*/*V* projections (3 × *T* · *D*), and MLP activations (*T* · *F*). Total activation memory is the sum of these terms multiplied by the number of saved layers and by the element byte-width; we store activations in bfloat16 (2 bytes/element). For Evo, which replaces attention with Hyena operators, there are no *Q*/*K*/*V* projections and we only calculate the residual (*T* · *D*) and MLP (*T* · *F*) terms, preserving the same linear-in-*T* scaling. Even under this assumption, the absolute memory requirements remain enormous for large long-context models: Evo already requires hundreds of GB for *M. genitalium* and rises above 10^4^ GB for *S. coelicolor* in ***Figure 1C***, far beyond the memory of a single H100 GPU. By contrast, Bacformer-style bacterial LMs remain in the single-digit-GB range across all three genomes, helping explain why full-genome finetuning is only practically accessible for this model family and why linear-probe evaluations are a necessary, controlled proxy for model selection at scale.

##### Strain clustering

###### Benchmarking

We compared *Bacformer* to other protein-based methods and the mixed-modality model *gLM2* [23]. We assembled a randomly sampled dataset of 6, 071 MAGs from *MGnify* [11], spanning 25 species across 10 genera and 7 families, selected to provide a balanced phylogenetic spread. For every model, we took the final-layer hidden states and mean-pooled across the entire genome to yield one fixed-length embedding per strain.

Additionally, we benchmarked *Bacformer* against traditional genome clustering methods, average amino-acid identity (AAI) [25] and average nucleotide identity (ANI) [26]. To cluster 6, 071 MAGs using AAI, we extracted all predicted protein sequences for each MAG and performed an all-vs-all protein search with MMseqs2. Hits were filtered by e-value (≤ 10^−5^), query and target coverage (≥ 50%), and minimum sequence identity (≥ 50%). For each genome pair, AAI was computed as the mean percent identity across reciprocal best protein hits, retaining pairs with reciprocal hits covering at least 20% of the smaller proteome. The resulting AAI matrix was converted to a distance matrix and clustered with Leiden clustering over a grid of nearest-neighbour and resolution parameters.

We also attempted to run ANI; however, it failed to produce meaningful similarities for many genome pairs because ANI requires sufficient nucleotide-level homology between genomes and is primarily reliable for same-species or closely related genomes, typically in the ∼80–100% ANI range.

To compute strain-clustering metrics using the representations from the deep learning methods or the distance matrix from AAI, we ran Leiden clustering over a parameter grid, varying the resolution in `[0.1, 0.25, 0.5, 1.0]` and the number of neighbours in [5, 10, 15], and evaluated all pairwise combinations. We then retained the combination of resolution and number of neighbours that maximised the mean of the three metrics described above (ARI, NMI and ASW) at the species level. Full results are available in *Table S12*. We used Scanpy [27] to perform the clustering.

To examine whether performance changed on genomes that were not seen during training (i.e. zero-shot), we sampled a set of 2, 612 genomes of taxa not seen during training from the MAG corpus validation set, containing 15 distinct species from 10 distinct families. We then embedded these genomes using *Bacformer-L* and followed the same evaluation procedure as described above.

Code for strain-clustering benchmarking is available at https://github.com/macwiatrak/BacBench/tree/main/bacbench/tasks/strain_clustering, and the strain-clustering datasets used for benchmarking are available via HuggingFace at https://huggingface.co/datasets/macwiatrak/bacbench-strain-clustering-dna-small (DNA) and https://huggingface.co/datasets/macwiatrak/bacbench-strain-clustering-protein-sequences-small (protein sequences).

To test whether *Bacformer* preserves intra-family similarity, we computed average species embeddings for the most common bacterial order in *MGnify* [11], *Bacteroidales* (*N* = 64, 713), and calculated cosine similarities between the 100 most prevalent species. Species embeddings were obtained by averaging the embeddings of all strains belonging to that species. *Bacformer* retained high similarity within taxonomic groups, achieving an average cosine similarity of 0.96 ± 0.02 for species in the same genus and 0.93 ± 0.05 for species in the same family, versus 0.81 ± 0.03 and 0.80 ± 0.03 for species from different genera and families, respectively. These results show that *Bacformer* can robustly cluster species while maintaining phylogenetic relationships in an unsupervised setting. All strain-clustering experiments used the model pretrained on MAGs with the causal (autoregressive) objective.

###### Revealing sub-populations in *Escherichia coli*

To evaluate whether *Bacformer* can discover sub-populations within a single species, we clustered embeddings of *E. coli* strains. Average genome embeddings from *Bacformer-L* for 7, 585 *E. coli* strains from *MGnify* were clustered with *Leiden* clustering, yielding six well-defined groups (***Figure 1G***). Repeating the analysis with *ESM-C* embeddings failed to resolve distinct clusters (***Figure S5***) on a UMAP plot. To ensure that the clusters produced by *Bacformer-L* align with existing methods and to quantify the difference between *Bacformer-L* and its base model *ESM-C*, we ran ANI [26] and Panaroo [28] on a subset of 2, 479 *E. coli* genomes. ANI values were converted to distances (1 ⌐ ANI/100), missing ANI values were treated as maximal distance, and a 15-nearest-neighbour graph was clustered with *Leiden* clustering. Panaroo gene presence/absence profiles were reduced to 50 principal components, converted to a 15-nearest-neighbour graph, and clustered with *Leiden*. For each method, we ran *Leiden* over the same resolution grid, selected the resolution with the highest value, and quantified agreement between model-derived clusters and ANI/Panaroo clusters using normalised mutual information and adjusted Rand index. To obtain uncertainty estimates, we performed bootstrapping from 100 random seeds using 80% genome subsamples.

To exclude the possibility that the observed structure arose from technical batch effects, we asked whether strains grouped by sample or study identifier (***Figure S5***). The ASW for both metadata labels was negative, indicating no such grouping and confirming that *Bacformer* captures genuine biological structure.

Having identified the sub-populations, we sought the genes specific to each cluster. Using *Panaroo* [28], which provides a binary gene-presence matrix, we performed Fisher’s exact tests [29] comparing each cluster against all others. Genes significantly enriched in a cluster (*p* < 0.05) were considered differentially abundant; the top five per cluster are shown in (***Figure S5***).

Variation within *E. coli* is often driven by the accessory genome [30]—genes present in only a subset of strains. Defining rare genome as all genes present in < 5 % of all strains, we found that 41% of cluster-specific genes were rare. Given that the median genome of *Escherichia coli* contains 112 rare genes versus 3, 718 core genes (those present in> 95% of all strains), this represents a 23-fold enrichment (*p* < 10^−20^), highlighting the role of the accessory genome in shaping sub-populations and the utility of *Bacformer* for their discovery.

###### Scalable search for similar bacteria

Leveraging its learned genome representations, We used *Bacformer* trained with masked objective to embed all 1, 314, 344 genomes in the MAG corpus together with metadata into a unified database. Users can submit an unannotated bacterial genome, retrieve its nearest neighbours and obtain contextual metadata such as environmental origin.

We validated this approach with a *k*-nearest-neighbour (kNN) classifier. Using *k* = 10, we randomly held out 10% of genomes, filtering out species represented by fewer than five genomes. A *k*NN model trained on the remaining embeddings achieved balanced accuracies of 0.83±0.32, 0.94±0.17 and 0.95 ± 0.16 at the species, genus and family levels respectively, thus demonstrating robust performance across taxonomic ranks. This validated functionality is openly available on GitHub (https://github.com/macwiatrak/Bacformer), enabling researchers worldwide to analyse unannotated bacterial genomes with unprecedented scale and accuracy.

##### Protein-protein interaction prediction

###### Protein–protein interaction prediction using *STRING DB*

We downloaded the complete data archive from the *STRING DB* (v12.0) [31] download site (https://string-db.org/cgi/download). Using the species-metadata file, we selected only bacterial organisms and retrieved their protein sequences together with all available protein–protein interaction scores. After running the download scripts, we obtained 10, 533 unique strains spanning 6, 956 species. *STRING DB* provides several evidence-channel scores; here we used the *combined*, *co-occurrence*, *neighbourhood* and *co-expression* channels.

To binarise the interaction scores, we applied a threshold of 0.6 (≥ 0.6 = interaction, < 0.6 = no interaction), resulting in ∼10% of all interactions having a positive label. This cut-off was selected empirically by maximising the average AUROC and AUPRC on a validation set. The data were randomly split into training, validation and test sets in a 70/10/20% ratio. For each strain, we retained up to 2,000,000 protein-pair labels due to memory constraints, corresponding to more than 99% of all available pairs.

For zero-shot *Bacformer*predictions and evaluation, we computed pairwise cosine-similarity scores between contextualised protein embeddings for every strain in the test set. *Bacformer* embeddings were taken from the last hidden layer of the transformer, pretrained on complete genomes with a masked objective. We benchmarked *Bacformer* predictions for interactions against *ESM-2* [2], whose non-contextual embeddings serve as input to the *Bacformer* transformer (*Table S13*). Metrics were calculated per interaction type across genomes.

After confirming that *Bacformer* captures protein–protein interactions in a zero-shot setting, we trained a linear layer using the contextual representations from the last layer of the *Bacformer* encoder. The linear layer takes as input the mean of the two contextual protein embeddings and is preceded by a dropout layer (rate = 0.2). The resulting vector for each protein pair is passed through a final binary-classification layer, again preceded by dropout 0.2. Training minimises binary cross-entropy loss; we trained for at most 10 epochs with no early stopping, clipped gradients at 2.0, and used Adam [15] with weight decay 0.01, monitoring validation loss across genomes. Hyperparameters were tuned on the validation set. The finetuned model was initialised from the *Bacformer* weights pretrained on complete genomes.

We visualised AUROC and AUPRC curves for representative strains from distinct species in the test set using the predicted interaction scores from the finetuned *Bacformer* model.

We make the *Bacformer* model finetuned for PPI prediction on the *STRING DB* dataset available at https://huggingface.co/macwiatrak/bacformer-for-ppi-prediction. The preprocessed dataset is available at https://huggingface.co/datasets/macwiatrak/bacbench-ppi-stringdb-protein-sequences.

###### Benchmarking

For benchmarking, we evaluated the quality of the representations from each model on the protein–protein interaction task. Using the species metadata file, we selected only bacterial organisms and randomly sampled 261 genomes from STRING DB across 215 genera. Random sampling was performed to balance phylogenetic breadth with computational tractability, since each strain contributes a very large number of scored protein pairs. We then downloaded the protein sequences for these strains together with protein–protein interaction scores for protein pairs. We used the *combined* interaction score, which combines information from various sources to obtain a final score. STRING DB provides only protein sequences and no DNA; therefore, to provide DNA, we matched the strain names to genomes available in GenBank and ensured that the protein sequences provided by STRING DB and GenBank matched. We then used GenBank to provide DNA for the 261 strains. To binarise the interaction scores, we set the threshold at 0.6 (≥ 0.6 = interaction, < 0.6 = no interaction). This threshold was chosen by conducting small-scale experiments and evaluating the average AUROC and AUPRC on the validation set across genomes, selecting the threshold that attained the best average performance across the two metrics.

We used a phylogeny-aware split at the genus level, assigning entire genera to the training, validation, and test sets in a 60/20/20% ratio. This prevents closely related genomes from the same genus from appearing in different splits.

For the **zero-shot** setting, given the embeddings of two proteins, we computed the cosine similarity between the two and used the binary labels from STRING DB to compute the metrics. We report the results on the test set.

In the **finetuning** setting, we trained a linear classifier on binary labels (1 = interaction, 0 = no interaction) to predict whether two proteins interact. Each gene/protein is represented by its frozen embedding. The pair of protein embeddings is averaged and processed by a single-layer neural network preceded by layer normalisation and dropout (rate 0.2). In addition to the average of the protein embeddings, we also included the cosine similarity between the two embeddings, which empirically improved performance on the validation set. Therefore, the input to the linear layer is *D* + 1, where *D* is the model dimensionality and 1 is the cosine similarity value. The model was optimised with binary cross-entropy loss. We ran a learning-rate sweep for each model on the validation set. Training proceeded for up to 20 epochs with early-stopping patience of 5 based on AUROC. We used the Adam optimiser [15] with weight decay 0.01. We report results on the test set.

For zero-shot evaluation, the experiment took approximately 90 minutes on a single NVIDIA H100 Tensor Core GPU (96 GB GPU memory) with 72 CPU cores (Grace CPU) for all models. For finetuning, it took 91 hours on the same hardware, including the learning-rate sweep. This time does not include embedding the genomes for each model; for this, see the section on *Computational complexity analysis*.

The datasets used for benchmarking are available at https://huggingface.co/datasets/macwiatrak/bacbench-ppi-stringdb-dna-small (DNA) and https://huggingface.co/datasets/macwiatrak/bacbench-ppi-stringdb-protein-sequences-small (protein sequences). The code for zero-shot and finetuning evaluation is available at https://github.com/macwiatrak/BacBench/tree/main/bacbench/tasks/ppi. Full results are available in *Table S13*.

###### Genomic distance-aware sampling for bacterial PPI

To assess whether protein–protein interaction (PPI) prediction was influenced by gene-proximity bias (***Figure S8A***), we finetuned and evaluated the *Bacformer Large* encoder and PPI prediction head on the same 261-strain benchmark dataset described above. The dataset was split at the genus level to reduce phylogenetic leakage. Each genome was represented using per-protein *ESM-C* embeddings of dimension 960, and for each strain we retained up to 2,000,000 labelled protein pairs, corresponding to more than 99% of all available pairs.

We jointly finetuned the pretrained masked *Bacformer Large* encoder and a randomly initialised PPI head. For each retained protein, *Bacformer Large* produced a contextual representation. For every labelled protein pair, the prediction head combined the two contextual protein embeddings with their cosine similarity to produce a single interaction logit. The model was trained using binary cross-entropy loss with the AdamW optimiser, a learning rate of 3 ×10^−4^, weight decay of 0.01, batch size of 1, gradient clipping at 2.0, and early stopping with a patience of 3 epochs, for a maximum of 10 epochs.

We compared two training strategies while keeping the model architecture, loss function and held-out evaluation unchanged. In the control strategy, training pairs from each genome were sampled randomly without replacement. This preserved the natural imbalance between positive and negative labels, as well as the natural distribution of short- and long-distance protein pairs (***Figure S8A***).

In the distance-aware strategy, we used distance- and label-balanced sampling. Protein-pair distances were defined as the absolute difference between within-contig protein indices and assigned to four bins: 1–5, 6–20, 21–100 and > 100. These distance bins were combined with the binary interaction label (0 = no interaction, 1 = interaction) to define eight distance-by-label groups. During training, pairs were sampled from these groups with equal probability and with replacement. This reduced the extent to which the model could rely on proximity-associated signal alone, while preserving the same architecture and objective.

For validation and testing, all retained pairs from each held-out genome were scored. AUROC and AUPRC were computed for each genome both overall and within each distance bin, and summarised across genomes (***Figure 2D***). To avoid favouring checkpoints dominated by the largest or easiest distance bins, we used the geometric mean of bin-level AUROC and AUPRC values as the monitoring metric. This analysis quantified the contribution of genomic proximity to PPI prediction performance, while the standard training strategy was retained for downstream prediction to preserve performance on the full empirical data distribution.

###### Identifying protein–protein interactions in *Pseudomonas aeruginosa*

To systematically identify potential protein–protein interactions in *Pseudomonas aeruginosa* PAO1, we applied the finetuned *Bacformer-L* interaction prediction model trained on the same dataset as used for benchmarking on the PPI task. Using finetuned model (encoder + head), we computed interaction probabilities for all possible protein pairs and retained only pairs with a predicted probability of at least 0.5.

The top 150-scoring protein pairs were then evaluated for direct physical interactions using *AlphaFold3* [32]. For each pair, we extracted the inter-protein *TM*-score (*ipTM*), which quantifies confidence in the predicted protein complex structure; higher *ipTM* values indicate a greater likelihood of a true physical interaction.

To test whether *Bacformer* prioritised protein pairs with higher *AlphaFold3 ipTM* scores, we sampled a random control set of 150 protein pairs matched by genomic distance. This matching was necessary because bacterial protein–protein interactions are biased towards neighbouring genes (***Figure S7***). We evaluated the control pairs with *AlphaFold3* and compared their *ipTM* scores with those of the *Bacformer*-selected pairs. Significance was assessed using a one-sided, distance-stratified permutation test on the difference in mean *ipTM*. Protein pairs were grouped by absolute genomic index distance, and *Bacformer-L*/control labels were permuted only within each distance stratum to preserve the distance distribution. The empirical *p* value was calculated from 20,000 permutations as the fraction of permuted mean differences greater than or equal to the observed difference. We reported the effect size both as the raw mean *ipTM* difference between *Bacformer* and control pairs and as Hedges’ *g*, calculated from the pooled standard deviation with small-sample correction.

These results demonstrate that *Bacformer* can generate useful hypotheses about gene–gene interactions and gene networks, providing a guide for experimental validation. This is particularly notable because the model is trained using *STRING DB*, which integrates diverse evidence sources and may include interactions without high-quality experimental support, leading to some false positives. *AlphaFold3*’s state-of-the-art structural predictions provide an additional screening step for prioritising these candidate interactions.

###### Attention-weight analysis

To analyse the genomic relationships captured directly by the transformer, we examined the attention weights of a *Bacformer* model trained with a masked-prediction objective on complete genomes. We collected attention scores for protein pairs from 45 diverse organisms randomly sampled from *STRING DB* [31], together with their corresponding STRING DB interaction scores. To determine which biological signals were reflected in the attention patterns, we evaluated attention weights against four STRING DB interaction categories: combined score, co-occurrence, neighbourhood, and co-expression. We compared attention weights from individual layers and the mean across all layers, and found that final-layer attention weights, averaged across attention heads, gave the most informative signal. This analysis was performed in a zero-shot setting, without any additional training.

###### Zero-shot operon prediction using long-read RNA sequencing

We mapped the reads from each replicate using minimap2 [33] and SAMtools [34] to genes present in the genome assemblies. We then merged the predictions from the triplicates for each strain and used these as our operon annotations. For each sequenced strain, the following genome assemblies and gene annotations from NCBI RefSeq [35] were used:

- `GCF_000013425.1` (*Staphylococcus aureus RN450*),
- `GCF_000195955.2` (εleuD εpanCD *Mycobacterium tuberculosis H37Rv 102J23*),
- `GCF_000069185.1` (*Mycobacterium abscessus ATCC 19977*),
- `GCF_000006765.1` (*Pseudomonas aeruginosa PAO1*),
- `GCF_002899475.1` (*Escherichia coli DH5*ϑ).

To predict operons, we posed operon prediction as a binary boundary prediction problem, evaluating whether two contiguous genes form an operon or not. We established a method that uses: (1) contextualised protein embeddings from *Bacformer*, (2) the distance between the two genes (intergenic region in base pairs), and (3) the strand of each gene. We then constructed a model that takes these features as input and outputs a binary value indicating whether the gene pair forms an operon, based on:

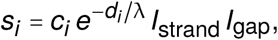

where

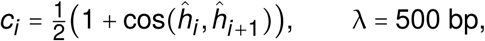

and

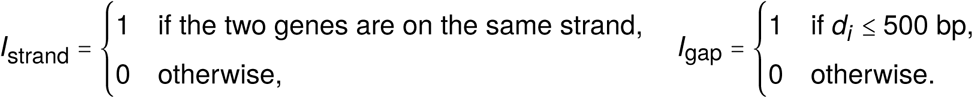

Here *ĥ_i_* and *ĥ_i_*_+1_ are the ϖ_2_-normalised *Bacformer* embeddings of genes *i* and *i*+1, *d_i_* is their intergenic distance, and *s_i_* ∈ [0, 1] serves as the operon-membership score; a pair is classified as belonging to the same operon when *s_i_* exceeds a threshold. The exponential factor 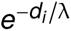 reduces the score of pairs separated by larger intergenic distances, while the two indicator terms *I*_strand_ and *I*_gap_ act as a hard veto, nullifying the score if genes are transcribed in opposite directions or if the gap exceeds 500 bp. This reflects the empirical observation that bacterial operons almost always consist of closely spaced, co-oriented genes.

We evaluated operon identification in a zero-shot manner; therefore, we did not split the data into training, validation and test sets, and instead used the entire dataset as a test set.

The experiment was run on a single NVIDIA H100 Tensor Core GPU (96 GB GPU memory) with 72 CPU cores (Grace CPU) and took < 5 minutes for all models. This time does not include embedding the genomes for each model; for this, see the section on *Computational complexity analysis*.

###### Benchmarking

We performed benchmarking by utilising proprietary long-read RNA sequencing results. The evaluation was performed in a zero-shot manner.

To focus the evaluation on the quality of the embeddings, we did not use intergenic distances and instead used the cosine similarity between contiguous gene/protein embeddings as well as strand information to evaluate performance. Specifically, we posed operon prediction as a binary boundary prediction problem, evaluating whether two contiguous genes form an operon or not. Given *c_i_* as the cosine similarity between two contiguous genes and *I*_strand_ as a binary value indicating whether the two genes are on the same strand, the score for the two genes belonging to the same operon is calculated as *s_i_* = *c_i_* × *I*_strand_. We report performance averaged across all five sequenced strains on the area under the receiver operating characteristic (AUROC) and precision-recall (AUPRC) curves.

The preprocessed datasets for operon identification are available at https://huggingface.co/collections/macwiatrak/bacbench. The code used for benchmarking is available at https://github.com/macwiatrak/BacBench/tree/main/bacbench/tasks/operon. Full operon-identification results and paired statistical comparisons are available in *Table S1* and *Table S2*, respectively.

##### Uncovering gene networks from contextual embeddings

To test whether *Bacformer*’s contextualised representations predict gene co-expression and co-regulated genes, we compared zero-shot clusters obtained from pairwise cosine similarities between *Bacformer* embeddings with those from experimental co-expression data. Normalised RNA-seq data from 411 samples of *P. aeruginosa* were obtained from iModulonDB [36, 37] and used to compute a gene–gene co-expression matrix. Equivalent matrices were produced from *Bacformer* and from *ESM-2* embeddings to benchmark *Bacformer* results. Each matrix served as input to the Leiden clustering algorithm [38] across multiple resolutions. Cluster similarity between co-expression and model-predicted partitions was quantified using normalised mutual information (NMI, range 0–1; ***Figure S7***). For qualitative inspection, we extracted clusters at resolution 40 containing key regulators such as *lasR* (***Figure 2H***), *fleQ*, and *algU* (***Figure S7***).

##### Protein function annotation

###### Gene essentiality prediction and benchmarking

We obtained bacterial gene essentiality annotations from the Database of Essential Genes (DEG v15, http://origin.tubic.org/deg) [39], including metadata, annotations, and the amino-acid and nucleotide sequences provided by DEG. Because DEG is organised at the gene level rather than the genome level, we used the RefSeq genome identifiers in DEG to retrieve the corresponding assemblies from NCBI GenBank (https://www.ncbi.nlm.nih.gov/genbank/) in both DNA and protein sequence modalities. We then mapped DEG essential genes back to genes in the downloaded genomes using a multi-step matching procedure based on exact protein-sequence matches, locus tags, UniProt IDs, protein IDs, gene names, and length-based matches with more than 90% identity. This strategy maximised correct assignments and resolved ambiguous cases using locus tags or other identifiers when available. Of the 66 genomes present in DEG, we removed duplicate genomes, leaving 55 genomes in the final dataset. For each genome, we retained protein and DNA sequences together with gene-level metadata including strand, start, end, and related annotations. The final dataset contains 193,213 proteins, of which 21,626 are labelled as essential and 171,587 are non-essential.

We annotated each genome with its genus and used a phylogeny-aware split at the genus level, assigning entire genera to the training, validation, and test sets in a 60/20/20% ratio. This prevents closely related genomes from the same genus from appearing in different splits.

The datasets are available at https://huggingface.co/datasets/macwiatrak/bacbench-essential-genes-protein-sequences (protein sequences) and https://huggingface.co/datasets/macwiatrak/bacbench-essential-genes-dna (DNA).

For gene essentiality prediction, we kept the pretrained encoder frozen and trained only a task-specific linear head on top of the gene embeddings. This head consists of a single linear layer preceded by dropout of 0.2 and layer normalisation, and is trained with binary cross-entropy loss to predict gene essentiality (1 = essential, 0 = non-essential). We trained the model for a maximum of 100 epochs with early stopping patience of 10, monitoring the macro AUROC across genomes on the validation set. For each model, we ran a learning-rate sweep to tune it on the validation set. All reported results are on the test set, and we include the best learning rate in the code (https://github.com/macwiatrak/BacBench). The weight decay for the Adam optimiser was set to 0.02 for all models. For Evo and Evo-2, performance was better without layer normalisation; therefore, we report performance without layer normalisation. Full benchmark results are available in *Table S14*.

The experiment was run on a single NVIDIA H100 Tensor Core GPU (96 GB GPU memory) with 72 CPU cores (Grace CPU) and took approximately 5 hours, including the learning-rate sweep and testing. This time does not include embedding the genomes for each model; for this, see the section on *Computational complexity analysis*.

AUROC and AUPRC curves per species were generated by selecting five strains from the test set and plotting predicted versus true labels per gene using the trained *Bacformer* (***Figure S9***). To assess the contribution of context alone, we repeated the experiment after masking all essential genes and 20% of non-essential genes, training the same linear layer only on the remaining masked embeddings and comparing results to the unmasked baseline (***Figure S9***).

To evaluate non-essential genes with the highest predicted probability of essentiality, we extracted the binary predictions on two strains from the test set—(1) *Staphylococcus aureus* (`NC_002952`) and (2) *Salmonella enterica* (`NC_003197`)—using finetuned *Bacformer* models trained on three different random seeds. The predictions for the two strains are available in *Table S3* and *Table S4*, respectively.

###### Protein function prediction

To evaluate *Bacformer’s* capability to predict protein function and benchmark against other methods, we randomly sampled a set of 300 bacterial strains from KEGG (https://www.kegg.jp/) [13] and retrieved the annotations for each genome, together with the associated genome assemblies from NCBI RefSeq [35]. We then matched the gene/protein annotations from KEGG with the genome assemblies. This resulted in 590, 657 annotated proteins out of a total of 1, 126, 510 proteins and 9, 059 unique KEGG Orthology (KO) groups. The task was a multi-class classification problem with 9, 059 classes. The 300 annotated strains were then split randomly into 60/20/20 train, validation and test sets, respectively. We chose the lowest level of annotations in KEGG, the KO groups, as our labels. We report macro metrics to ensure the models perform across distinct KO groups.

To evaluate **zero-shot** performance of different methods, we used a *k*-nearest-neighbour classifier (*k*NN) with the gene/protein embeddings extracted from each model. We embedded all genes present in the dataset for each model and evaluated performance for different values of *k* (3, 5, 7), fitting the *k*NN model on the training dataset and evaluating on the validation set. We then selected the *k* with the highest macro F1 score. Finally, we evaluated zero-shot performance on the test set. We used the NVIDIA cuML library [9] to run the accelerated *k*NN algorithm.

For **finetuning**, we stacked a linear layer on top of the frozen gene/protein embeddings, preceded by layer normalisation and dropout of 0.2, and trained it to predict the protein function label (KO group). We trained the model for 40 epochs with early stopping patience of 4, monitoring macro F1 score on the validation set. We performed a learning-rate sweep for each model on the validation set. We used the Adam optimiser with weight decay of 0.01 and cross-entropy loss for model training. Full results are available in *Table S15*.

###### Annotating proteins with unknown function

A large number of proteins across diverse bacterial genomes lack functional annotation, including many labelled as hypothetical. We finetuned the entire *Bacformer-L* (encoder + classification head) model and achieved > 98% macro F1 and macro accuracy across strains on the held-out test set. We then used the finetuned model to infer labels for proteins without KEGG annotations. For each protein, we took the top class from the softmax over more than 9, 000 classes and required that its predicted probability be at least 0.20. With thousands of classes, this is a conservative cut-off: random chance for any given class is on the order of 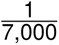, so a 0.20 posterior reflects strong model confidence and prioritises precision while leaving uncertain cases unassigned. We assigned the predicted class to all proteins whose top probability exceeded this threshold.

To visualise the results, we embedded protein vectors from the finetuned *Bacformer-L* using UMAP and coloured them by (i) true label for annotated proteins and (ii) annotated vs. unannotated status. We plotted the UMAP using the Scanpy package with 15 neighbours for the *k*NN graph and a minimum distance of 0.5 for the UMAP.

The dataset of 300 strains for protein function prediction is available at https://huggingface.co/datasets/macwiatrak/bacbench-protein-function-kegg-dna-small (DNA) and https://huggingface.co/datasets/macwiatrak/bacbench-protein-function-kegg-protein-sequences-small (protein sequences). The code for benchmarking is available at https://github.com/macwiatrak/BacBench/protein_function.

##### Phenotypic traits prediction

We collected phenotypic trait annotations by collating two major sources [40, 41]. We prepended each label with its source name so that the provenance of every annotation was explicit and potential name collisions between the two catalogues were avoided. When the same trait was present in both sources, we quantified agreement using Cohen’s κ [42] and removed traits with κ below 0.5, as these showed insufficient agreement between sources. Using the taxonomy IDs and assembly accessions provided in the phenotypic-trait datasets, we downloaded the associated genome DNA and protein sequences from NCBI GenBank (https://www.ncbi.nlm.nih.gov/genbank/). We restricted the benchmark to categorical phenotypes and filtered out traits with fewer than 500 genomes to ensure that models were trained and evaluated on a sufficiently large sample. Additionally, we removed classes with fewer than 50 samples. This resulted in 138 unique phenotypes across 24,462 genomes.

For each phenotype, we split the data into 60/20/20 train, validation and test partitions, respectively. As similar genomes often share the same phenotype, we split the data for each phenotype by genus—placing all genomes from a genus in one split—and evaluated on held-out genera, enforcing generalisation to phylogenetically distant strains. Because many traits are rare, we repeated this procedure across three independent runs with different data splits to improve robustness and obtain more stable performance estimates.

The datasets are available at https://huggingface.co/datasets/macwiatrak/bacbench-phenotypic-traits-protein-sequences (protein sequences) and https://huggingface.co/datasets/macwiatrak/bacbench-phenotypic-traits-dna (DNA).

###### Unsupervised clustering of bacterial genomes by phenotype

To evaluate whether bacterial genomes cluster by phenotype without supervision, we considered two traits: (1) environmental niche and (2) gram stain. For environmental niche, we focused on the *Bifidobacterium* genus, chosen because its members colonise a range of mammalian gastrointestinal tracts spanning infant, adult and ruminant guts and must adapt to the distinct nutrient profiles and immune pressures found in each host. Strains were extracted from the *MGnify* dataset [11] and 70 were randomly sampled across three environments. For every strain, we computed a *Bacformer* genome embedding (model pretrained on MAGs with a masked objective) and calculated pairwise cosine similarity between all strains.

For gram stain, we used the annotations from [40] to cluster strains by Gram stain. After filtering for genomes with a Gram-stain label, 14, 052 genomes remained. Each genome was embedded with *Bacformer* pretrained on complete genomes, with the genome-level vector defined as the mean of all protein embeddings. The embeddings were then projected with UMAP and the two-dimensional plot was coloured by Gram stain.

###### Phenotypic traits prediction and benchmarking

To predict a phenotypic trait from a genome-level embedding, we trained a linear model for each phenotype and method combination. The linear model is a single-layer neural network preceded by dropout of 0.1 and layer normalisation. As all labels are categorical, we optimised it to minimise cross-entropy loss. We set the maximum number of epochs to 100 and early stopping patience to 10, monitoring the validation AUROC. We ran a learning-rate sweep for each model across seeds and phenotypes and used the best-performing model for testing. Full benchmark results are available in *Table S16*. We tried balancing class weights for each phenotype due to imbalance in the label counts; however, it yielded worse validation results across models compared to training and evaluating the models without class weights.

The experiment took approximately 19 hours on a single NVIDIA H100 Tensor Core GPU (96 GB GPU memory) with 72 CPU cores (Grace CPU). This included tuning, training and testing for all models across seeds. This time does not include embedding the genomes for each model; for this, see the section on *Computational complexity analysis*.

###### Phenotypic traits annotation

To annotate the MAG corpus, we trained a separate linear model for each phenotypic trait using *Bacformer* embeddings as described above. As the phenotypic traits dataset was mainly derived from genomic isolates, not MAGs, we first examined potential distribution shifts, since the classifiers were trained on complete genomes whereas MAGs are fragmented and sometimes incomplete. *Bacformer* embeddings were generated for every MAG and each phenotype model was retrained on the MAG-derived vectors, revealing only a minor decrease in balanced accuracy. Thus, the embedding space learned on MAGs generalised well to the original tasks. We then selected phenotypes whose labels could be transferred with reasonable confidence via strain lineage, annotated the entire MAG corpus and evaluated performance where ground-truth labels were available. Balanced accuracy remained above 94% across diverse phenotypes, indicating robustness to genome type.

Phenotypes were grouped by held-out balanced accuracy: five exceeded 93% (high confidence), thirteen lay between 80% and 93% (medium) and fourteen fell between 75% and 80% (low). Predictions for every confidence tier were generated for the full MAG collection and released with accompanying confidence flags. The prediction files are available for download at https://drive.google.com/drive/folders/1PSR0mcE2fTexP9ADDVgYz3lkQ_qcyEsy?usp=sharing.

###### Polymicrobial communities annotation

To annotate bacterial genomes by phenotypes in a single sample, we selected the metagenomic sample with the largest number of MAGs (*SRS1996016*; *MGnify* [11]; human gut), containing 5, 811 MAGs. We then annotated this sample with phenotypic-trait prediction following the approach above; however, instead of a linear head we used an XGBoost model trained on top of frozen *Bacformer* embeddings to speed up annotation across all traits. For each trained trait model, we produced continuous scores for every genome using: (i) the predicted probability of the positive class for binary traits; and (ii) the softmax score of the highest-scoring class for multi-class traits. Continuous outputs yielded more informative embeddings than discrete labels. The result was a MAG × trait matrix of shape (5811, 139).

To examine whether annotated MAGs cluster by taxonomy, we kept MAGs from the 50 most prevalent species (using *MGnify* taxonomy) and built a *k*NN graph (*k* = 10) on the continuous trait profiles, then computed UMAP (distance = 0.5) and visualised points coloured by family. We also quantified which phenotypic traits drive the community structure by running PCA on the MAG × trait matrix and computing a variance-weighted loading score per trait for the top 20 principal components. This score ranks traits by their contribution to variance. Finally, we visualised lactose utilisation, one of the highest-scoring phenotypes, using the computed UMAP. We used the Scanpy package [27] to compute the *k*NN, UMAP, and PCA, and to generate the visualisations.

###### Identifying phenotype-specific genes

We traced phenotype-linked genes for four traits: sporulation, motility type, anaerobic growth and environmental niche. The first three used the complete-genome trait data described above; environmental-niche prediction required a new dataset. Families, genera and species occurring in multiple habitats were chosen so that the model could not rely on coarse taxonomic signals. From this pool we selected *Lactobacillus johnsonii* species and the genus *Prevotella* for detailed study because they occupy distinct niches yet share close taxonomic ranks, facilitating separation of lineage effects from habitat-specific adaptations. The MAGs for both were extracted from *MGnify* [11].

A separate *Bacformer* model was finetuned for each phenotype. Each dataset was randomly split into 60% training, 20% validation, and 20% test sets. For sporulation, motility, and anaerobic growth, we initialised from the *Bacformer* checkpoint pretrained on complete genomes; for environmental-niche models, we used the MAG-pretrained checkpoint.*Input × Gradient* attribution [43] was applied to produce an importance score for each gene in every test genome, by summing over the contextualised protein-embedding vectors. To assess biological plausibility, we averaged scores across all occurrences of each annotated protein, discarded genes lacking annotations, and compared mean scores of proteins matching relevant keywords (e.g., “flagellum,” “spore formation,” “anaerobe”) to the remainder.

Sporulation served as a convenient qualitative benchmark due to its well-characterized genetic machinery. Ranking proteins by attribution successfully recovered known sporulation factors, especially the *spo* gene set—among the highest-scoring hits. After excluding proteins lacking locus tags in at least three strains, we averaged the top 100 ranked proteins across strains, clustered them using the Leiden algorithm (resolution 0.1, *k* = 5), and identified four functionally coherent modules corresponding to successive stages of spore formation.

We repeated this procedure to investigate environment-specific adaptation in *L. johnsonii*. For each host gut, the 100 most important proteins were averaged across strains and visualised with UMAP, with points coloured by environment. Manual inspection highlighted several proteins previously implicated in host-specific colonisation, confirming that the model identified meaningful determinants of niche preference. The environment-specific gene table for *L. johnsonii* is available in *Table S5*.

Because many genomes lack reliable gene names or product descriptions due to fragmentation and annotation errors, we repeated the analysis using the protein family (i.e. clusters) labels assigned during pretraining. As all proteins carry such a label, this approach ensured no sequences were discarded. Within the *Prevotella* test set, we computed mean importance for each family, averaged the top 100 families per environment, and visualised the results with UMAP, again revealing clear environment-specific clusters.

##### Predicting antimicrobial resistance

We leveraged the antibiotic sensitivity readings from the NCBI AST browser (https://www.ncbi.nlm.nih.gov/pathogens/ast), which contains hundreds of thousands of antibiotic-susceptibility test (AST) records for a diverse set of antibiotics. Using the genome assembly identifiers provided by the NCBI AST browser, we downloaded the DNA and protein sequences for each genome from NCBI GenBank (https://www.ncbi.nlm.nih.gov/genbank/) and matched them with the antibiotic resistance readings. This left us with 26, 052 unique genomes with matched antibiotic resistance labels. We then processed the antibiotic resistance readings into *(i)* binary and *(ii)* regression labels, described below.

For **binary** prediction, antibiotic sensitivity labels from the NCBI AST browser of either *sensitive* (S) or *resistant* (R) were used for training and testing. If the antibiotic sensitivity test had no S/R label, it was not included. We removed antibiotics that (1) had fewer than 500 available unique genomes in total or (2) had fewer than 50 unique genomes per class. This was motivated by the need to ensure that every classifier was trained on a sufficiently large and reasonably balanced dataset; with fewer than 500 genomes overall, or fewer than 50 genomes in either class, the resulting model would suffer from poor statistical power and unreliable performance estimates. This resulted in 37 unique antibiotics. *Table S17* shows the number of available genomes per drug, including the number of susceptible and resistant genomes. The number of available readings varies strongly between antibiotics, which partly explains the high variance in per-drug performance. For **regression** MIC prediction, the NCBI AST browser quotes minimum inhibitory concentrations (MICs) as < *x*, ≤ *x*, = *x*, ≥ *x* and > *x*. These strings were translated to a numerical value (*y*) by setting *y* = *x* if NCBI quoted MIC = *x*, ≤ *x* or ≥ *x*; *y* = 2 × *x* if MIC was quoted as MIC > *x*; and *y* = 0.5 × *x* if MIC was < *x*.

We filtered out antibiotics with fewer than 500 available readings to ensure that the models were trained on a sufficiently large sample. This resulted in 56 antibiotics. To dampen the long tails, we normalised the MIC with a log(1 + *x*) transformation.

Because many antibiotics have relatively few labelled strains and because resistance mechanisms are often species-specific, we do not use a genus-level phylogeny-aware split for this task. Instead, for each antibiotic and random seed, we perform a random train/validation/test split in a 70/10/20% ratio, tune the model on the validation set, and report performance on the held-out test set. We repeat this procedure across three random splits to reduce sensitivity to a single partition of the data.

###### Benchmarking

To compare the predictive power of various models’ representations, we extracted the frozen genome representation for all proteins in the genome and averaged across the protein embeddings to create genome embeddings for each model. To predict antibiotic resistance, we trained a linear model for each drug and method combination. The linear model is a single-layer neural network preceded by dropout of 0.1 and layer normalisation. We trained the models separately for the *(i)* binary and *(ii)* regression MIC prediction cases. We optimised the former for binary cross-entropy loss and the latter for mean squared error loss. We trained all models for a maximum of 100 epochs with early stopping patience of 10, monitoring validation AUROC in the binary setup and validation *R*^2^ in the regression setup. We ran a learning-rate sweep for each model across seeds and drugs and used the best-performing model for testing. We set the weight decay in the Adam optimiser to 0.01. Full benchmark results are available in *Table S18*.

We also experimented with training a multi-task linear model, which simultaneously predicts antibiotic resistance of a genome to multiple drugs; however, it performed worse than a separate linear model for each drug.

The experiment took approximately 50 hours on a single NVIDIA H100 Tensor Core GPU (96 GB GPU memory) with 72 CPU cores (Grace CPU). This included tuning, training and testing for all models across seeds. This time does not include embedding the genomes for each model; for this, see the section on *Computational complexity analysis*.

The AMR datasets are available at https://huggingface.co/collections/macwiatrak/bacbench, and code for benchmarking is available at https://github.com/macwiatrak/BacBench/tree/main/bacbench/tasks/antibiotic_resistance.

###### Identifying AMR-associated genes

*Bacformer* is the first genomic LM that enables deep-learning embedding representations of entire genomes for which finetuning on multiple individual tasks (such as AMR prediction) is computationally feasible (see the Computational complexity analysis section).

Finetuning *Bacformer* to predict AMR was performed for three example bacteria/antibiotic pairs, chosen to span a range of drug actions and phylogenetic diversity and because each had more than 500 publicly available genomes in NCBI with at least 200 in both the S/R classes. These include *E. coli* resistant to ampicillin (*N* = 5, 876 genomes with ASTs: 3, 305 S / 2, 571 R); *Acinetobacter baumannii* resistant to gentamicin (*N* = 970: 217 S / 753 R); and *Salmonella enterica* resistant to tetracycline (*N* = 8, 828: 4, 339 S / 4, 489 R).

The genomes from each species were split into training/validation/test sets (60/20/20). We then finetuned the *Bacformer* transformer to predict AMR from average protein embeddings using a single-layer neural classifier with layer normalisation, dropout 0.2, learning rate 0.00015, 10% warm-up and the Adam optimiser [15].

The end-to-end model was finetuned for up to 100, 000 steps, with validation every 250 steps monitored by AUROC and early stopping after 30 validation steps. Finetuning on one A100 GPU took ∼18–24 hours for each species/antibiotic pair.

Attribution scores for each protein were derived by *in silico* knockout for each finetuned model (each species/drug pair) using Captum (v0.8; https://captum.ai/api/feature_ablation.html) [44]. Masking all dimensions of each protein embedding provided a summed score for each protein, one at a time; the attributions from all 480 ablated dimensions were summed and then divided by 480 to give an attribution score for each protein. This gives a score reflecting the logit for model prediction with the protein present, minus the prediction when the protein is absent (ablated).

AMR Finder Plus [45] was used to annotate the proteins in each genome, and the annotations were stored alongside their attribution scores. We used a one-sided *Mann-Whitney* U rank-sum test to determine if the attribution scores for each annotation group (i.e., each element symbol or gene name) were greater than the background distribution of all protein attributions. Given the large size of our dataset (over 1 million proteins per species), the sampling distribution of the *Mann-Whitney* U statistic is well approximated by a normal distribution. We therefore calculated p-values based on z-scores of the U-statistics. The resulting *p*-values were adjusted for multiple comparisons using the *Benjamini-Hochberg* procedure to control the false discovery rate at 0.05 (*Tables S6* (E. coli and ampicillin), *S7* (A. baumannii and gentamicin), and *S8* (Salmonella and tetracycline)) for each species/drug pair.

##### *De novo* genome generation

Using a pretrained *Bacformer* model with an autoregressive objective on complete genomes, we computed perplexity, a metric that indicates how well the model predicts the next element in a sequence. We then analysed performance across test genomes, including species not seen during training.

###### Generating genomes with a broad range of protein functions

We assessed whether *Bacformer* can generate genomes that encompass a broad range of essential functions—an important advance over existing generative models that are limited to designing individual molecules [46, 47, 48, 49, 50] or relatively simple multi-component systems [51].

In its original formulation *Bacformer* represents each protein with a continuous embedding produced by a protein language model from the raw amino-acid sequence. This fine-grained representation allows residue-level reasoning, yet it renders generation impractical: a decoder (the autoregressive component that generates sequences in transformers), which takes a vector in ℝ*^D^* as input (where *D* is the dimension of the embedding), would also have to predict an exact vector output in **R***^D^*, an unbounded task, rather than producing a symbol from a finite vocabulary.

To enable genome generation we discretise the input. A vocabulary *F* = {1, *…*, *F* } of *F* = 50, 000 discrete protein clusters is obtained by leveraging the results of unsupervised clustering used for pretraining. Each protein is mapped to its cluster index so that a genome *f* becomes a sequence

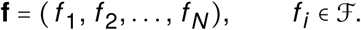

The model now looks up a learnable token vector **x***_i_* = *E_fi_* from an embedding matrix *E* ∈ ℝ*^F^*^×*D*^, exactly as in word-piece language models [14]. All other components of the architecture remain unchanged.

We initialise *E* with the rows of the protein family cluster *classification head W* ∈ ℝ*^F^*^×*D*^ learned during the autoregressive MAG pretraining. As *W* already captures the distinguishing features of each cluster in the same latent space, this transfer (i) aligns the new discrete-token model with the previously learned semantic manifold, (ii) speeds up convergence, and (iii) avoids catastrophic forgetting of cluster-specific information.

Starting from this initialisation we further pretrained the discrete-token model on the complete-genome corpus, following the identical optimisation schedule used for the continuous-embedding *Bacformer*. This procedure leverages uninterrupted chromosomal context while retaining compatibility with sequence generation tasks.

We prompted pretrained autoregressive protein-cluster *Bacformer* to generate a set of protein sequences ordered by their chromosomal positions from a prompt of 500 proteins starting from the origin of replication for *Staphylococcus aureus* (NCBI RefSeq ID `GCF_001019415.2`) and *Escherichia coli* (NCBI RefSeq ID `GCF_003627195.1`). The origin of replication represents the natural starting point for bacterial chromosome replication, providing a biologically meaningful context for genome generation.

We assessed functional diversity by mapping protein families to *KEGG* BRITE ontology categories [13]. We extracted *KEGG* annotations for bacterial genes by downloading gene annotations and genome assembly links from NCBI RefSeq and GenBank via the *KEGG* website. We processed the downloaded genomes and embedded them using the *ESM-2*model (12 layers, 35M parameters) [2]. After discarding proteins without *KEGG* annotations, we obtained over 600,000 annotated proteins. We assigned *KEGG BRITE* functions to our protein family clusters using *k* nearest neighbour classification (*k* = 7) based on centroid embeddings with Euclidean distance. We validated the annotation quality by examining a random sample of protein clusters and their *k*-nearest-neighbour assignments, and verified that the overall distribution of *KEGG* protein functions across protein families resembled real-world distributions. For generated genomes (excluding the 500-protein prompt), we assigned each protein family a *KEGG BRITE* function and compared the distribution of true versus predicted *KEGG* labels. This analysis confirmed that *Bacformer* generates genomes with functional repertoires that closely match those of true genomes (***Figure 4B***; ***Figure S13***).

To ensure that the model can generate diverse genomes and does not simply recapitulate genomes seen during training, we generated multiple genomes using the same prompt while varying the temperature parameter to introduce controlled diversity, and we computed the distance between the generated genomes and those in the pretraining corpus. The results (***Figure S13***) show that *Bacformer* generates diverse genomes that are not merely replicas of any single training genome.

###### Genome completion

To test *Bacformer*’s ability to recognise genomes with missing sections, we used the held-out set (*N* = 9, 688) of complete genomes and systematically removed 5%, 10%, 25% or 50% of the proteins by trimming sequentially from the genome’s end. Both truncated and intact genomes were embedded with *Bacformer* (trained on complete genomes that were not part of the held-out set, using the autoregressive objective) and placed in the same embedding space for comparison. For each truncated genome, we measured how highly its intact counterpart ranked among all genomes in the embedding space, reporting the mean reciprocal rank (MRR), where 1.0 indicates a perfect match.

To visualise the effect of missing data, we projected *Bacformer* embeddings of full genomes and their 50%-ablated versions onto two dimensions using UMAP (10 neighbours). As a non-contextual baseline, we performed the same procedure with averaged *ESM-2* protein embeddings, omitting the *Bacformer* encoder (***Figure S14***).

###### Retrieving protein sequences from contextual embeddings

*Bacformer* generates contextual protein representations by associating each protein with one of 50, 000 predefined protein clusters, rather than encoding each unique protein sequence individually. This clustering approach addresses the challenge posed by the immense diversity of protein sequences, enabling the model to generalise across related proteins while maintaining computational tractability.

By inverse mapping each embedding to its corresponding protein cluster, it becomes possible to retrieve the native protein sequence or a close homolog from a large candidate pool. This is particularly valuable for applications such as genome completion and synthetic biology, where identifying the most likely protein sequence from an embedding is essential.

To facilitate this mapping, *Bacformer*’s output embedding for each gene **h***_i_* ∈ ℝ*^d^* is trained using a contrastive loss. This loss encourages the *Bacformer* embedding to closely match the *ESM-2* embedding of the same protein within a batch, while remaining distinct from embeddings of other proteins. As a result, the learned embedding space supports accurate retrieval of protein cluster identities from contextual representations.

To achieve this, both sets of embeddings are *l*_2_-normalised and similarity is measured across protein embeddings using the cosine similarity, where sim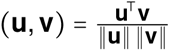 and temperature *τ* = 1, the noise-contrastive loss for a positive pair (**h***_i_*, **e***_i_*) is

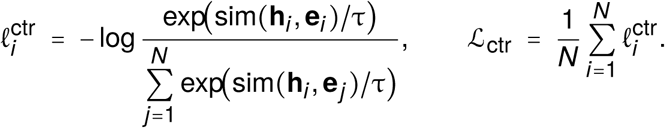

Masked-family prediction uses the usual cross-entropy with the true family labels at masked positions and the logits from *Bacformer*’s classification head. The combined training objective is

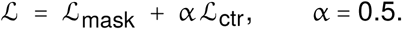

To train the model with the new loss we initialised the model weights from the MAG-pretrained *Bacformer* (26M) with masked objective and trained it on the complete genome corpus following the same pretraining setup as with the standard *Bacformer* model without contrastive loss. The *L*_mask_ loss was computed on all masked tokens while the *L*_ctr_ loss on all non-special tokens, we then back-propagated the combined loss.

Using a trained model, we ran inference on test genomes and calculated the cosine similarity between the real and predicted embeddings. For both masked and unmasked proteins, we measured the cosine similarity between the positive (true) and negative (false) protein embeddings. Furthermore, we assessed model performance using the recall@K metric, which measures how often the correct embedding is found among the top K most similar embeddings. While the model performed strongly on unmasked proteins, it achieved significantly worse results in retrieving the true embedding at low *K* values for masked proteins. This highlights the importance of the *Bacformer* transformer backbone’s capacity to update protein embeddings, as the contextualised representations differ significantly from the initial protein embeddings used as input.

###### Generating genomes with desired properties with conditional finetuning

To explore how *Bacformer* could be applied to generate entire bacterial genomes endowed with specific properties, we selected two categorical traits from the phenotypic dataset described above: oxygen requirements (specifically *aerobic*, *anaerobic* and *facultative*); and optimal growth temperature (*thermophilic* (> 80°C), *psychrophilic* (< 20°C) and *mesophilic* (20–40°C)).

We randomly split each dataset into train, validation and test splits with a 60/20/20% ratio. For training, we initialised the model weights with the protein family cluster *Bacformer* model pretrained on complete genomes described in the *Generating genomes with a broad range of protein functions* section above. For each phenotype, we introduced a property token at the start of each protein family sequence (***Figure 4F***). The property token was initialised randomly and trained from scratch during training. We then finetuned *Bacformer* to generate protein family sequences conditional on a phenotype token, monitoring the validation loss separately for each trait.

Following finetuning, we used each trained model to generate a representative subset of genomes (*N* = 20) from the test set given a prompt consisting of a property token and a set of 500 protein families. For comparison, we repeated the procedure without the property token. We then compared the generated protein families with and without the property token, excluding the 500 prompt tokens. To evaluate the difference between the protein family sequences generated with and without the property token, we associated the protein family tokens with *KEGG Orthology* (KO) groups [13], allowing us to identify specific orthologous protein groups that differ between the generated sequences. To do this, we used the annotated proteins from *KEGG* for bacterial genomes (see *Generating genomes with a broad range of protein functions*). As a protein may belong to multiple orthologous groups, we assigned protein family cluster tokens, using their centroids, to an orthologous group if 2 out of 10 nearest neighbours from the space of annotated proteins had the group label. We used Euclidean distance to compute the *k*NN. The threshold was selected after experimenting with different values and evaluating the distribution of the number of KO groups assigned to protein family tokens. After assigning each protein family token its KO groups, we aggregated the KO groups based on the generated protein families for the genomes generated with and without the property token and computed the log_2_ fold-change to evaluate the difference between the two. Finally, to evaluate statistical significance, we performed Fisher’s exact test [29] using the aggregated KO counts from the genomes with and without the property token. We then qualitatively analysed the positively and negatively enriched statistically significant KO groups (*p* < 0.075).

Full validation of synthetic genome viability remains outside the scope of this study and requires complex genome engineering [52]. Nevertheless, these findings underscore the promise of contextual, evolution-informed models for rational genome design, laying the groundwork for more sophisticated generative designs in microbial genomics.

### SUPPLEMENTARY FIGURES

**FIGURE S1.**
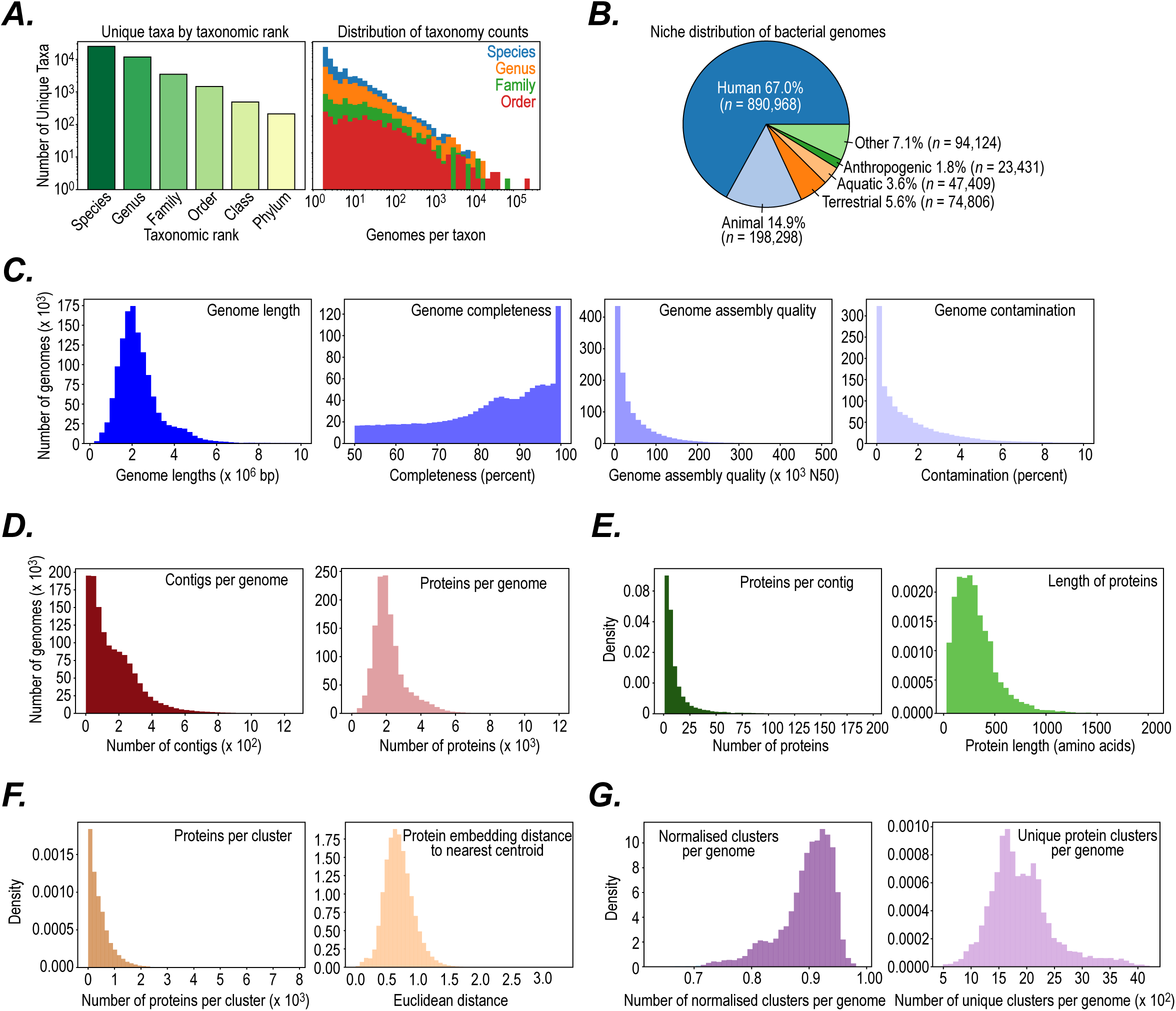
Statistics of the MAG pretraining corpus. (**A**) Taxonomy ranks statistics of the MAG Corpus. Number of unique taxa across taxonomic ranks (*left*). Distribution of genomes per taxon across taxonomy ranks (*right*). (**B**) Pie chart showing the distribution of bacterial genomes across environmental niches. (**C-E**) MAG corpus quality control distributions across genomes and metrics. (**F-G**) Protein family clusters statistics used for pretraining. (**F**) Distribution statistics of protein family clusters used for pretraining including number of proteins per cluster (*left*) and Euclidean distance of each *ESM-2* protein embedding (average of all residues) to the nearest cluster centroid (*right*). (**G**) Distribution statistics on the number of protein family clusters per genome: the number of unique protein family clusters divided by the total number of proteins in the genome (giving the normalised clusters per genome (*left*) and the absolute number of unique protein family clusters per genome (*right*), showing that most proteins in each genome were assigned to distinct clusters.

**FIGURE S2.**
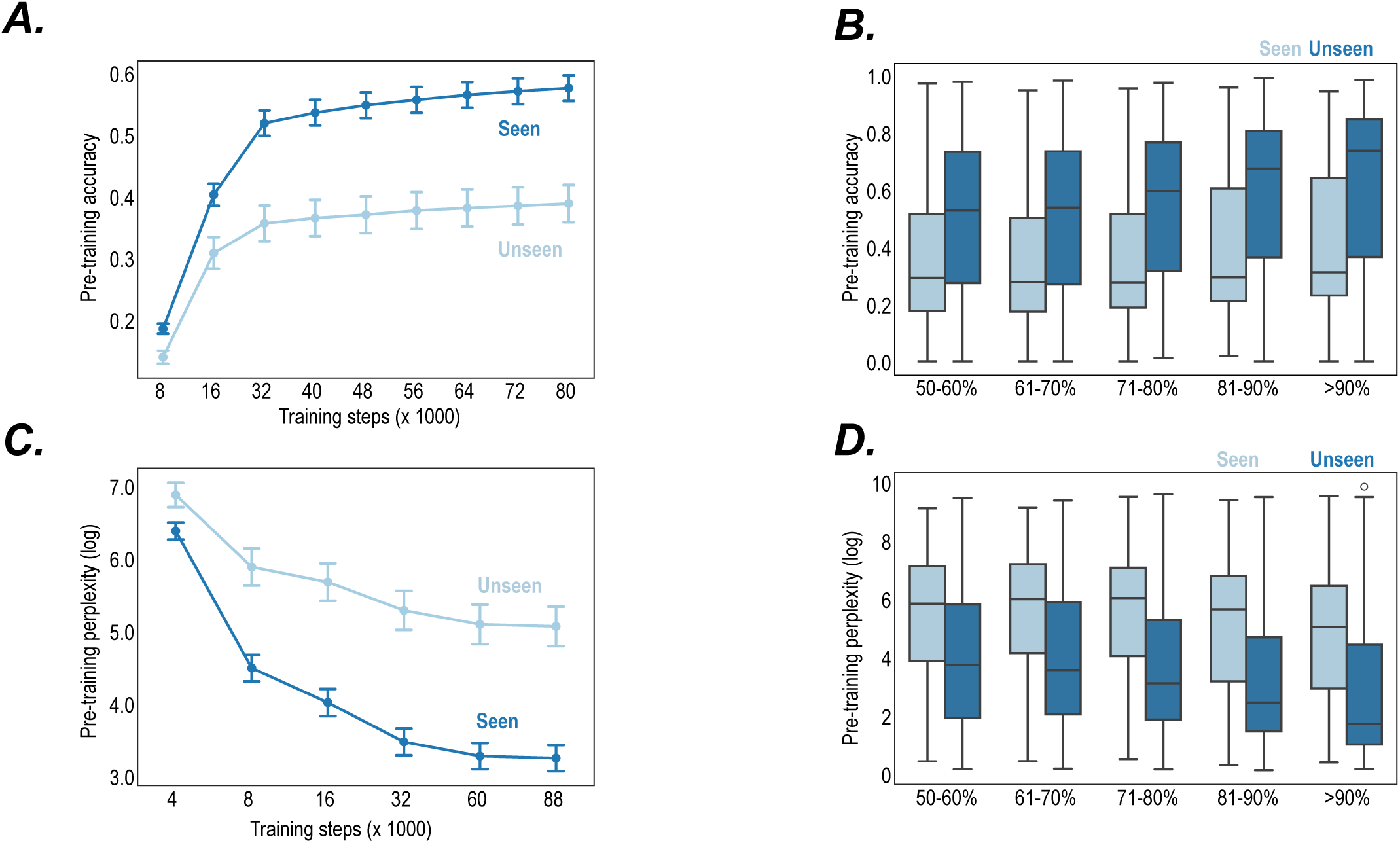
Pretraining performance of Bacformer. (**A,B**) Pretraining performance of *Bacformer* on the held-out evaluation set (*N* = 49,213) on the masked protein family prediction objective. Error bars represent standard error computed from 15 bootstrap resamples. (**A**) Accuracy across training steps on seen and unseen taxa during training. (**B**) Accuracy across genome completeness bins, stratified by whether the taxa has been seen during training (median and IQR shown; whiskers extend 1.5 × IQR). (**C, D**) Pretraining performance of *Bacformer* on the held-out evaluation set (*N* = 49,213) with autoregressive protein family prediction objective. Error bars represent standard error computed from 15 bootstrap resamples p < 10^−5^ paired Wilcoxon signed-rank test. (**C**) Perplexity across training steps on seen and unseen taxa during training. (**D**) Perplexity across genome completeness bins, stratified by whether the taxa has been seen during training (median and IQR shown; whiskers extend 1.5 × IQR).

**FIGURE S3.**
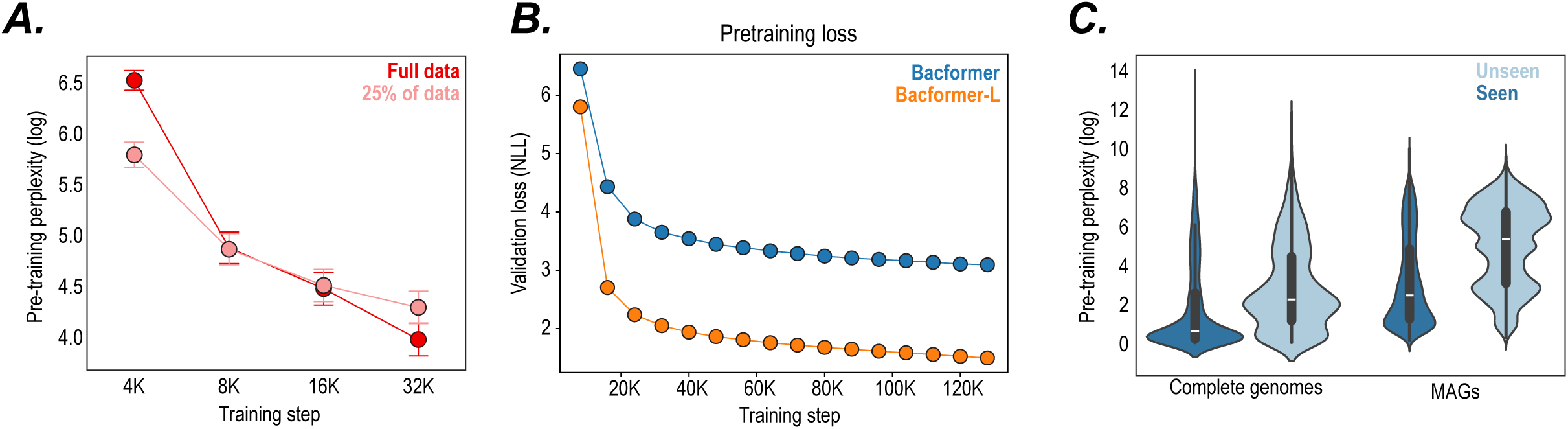
Bacformer scales with model & dataset size. (**A-C**) Pretraining performance of *Bacformer* on the held-out evaluation set (*N* = 49,213). (**A**) Perplexity across training steps for model with 25% and 100% of data. Error bars represent standard error computed from 15 bootstrap resamples. (**B**) Validation loss (Negative log-likelihood) for Bacformer (26M parameters) vs Bacformer-L (332M parameters) across training steps. (**C**) Perplexity on complete vs metagenome assembled genomes (MAGs), including genomes from seen and unseen taxa during training (median and interquartile range shown).

**FIGURE S4.**
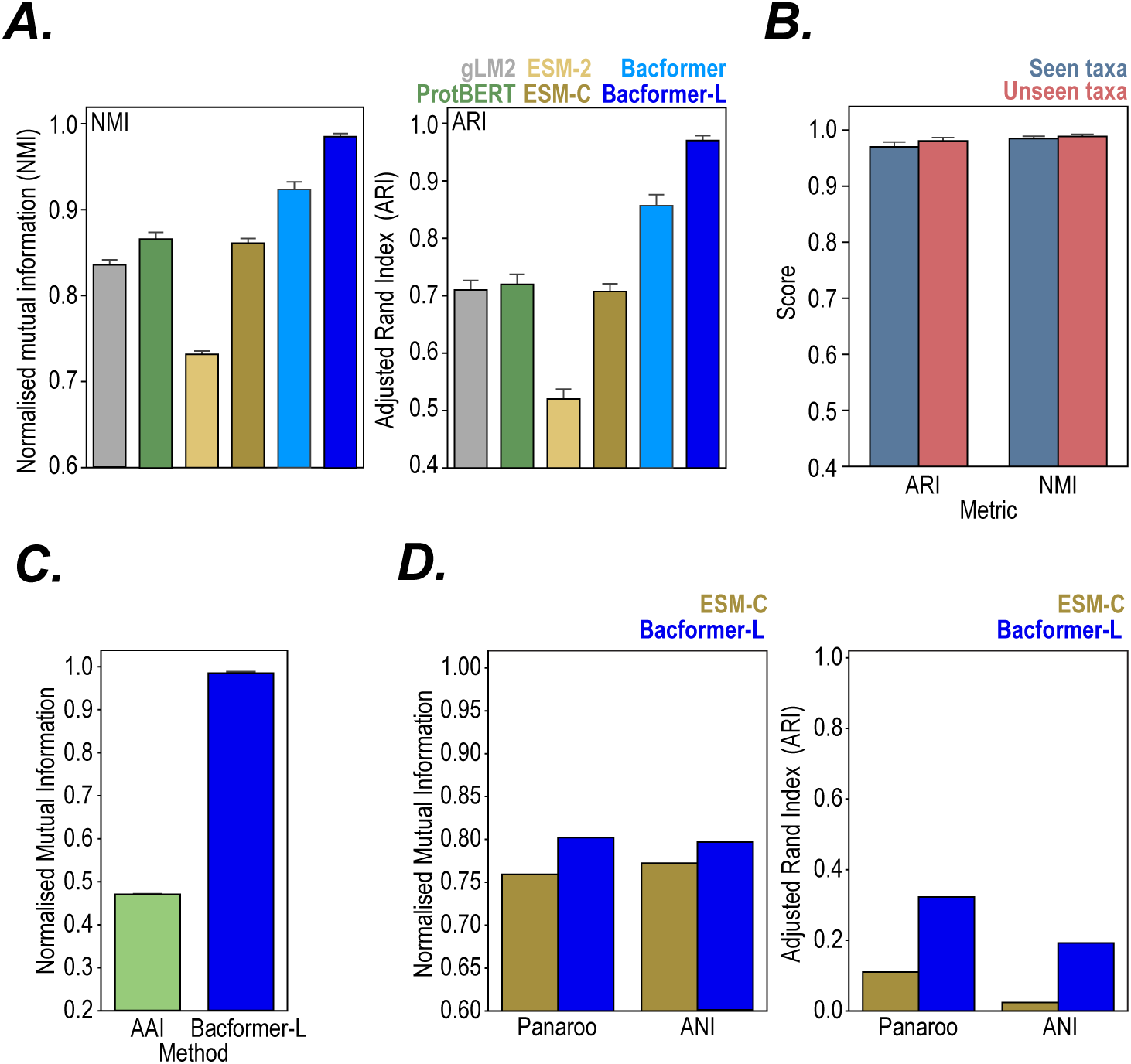
Additional genome clustering benchmarking. (**A**) Performance on the genome clustering by species task on a set of diverse MAGs (*N* = 6,071) from MGNify. We compared protein-based and mixed-modality LMs based on Adjusted Rand Index (ARI) and Normalised Mutual Information (NMI). Metrics vary from 0 to 1; higher is better. Error bars denote 95% confidence intervals estimated from 10 independent bootstrap resamples, each using 80% of the data. (**B**) Genome clustering performance on the seen (*N* = 6,071) and unseen taxa (*N* = 2,612) based on Adjusted Rand Index (ARI) and Normalised Mutual Information (NMI). Metrics vary from 0 to 1; higher is better. Error bars denote 95% confidence intervals estimated from 10 independent bootstrap resamples, each using 80% of the data. (**C**) Comparison of *Bacformer-L* vs Average Amino-acid Identity (AAI) on the strain clustering task (*N* = 6,071) as measured by Normalised Mutual Information (NMI). (**D**) Congruity of *Bacformer-L* and ESM-C with Average Nucleotide Identity (ANI) and Panaroo methods on a set of *E. coli* genomes (*N* = 2,479) as measured by Normalised Mutual Information (NMI) and Adjusted Rand Index (ARI). Metrics vary from 0 to 1; higher is better.

**FIGURE S5.**
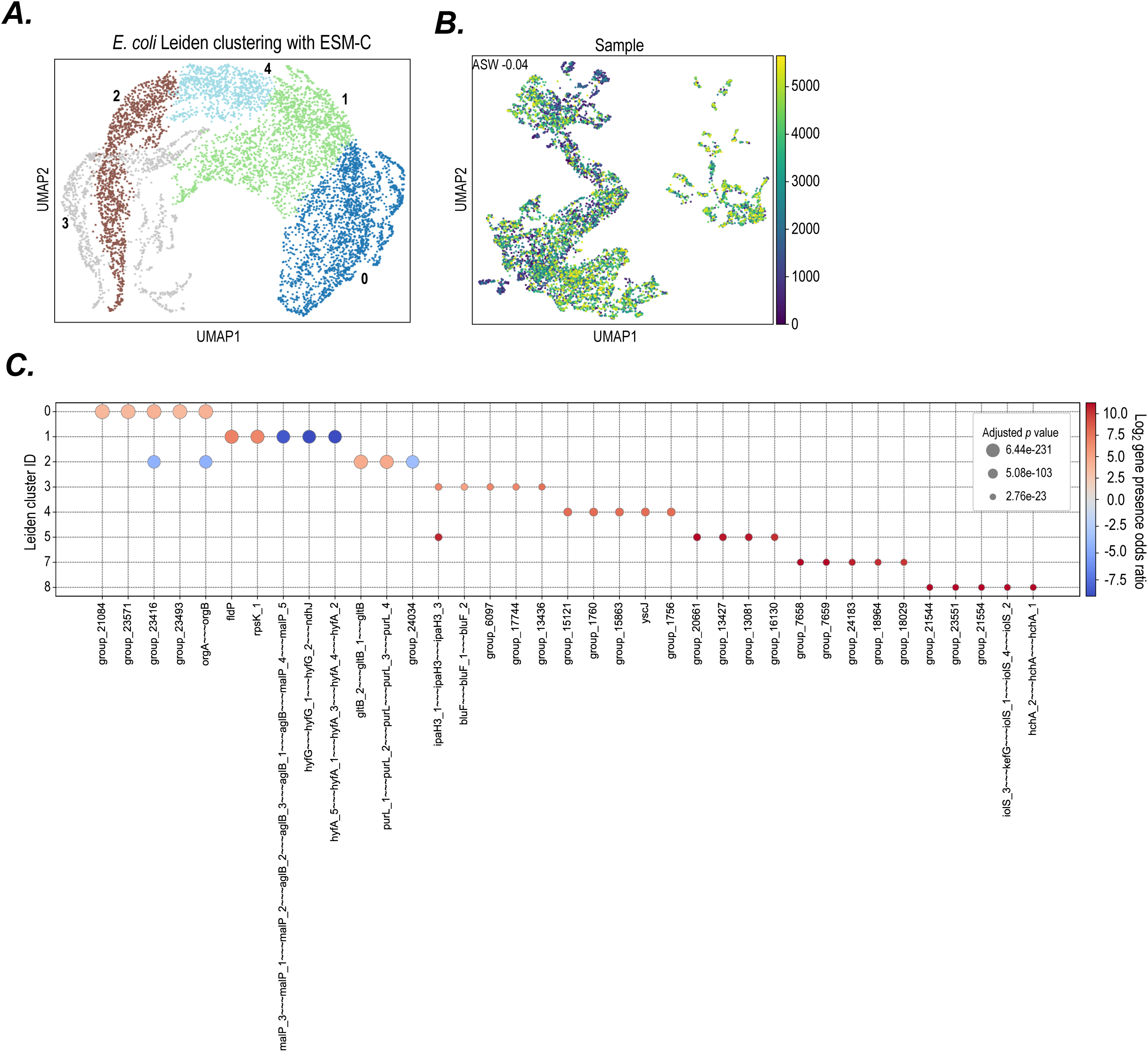
Bacformer clustering results for *Escherichia coli.* (**A**) UMAP plot of *E. coli* strains from the *MGNify* dataset using genome embeddings from *ESM-C* annotated by their Leiden cluster labels. (**B**) UMAP plot of *E. coli* strains from the *MGNify* dataset using genome embeddings from *Bacformer* annotated by the sample ID. ASW is Average Silhouette Width (0-1, higher is better)). (**C**) Top 5 cluster-specific genes based on gene presence/absence according to the p-value for 5 Leiden clusters from the *MGNify E. coli* population. The color represents the log2 gene presence odds ratio, showing whether the gene presence or absence is enriched in the subpopulation.

**FIGURE S6.**
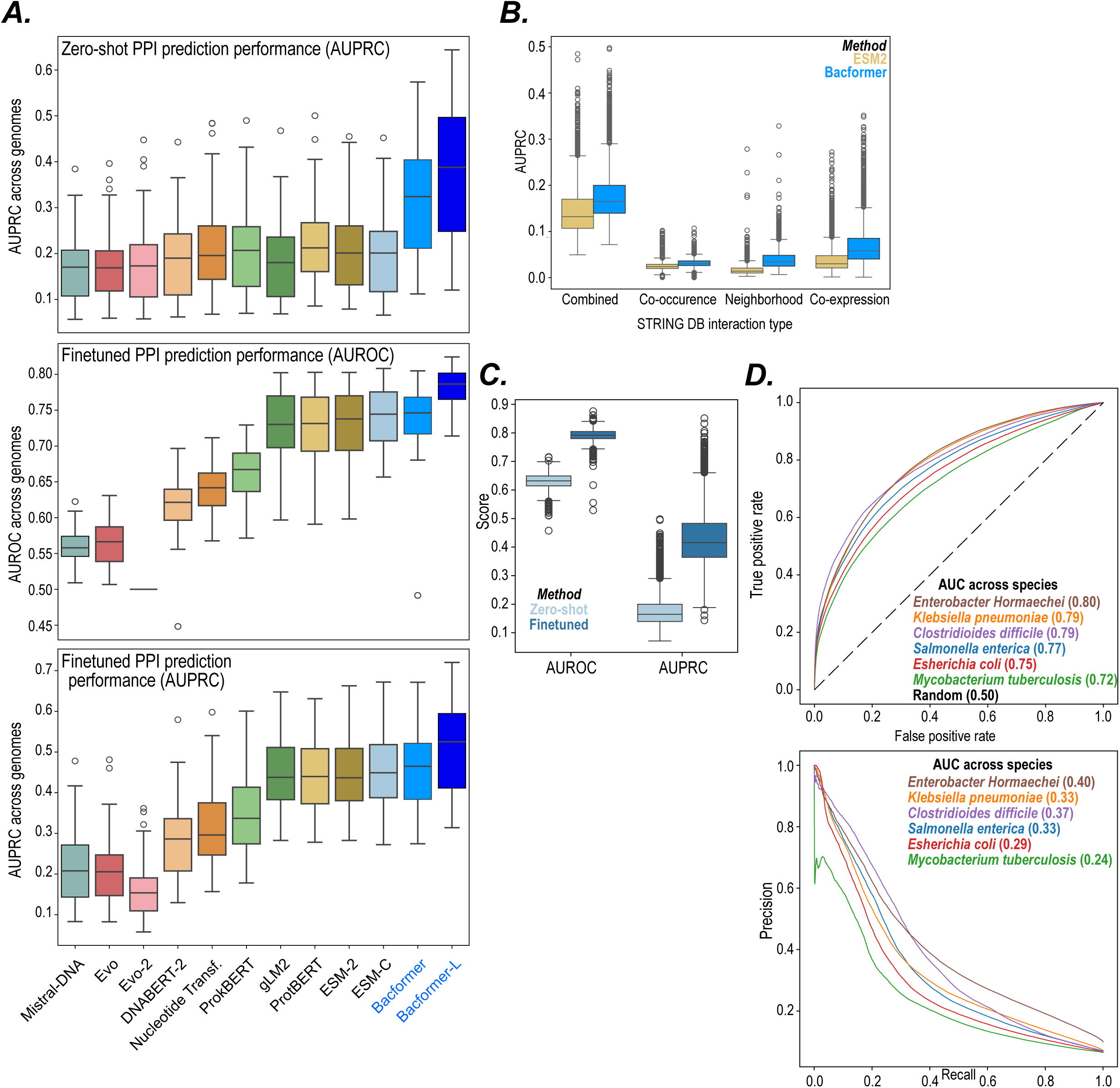
Additional results on protein-protein interaction prediction task. (**A**) Benchmarking performance across different genomic LMs on a randomly sampled set of strains (*n* = 54) from STRING DB using area under the receiver operating characteristic (AUROC) and Precision-Recall (AUPRC) curves. In the zero-shot setting (*top*) the interaction scores were calculated using cosine similarity between protein representations on held-out genomes (*see Supplementary Methods*). In the finetuning setting (*middle and bottom*) we trained a linear layer on top of the averaged protein representations. Line and boxes represent median and interquartile range (IQR) respectively; whiskers extend 1.5 × IQR beyond Q1 and Q3. Statistical significance was assessed across held-out genomes using paired two-sided Wilcoxon signed-rank tests comparing *Bacformer* or *Bacformer-L* against each baseline method (*p* < 0.005). The zero-shot AUROC plot is shown in Figure 2B. (**B**) Zero-shot predictions (evaluated using the area under the precision-recall curve; AUPRC) for *Bacformer* (*blue*) and its base pLM model - *ESM-2* (*yellow*) across different STRING DB interaction types on held-out genomes (*N* = 2,088). The interaction scores were calculated using cosine similarity between protein representations. Lines and boxes represent the median and interquartile range (IQR) respectively, with whiskers extending 1.5 × IQR beyond Q1 and Q3. Statistical significance was assessed across held-out genomes using paired two-sided Wilcoxon signed-rank tests (p < 10^−5^). (**C**) Finetuning entire *Bacformer* (encoder + classification head) model significantly boosts performance on the protein-protein interaction prediction task across held-out genomes (*N* = 2,088). We used STRING DB combined scores as labels. Line and boxes represent median and interquartile range (IQR) respectively; whiskers extend 1.5 × IQR beyond Q1 and Q3. Statistical significance was assessed across held-out genomes using paired two-sided Wilcoxon signed-rank tests (p < 10^−5^). (**D**) Receiver Operating Characteristic (ROC; *top*) and Precision-Recall (PR; *bottom*) curves for Bacformer finetuned on protein-protein interaction task with data from STRING DB across diverse species.

**FIGURE S7.**
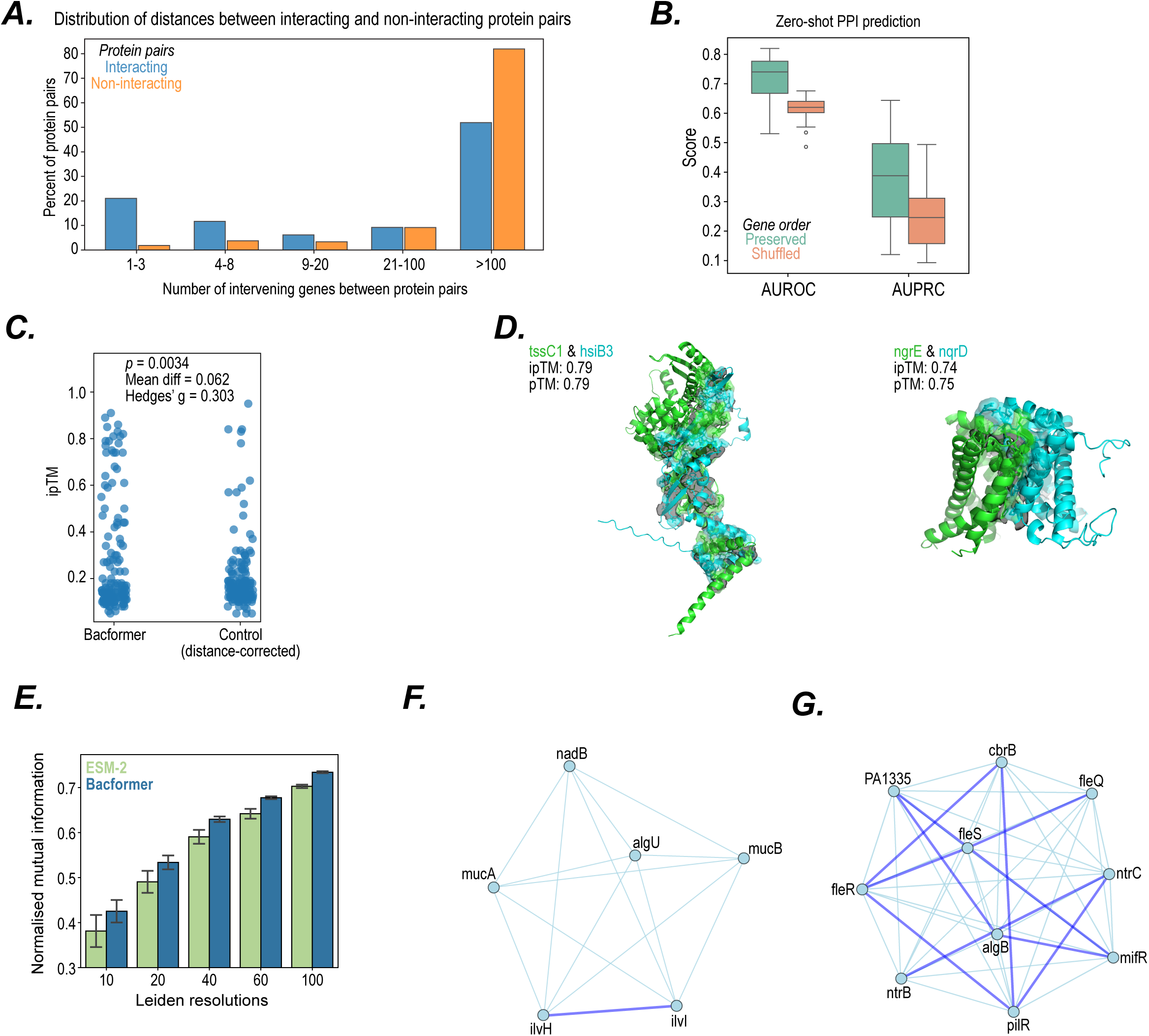
Bacformer predictions of protein-protein interaction across species. (**A**) Distribution of distance between interacting and non-interacting protein pairs using STRING DB across diverse bacterial genomes (*N* = 54, combined score with binary threshold of 0.6). The distance between the protein pairs was measured by the number of genes separating the two proteins of interest. (**B**) Ablation study results on the protein-protein prediction task using *Bacformer-L* with preserved vs randomly shuffled gene order using STRING DB benchmark (*N* = 54). Line and boxes represent median and interquartile range (IQR) respectively; whiskers extend 1.5 × IQR beyond Q1 and Q3. (**C**) ipTM scores of 150 top scoring protein pairs in *Pseudomonas aeruginosa* PAO1 genome using finetuned *Bacformer-L* vs ipTM scores of 150 distance-matched randomly selected protein pairs. The distance between the protein pairs was measured by the number of genes separating the two proteins of interest. The p-value were calculated using one-sided distance-stratified permutation test. (*D*) Predicted protein complex structures with *AlphaFold3* between two protein pairs with a high score from *Bacformer-L* model finetuned on the protein-protein interaction task. The ipTM score indicates the probability of the two proteins binding together and pTM score is a measure of overall predicted structure confidence. Both metrics are bound between 0-1, higher is better. (**E**) Evaluation of zero-shot gene clustering on *P. aeruginosa* PAO1 using co-expression and protein representations from *Bacformer* and *ESM-2* across Leiden resolutions. Higher normalised mutual information indicates higher overlap between clusterings. The error bars represent standard error across different numbers of neighbours used for Leiden clustering. (**F & G**) Gene networks from *P. aeruginosa* PAO1 genome extracted using zero-shot *Bacformer* contextual protein embeddings at Leiden resolution of 40 revealing (**F**) the algU gene network consisting interacting genes such as mucA and mucB and (**G**) the fleQ gene network consisting interacting genes such as fleS, fleR and algB.

**FIGURE S6.**
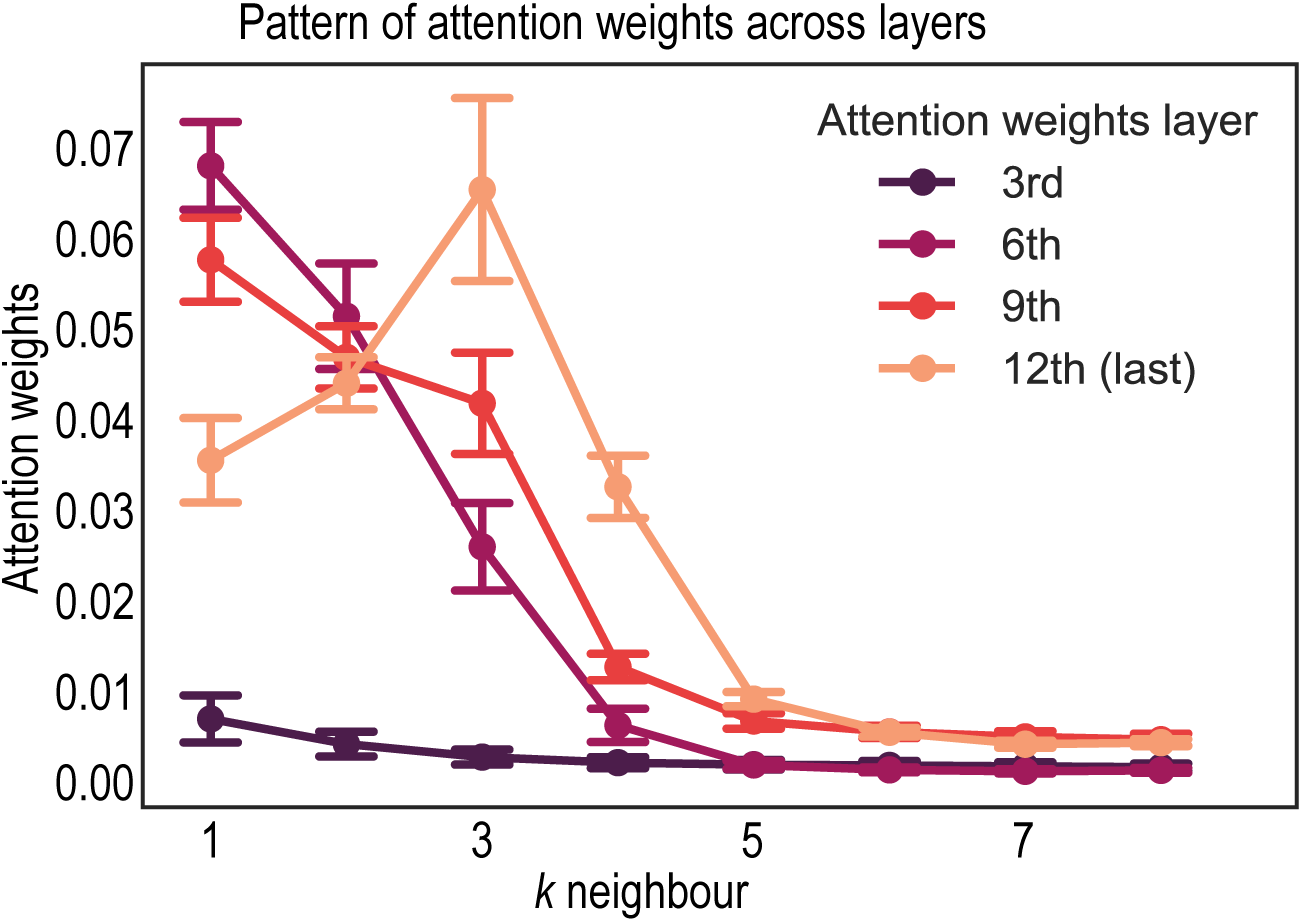
Attention weights analysis. Attention weights across layers and neighbours from the *Bacformer* model. The attention weights were computed by averaging attention heads across proteins from *N* = 45 genomes. Error bars represent standard error.

**FIGURE S9.**
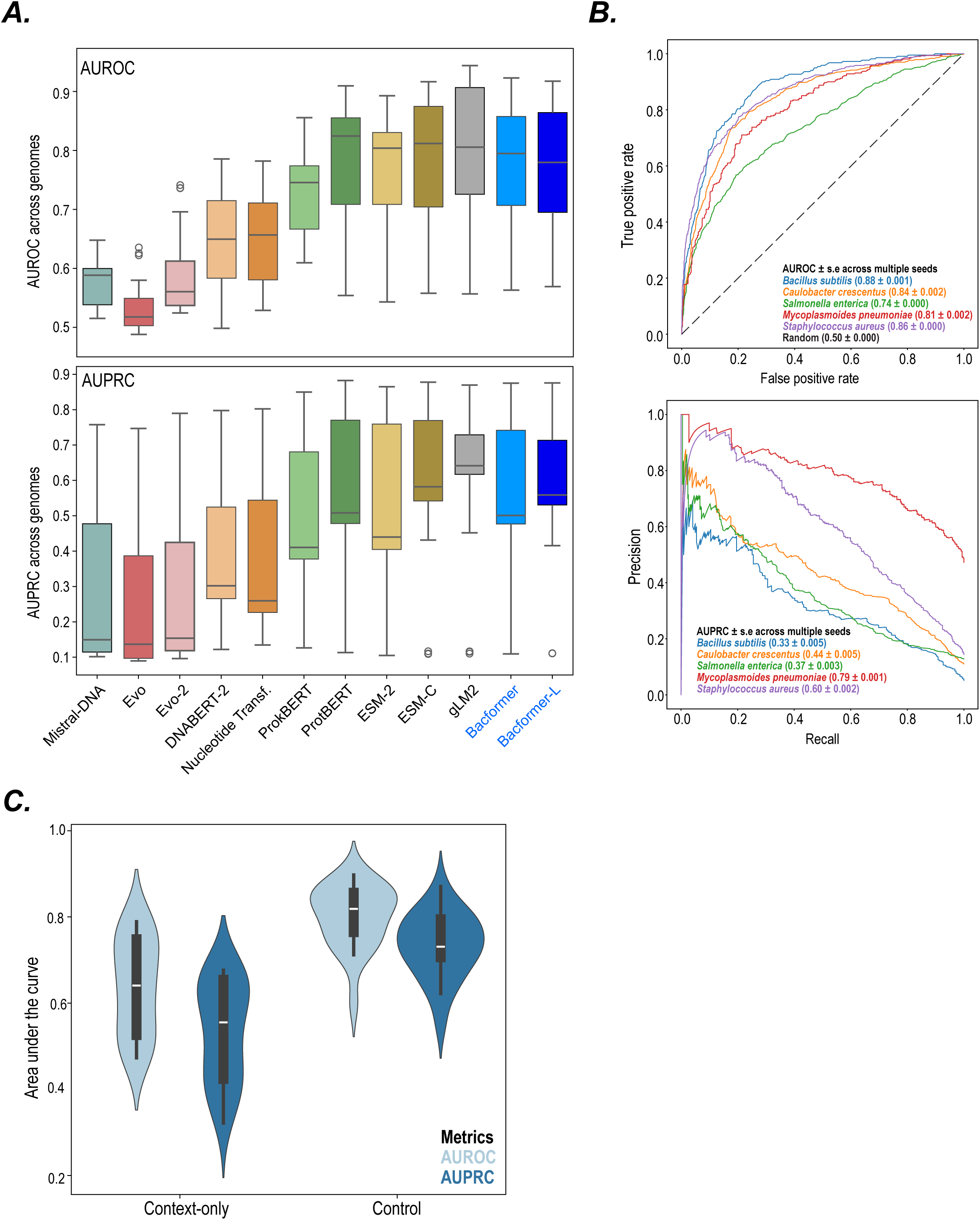
Gene essentiality prediction. (**A**) Performance across different methods on the gene essentiality prediction task across genomes on the held-out evaluation set (*n* = 11) evaluated by area under the Receiver Operating Characteristic (AUROC; *top*) and Precision-Recall (AUPRC; *bottom*) curves. We trained a linear layer using protein representations to predict gene essentiality for each method. Line and boxes represent median and interquartile range (IQR) respectively; whiskers extend 1.5 × IQR beyond Q1 and Q3. (**B**) Receiver Operating Characteristic (ROC; *top*) and Precision-Recall (PR; *bottom*) curves across diverse species computed using *Bacformer* finetuned for the gene essentiality task trained with different random seeds. (**C**) Finetuned *Bacformer* predicts gene essentiality based on context-only across diverse genomes. We masked the essential protein and asked the model to predict essentiality based only on the other proteins present in the genome (median and interquartile range shown).

**FIGURE S10.**
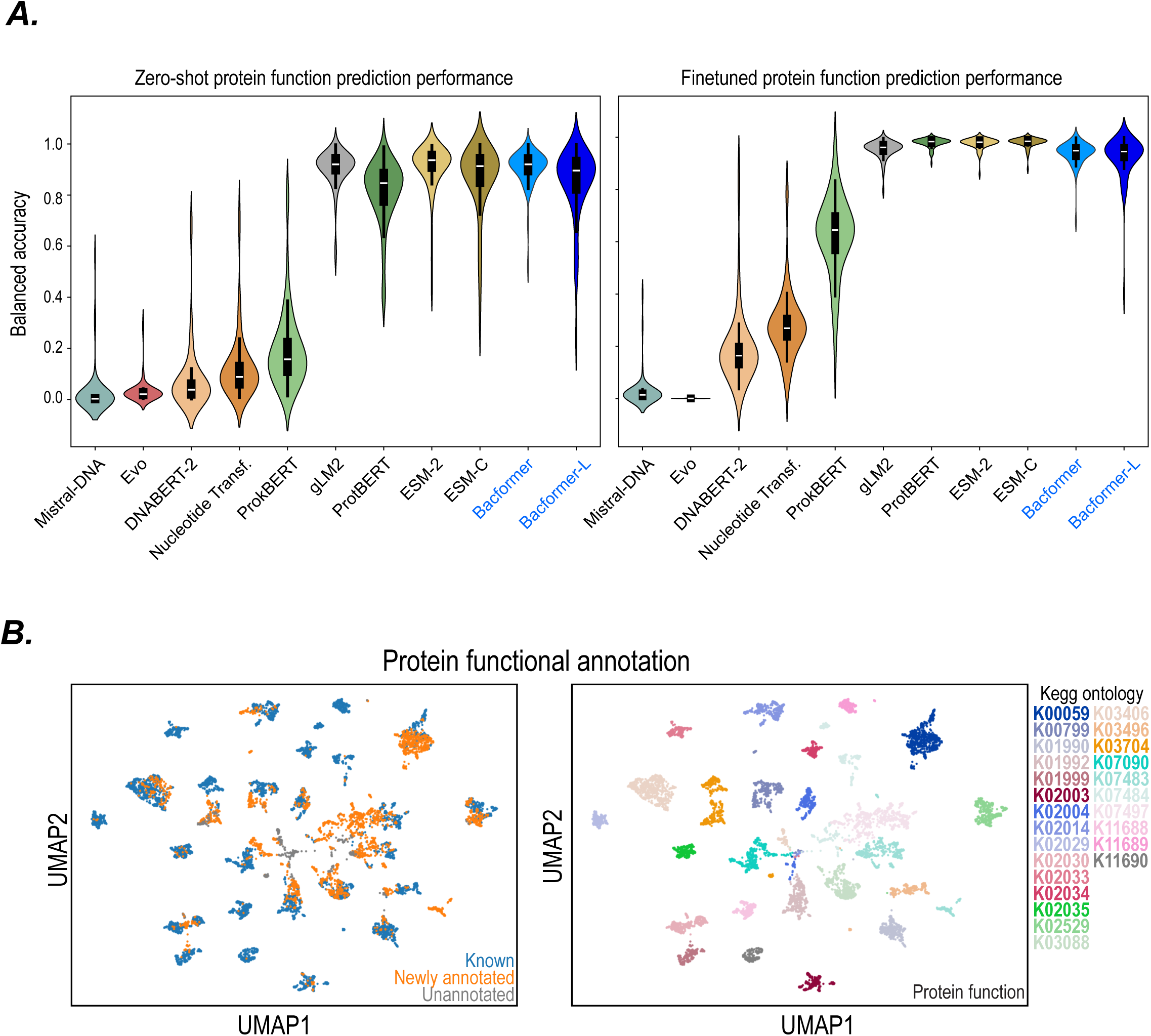
Protein function annotation prediction. (**A**) Performance across methods on the protein function annotation task on held-out genomes (**n** = 60) using balanced accuracy across unique protein functions (*N* = 9,059). In the zero-shot setting (*left*), we evaluate performance by using kNN and for finetuning (*right*) we train a linear layer on top of the protein representations. We used KEGG KO Orthology groups as ground truth (*see Supplementary Methods*). Violins show the kernel density; inner box and line show IQR and median, with whiskers extending to data within 1.5 × IQR of Q1 and Q3. (**B**) UMAP of proteins coloured by annotation status (*left*) and predicted KEGG Orthology group (KO). We used the proteins from top 25 most frequently predicted groups and annotated previously unannotated proteins based on the prediction from the finetuned *Bacformer-L* model, using a normalised prediction score as a threshold (*see Supplementary Methods*). We plotted the UMAP using Scanpy package using 15 neighbors for the kNN graph and minimum distance of 0.5 for the UMAP.

**FIGURE S11.**
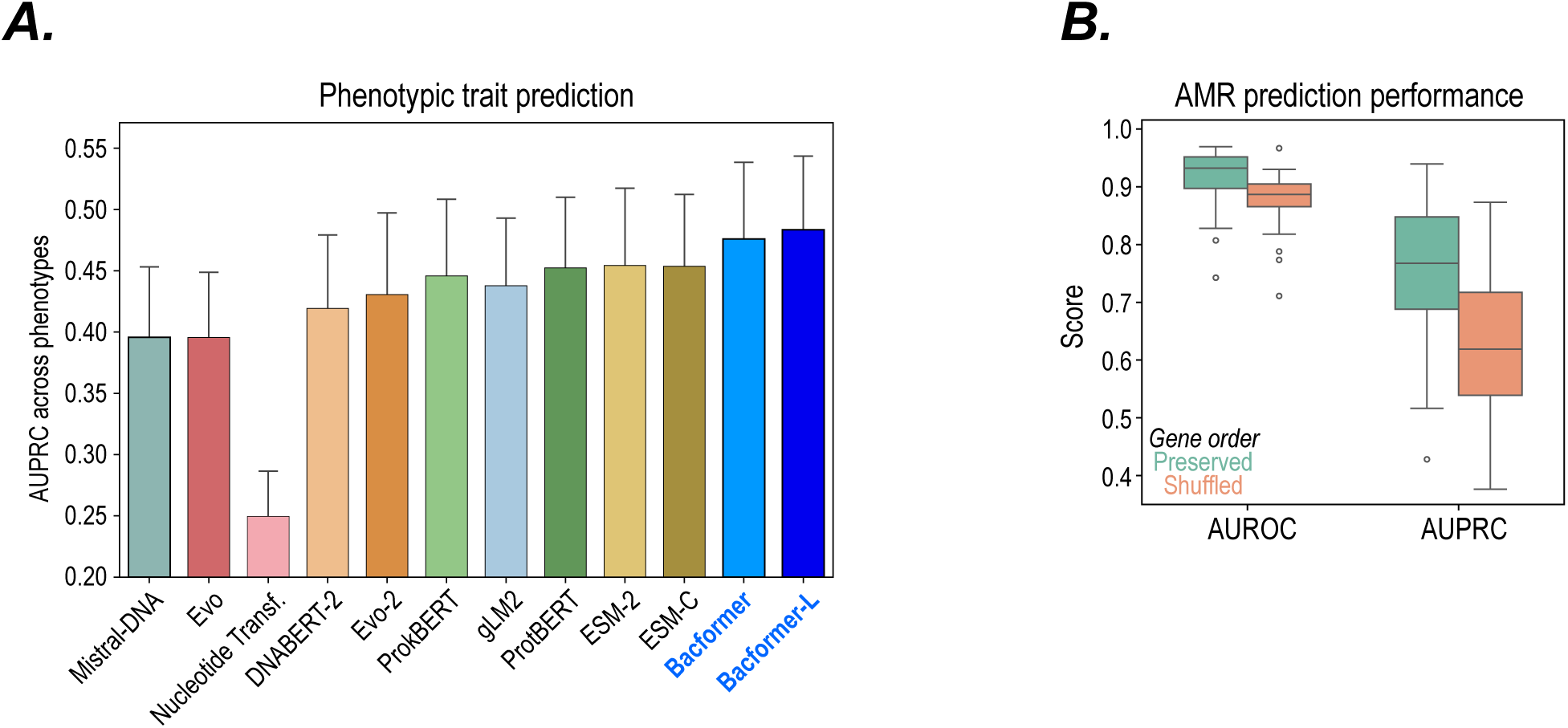
Additional benchmarking results on phenotype prediction tasks. (**A**) Benchmarking performance across methods on 138 distinct traits across random seeds as evaluated by the area under the precision-recall curve (AUPRC). For each method-trait combination we trained a linear layer on top of the genome embedding splitting the data by genus to ensure generalization across phylogenetically different bacteria (*see Supplementary Methods*). Error bars represent 95% confidence intervals across traits and random seeds. (**B**) Ablation study results on the AMR prediction task using *Bacformer-L* with preserved vs randomly shuffled gene order across distinct drugs (*N* = 14) and genomes (*N* = 6,000). Line and boxes represent median and interquartile range (IQR) respectively; whiskers extend 1.5 × IQR beyond Q1 and Q3.

**FIGURE S12.**
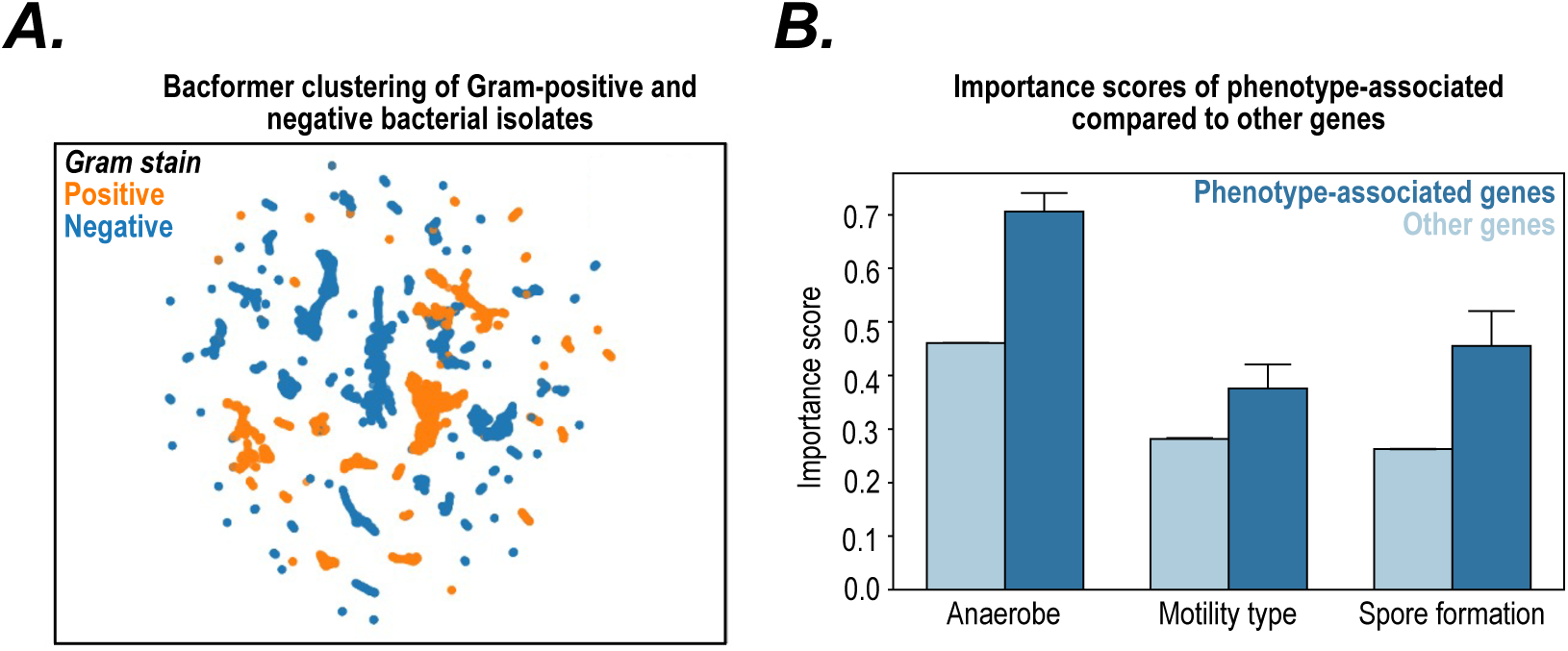
Bacformer prediction of phenotypic traits. (**A**) UMAP plot of gram-positive (*orange*) and negative (*blue*) strains. The genome embeddings were computed in a zero-shot manner using aggregated contextualised protein embeddings from the last layer of the pretrained *Bacformer* model. (**B**) *Bacformer* identifies phenotype-specific genes by computing importance scores per gene using phenotype-specific models. We used a trained phenotype-specific *Bacformer* model to extract gradient-based attribution scores per gene across samples. The genes with highest importance scores across samples are labelled “phenotype-specific genes”.

**FIGURE S13.**
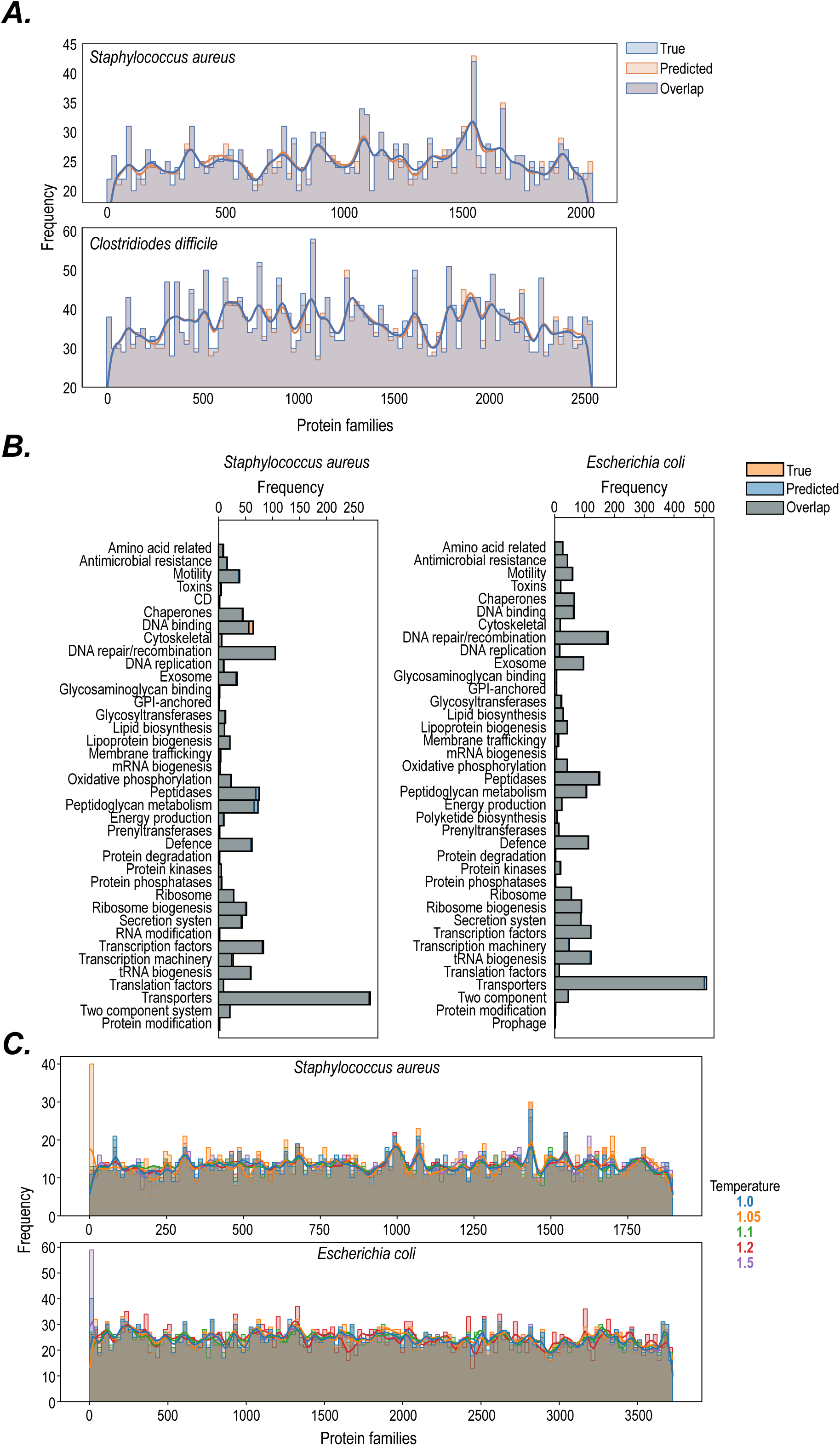
Generative modelling capacity of Bacformer. (**A**) *Bacformer* successfully generates genomes unseen during training. True (*blue*) and predicted (*orange*) protein family distributions shown for *Staphylococcus aureus* and *Clostridioides difficile*. (**B**) *Bacformer* generates genomes encompassing a broad range of functions. True (*blue*) and predicted (*orange*) protein function distribution on *S. aureus* (*left*) and *E. coli* (*right*). (**C**) *Bacformer* generates diverse protein family sequences. We prompted *Bacformer* with a set of 500 initial proteins on two different bacteria. The generated genomes varies across different temperature levels, which control the diversity of the generation process.

**FIGURE S14.**
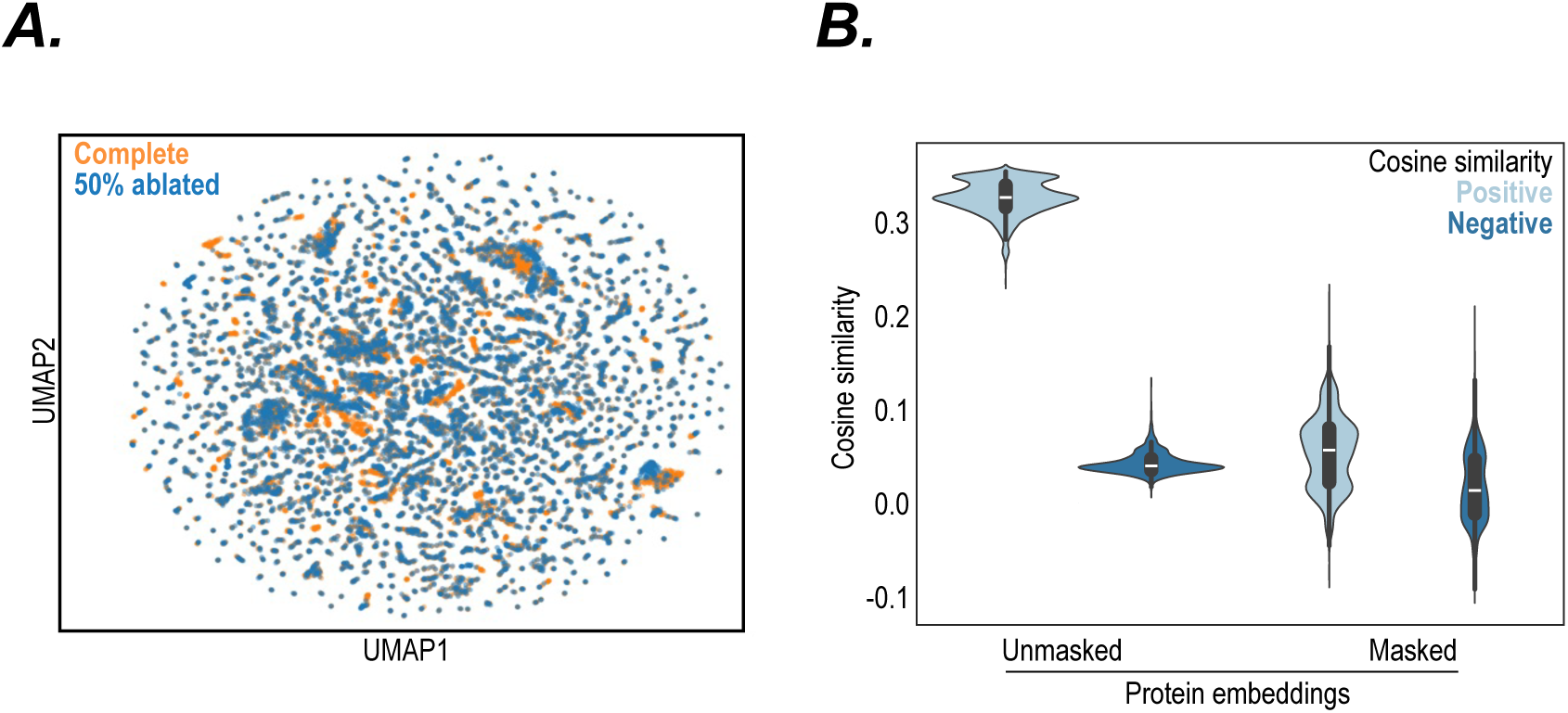
Retrieving complete genomes and real proteins from contextual Bacformer embeddings. (**A**) UMAP plot of *Bacformer* embeddings of complete genomes (*orange*) compared to the same genomes with 50% of the proteins artificially ablated (*blue*). (**B**) Cosine similarity between *Bacformer* contextual protein embeddings and *ESM-2* protein embeddings for masked and unmasked protein embeddings. The *ESM-2* embeddings are used as input to the *Bacformer* transformer block which computes contextual embeddings. The positive (*light blue*) represents the true protein used at input-level, while negatives (*dark blue*) represent other proteins in the genome (median and interquartile range shown). Statistical significance was assessed across held-out genomes using a two-sided Mann--Whitney U test (p < 10^−5^).

### SUPPLEMENTARY TABLES

**Table S1.** Overall operon identification performance across methods. Values are mean ± standard deviation; best value for each metric is bolded.

| Method | AUROC | AUPRC |
| --- | --- | --- |
| Mistral-DNA | 74.16 ± 9.70 | 47.48 ± 29.47 |
| Evo | 74.77 ± 8.33 | 48.21 ± 28.27 |
| Evo-2 | 71.63 ± 10.31 | 45.01 ± 29.14 |
| DNABERT-2 | 74.13 ± 10.14 | 47.53 ± 30.07 |
| Nucleotide Transformer | 74.50 ± 8.57 | 48.19 ± 27.47 |
| ProkBERT | 75.77 ± 9.69 | 49.03 ± 30.07 |
| ProtBERT | 75.11 ± 8.65 | 47.58 ± 27.98 |
| ESM-2 | 73.02 ± 10.30 | 46.27 ± 29.55 |
| ESM-C | 75.25 ± 9.26 | 47.79 ± 29.39 |
| gLM2 | 72.85 ± 10.77 | 45.86 ± 30.18 |
| Bacformer | **77.59 ± 6.64** | **50.31 ± 26.61** |
| Bacformer Large | 77.00 ± 7.46 | 49.98 ± 26.92 |

**Table S2.** Operon identification statistical analysis. Paired strain-level comparison on the operon identification task of Bacformer against benchmarked models. Mean differences are computed as comparator minus Bacformer, so negative values indicate better Bacformer performance. *p*-values were computed using a two-sided Wilcoxon test on *N* = 5 strains.

| Method | ΔAUROC | $p_{\text{AUROC}}$ | ΔAUPRC | $p_{\text{AUPRC}}$ |
| --- | --- | --- | --- | --- |
| DNABERT-2 | -0.0346 | 0.3125 | -0.0278 | 0.3125 |
| ESM-2 | -0.0457 | 0.0625 | -0.0404 | 0.0625 |
| ESM-C | -0.0234 | 0.1875 | -0.0252 | 0.1875 |
| gLM2 | -0.0474 | 0.1250 | -0.0445 | 0.1250 |
| Mistral-DNA | -0.0343 | 0.1250 | -0.0283 | 0.1875 |
| Nucleotide Transformer | -0.0309 | 0.0625 | -0.0212 | 0.0625 |
| ProkBERT | -0.0182 | 0.4375 | -0.0128 | 0.6250 |
| ProtBERT | -0.0248 | 0.1250 | -0.0273 | 0.1250 |
| Evo | -0.0283 | 0.0625 | -0.0210 | 0.1250 |
| Evo-2 | -0.0596 | 0.0625 | -0.0530 | 0.0625 |
| Bacformer large | -0.0059 | 0.4375 | -0.0033 | 0.4375 |

**Table S3. (separate file)** Gene essentiality prediction results for *Staphylococcus aureus* (NC_002952) from *Bacformer* model across random seeds.

**Table S4. (separate file)** Gene essentiality prediction results for *Salmonella enterica* (NC_003197) from *Bacformer* model across random seeds.

**Table S5. (separate file)** Environment-specific genes for *Lactobacillus johnsonii* across human, pig and chicken gut based on gradient saliency using *Bacformer* model.

**Tables S6-S8. (separate files)** Attribution Scores for AMR Finder Plus elements for *E. coli* resistant to *ampicillin* (Table S6), *A. baumannii* resistant to *gentamicin* (Table S7) and *Salmonella enterica* resistant to *tetracycline* (Table S8). Each table has rows for all AMR Finder Plus elements (including AMR and non-AMR related subtypes and AMR classes causal to the antibiotic and other AMR classes). The rows are ordered by the Rank Biserial (a measure of the rankings of attributions of genes in the group compared to all genes). The rate of resistance is defined as the percentage of genomes with the gene that are resistant to the drug. Element penetrance is defined by the percentage of genomes that have at least one copy of the gene (number of genomes with gene / total number of genomes). We note that the number of proteins per symbol may be higher than the number of genomes, reflecting multiple (duplicate or similar) genes with the same element symbol in the genome.

**Table S9.**
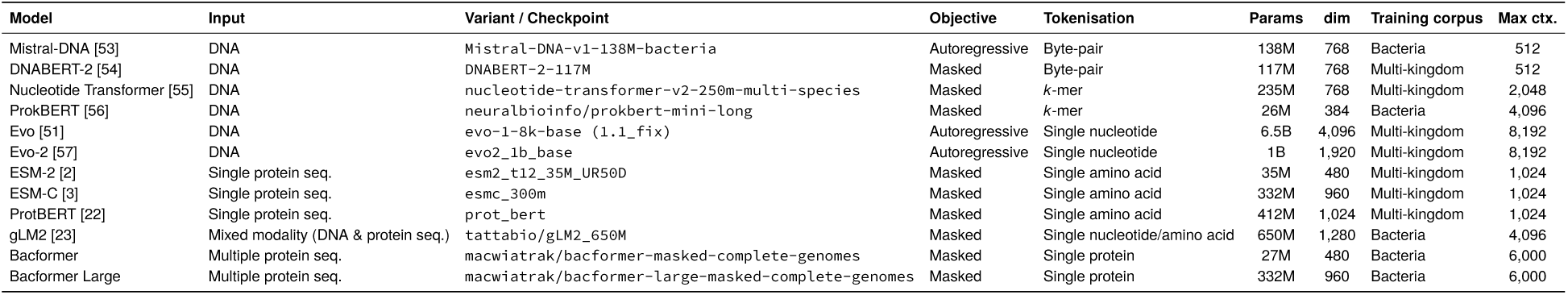
Summary of benchmarked models. “Max ctx.” = maximum context length supported at inference; for DNA, pLM and mixed-modality models it is reported in tokens, and for Bacformer in number of proteins present in the genome. “dim” = dimensionality of the output of the last hidden layer.

**Table S10.** Model-specific context parameters used in this study. “Max ctx.” is the maximum input length; “DNA-seq overlap” is the stride between consecutive windows when sliding across a genome; “Promoter length” is the upstream sequence length concatenated with the gene body used for DNA LMs.* For Bacformer the maximum input size of a protein is 1, 024 amino acids and maximum number of proteins in the genome is set to 6, 000.

| Model | Max ctx. | DNA-seq overlap | Promoter length |
| --- | --- | --- | --- |
| Mistral-DNA | 512 | 16 | 128 |
| DNABERT-2 | 512 | 16 | 128 |
| Nucleotide Transformer | 2,048 | 32 | 128 |
| ProkBERT | 4,096 | 64 | 128 |
| Evo | 8,192 | 32 | 128 |
| Evo-2 | 8,192 | 32 | 128 |
| ProtBERT | 1,024 | N/A | N/A |
| ESM-2 | 1,024 | N/A | N/A |
| ESM-C | 1,024 | N/A | N/A |
| gLM2 | 4,096 | 64 | 128 |
| Bacformer | 1,024 / 6,000* | N/A | N/A |
| Bacformer Large | 1,024 / 6,000* | N/A | N/A |

**Table S11.** Total GPU-hours required for embedding genomic data in BacBench using a single NVIDIA H100 Tensor Core GPU with 72 CPU cores. Values are reported in GPU-hours per model and task; the final column sums across tasks.

| Model | Essential gene | Operon | PPI | Phenotypic traits | Antibiotic resistance | Total GPU hours |
| --- | --- | --- | --- | --- | --- | --- |
| Mistral-DNA (138M) | 0.11 | 0.03 | 0.55 | 33.96 | 36.53 | 71.18 |
| Evo (6.5B) | 20.92 | 4.77 | 100.82 | 1426.37 | 1534.44 | 3087.32 |
| Evo-2 (1B) | 4.58 | 1.04 | 22.07 | 365.37 | 393.05 | 786.12 |
| DNABERT-2 (117M) | 0.20 | 0.05 | 0.99 | 61.35 | 66.00 | 128.58 |
| Nucleotide Transformer (235M) | 0.12 | 0.03 | 0.59 | 50.26 | 54.07 | 105.06 |
| ProkBERT (26M) | 0.13 | 0.03 | 0.62 | 84.22 | 90.60 | 175.60 |
| ProtBERT (412M) | 0.12 | 0.03 | 0.56 | 59.97 | 62.97 | 123.65 |
| ESM-2 (35M) | 0.08 | 0.02 | 0.38 | 40.40 | 42.42 | 83.30 |
| ESM-C (332M) | 0.17 | 0.04 | 0.80 | 86.49 | 90.81 | 178.31 |
| gLM2 (650M) | 0.22 | 0.05 | 1.07 | 142.12 | 0.00 | 143.46 |
| Bacformer (26M) | 0.08 | 0.02 | 0.39 | 42.30 | 44.41 | 87.20 |
| Bacformer Large (332M) | 0.18 | 0.04 | 0.85 | 91.54 | 96.12 | 188.73 |
| **Total** | **27.11** | **6.18** | **130.65** | **2622.97** | **2683.77** | **5470.68** |

**Table S12.**
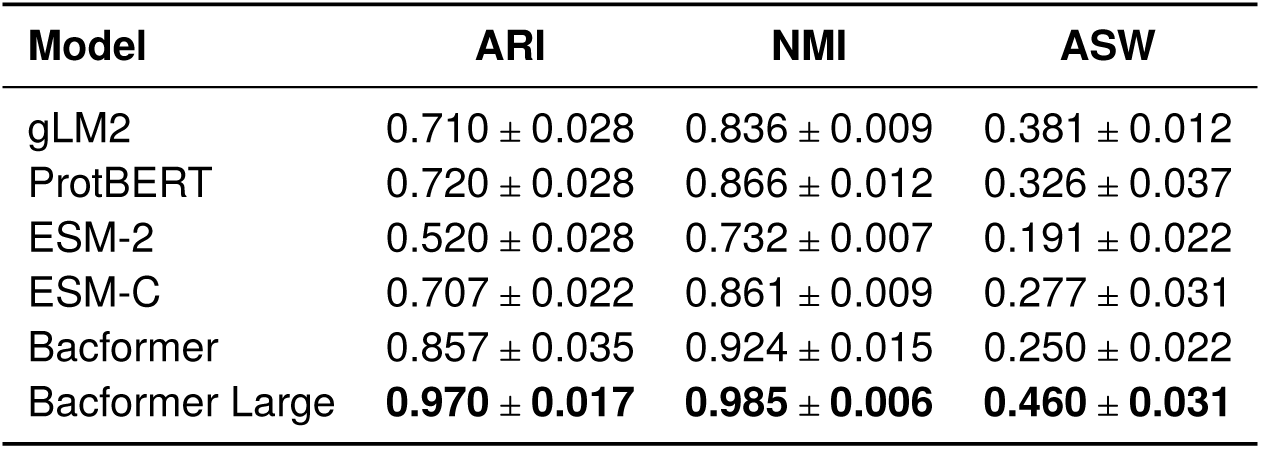
Clustering performance across species-level genome clustering, reported as Adjusted Rand Index (ARI), Normalised Mutual Information (NMI), and Average Silhouette Width (ASW). Best value for each metric is bolded. Mean ± standard deviation, where standard deviation was computed by performing 10 independent bootstrap resamples, each using 80% of the data.

| Model | ARI | NMI | ASW |
| --- | --- | --- | --- |
| gLM2 | 0.710 ± 0.028 | 0.836 ± 0.009 | 0.381 ± 0.012 |
| ProtBERT | 0.720 ± 0.028 | 0.866 ± 0.012 | 0.326 ± 0.037 |
| ESM-2 | 0.520 ± 0.028 | 0.732 ± 0.007 | 0.191 ± 0.022 |
| ESM-C | 0.707 ± 0.022 | 0.861 ± 0.009 | 0.277 ± 0.031 |
| Bacformer | 0.857 ± 0.035 | 0.924 ± 0.015 | 0.250 ± 0.022 |
| Bacformer Large | **0.970 ± 0.017** | **0.985 ± 0.006** | **0.460 ± 0.031** |

**Table S13.**
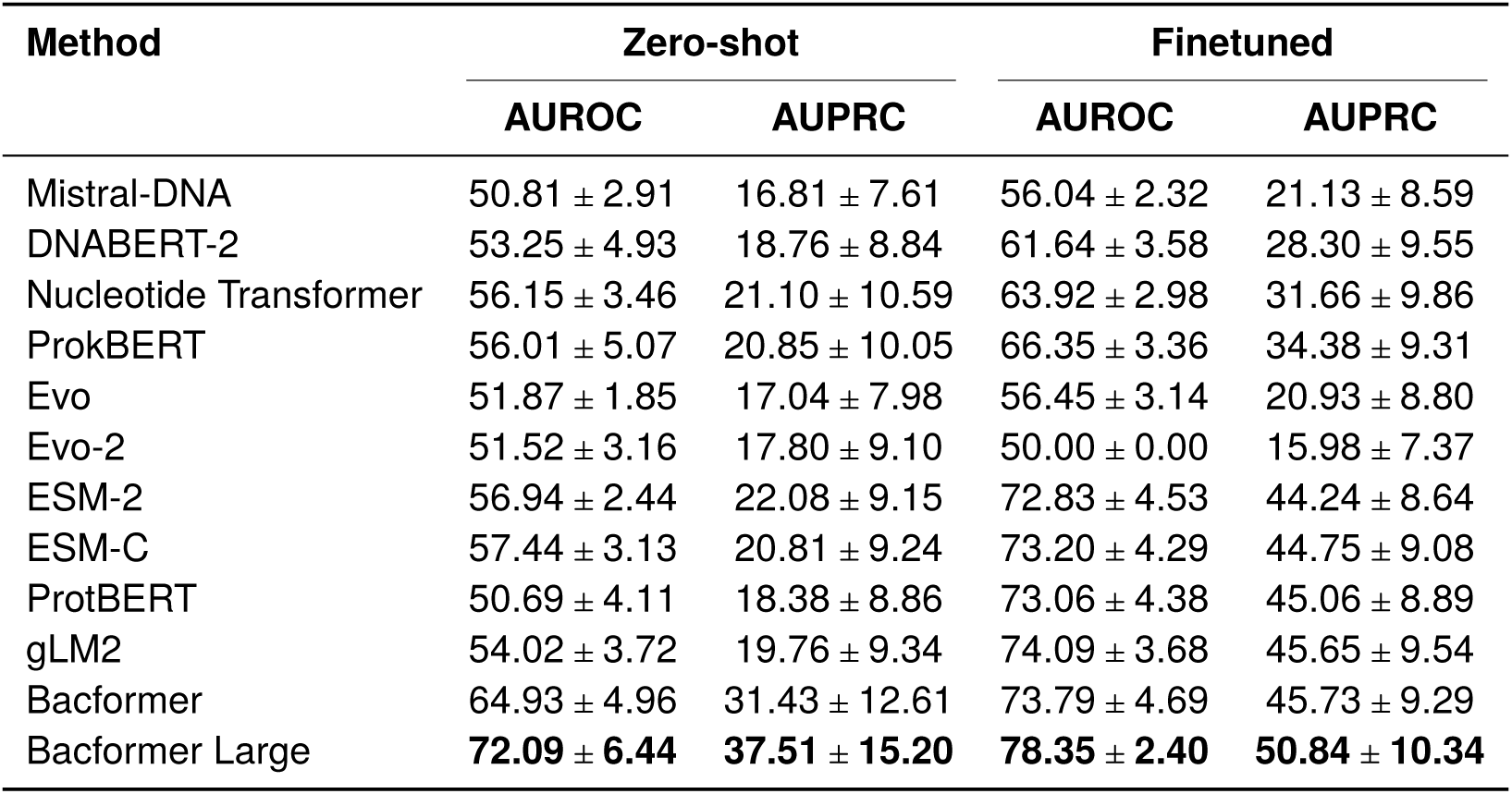
Protein–protein interaction performance on held-out genomes. Values are mean ± standard deviation across test genomes; best value for each metric is bolded.

| Method | Zero-shot |  | Finetuned |  |
| --- | --- | --- | --- | --- |
|  | AUROC | AUPRC | AUROC | AUPRC |
| Mistral-DNA | 50.81 ± 2.91 | 16.81 ± 7.61 | 56.04 ± 2.32 | 21.13 ± 8.59 |
| DNABERT-2 | 53.25 ± 4.93 | 18.76 ± 8.84 | 61.64 ± 3.58 | 28.30 ± 9.55 |
| Nucleotide Transformer | 56.15 ± 3.46 | 21.10 ± 10.59 | 63.92 ± 2.98 | 31.66 ± 9.86 |
| ProkBERT | 56.01 ± 5.07 | 20.85 ± 10.05 | 66.35 ± 3.36 | 34.38 ± 9.31 |
| Evo | 51.87 ± 1.85 | 17.04 ± 7.98 | 56.45 ± 3.14 | 20.93 ± 8.80 |
| Evo-2 | 51.52 ± 3.16 | 17.80 ± 9.10 | 50.00 ± 0.00 | 15.98 ± 7.37 |
| ESM-2 | 56.94 ± 2.44 | 22.08 ± 9.15 | 72.83 ± 4.53 | 44.24 ± 8.64 |
| ESM-C | 57.44 ± 3.13 | 20.81 ± 9.24 | 73.20 ± 4.29 | 44.75 ± 9.08 |
| ProtBERT | 50.69 ± 4.11 | 18.38 ± 8.86 | 73.06 ± 4.38 | 45.06 ± 8.89 |
| gLM2 | 54.02 ± 3.72 | 19.76 ± 9.34 | 74.09 ± 3.68 | 45.65 ± 9.54 |
| Bacformer | 64.93 ± 4.96 | 31.43 ± 12.61 | 73.79 ± 4.69 | 45.73 ± 9.29 |
| Bacformer Large | **72.09 ± 6.44** | **37.51 ± 15.20** | **78.35 ± 2.40** | **50.84 ± 10.34** |

**Table S14.**
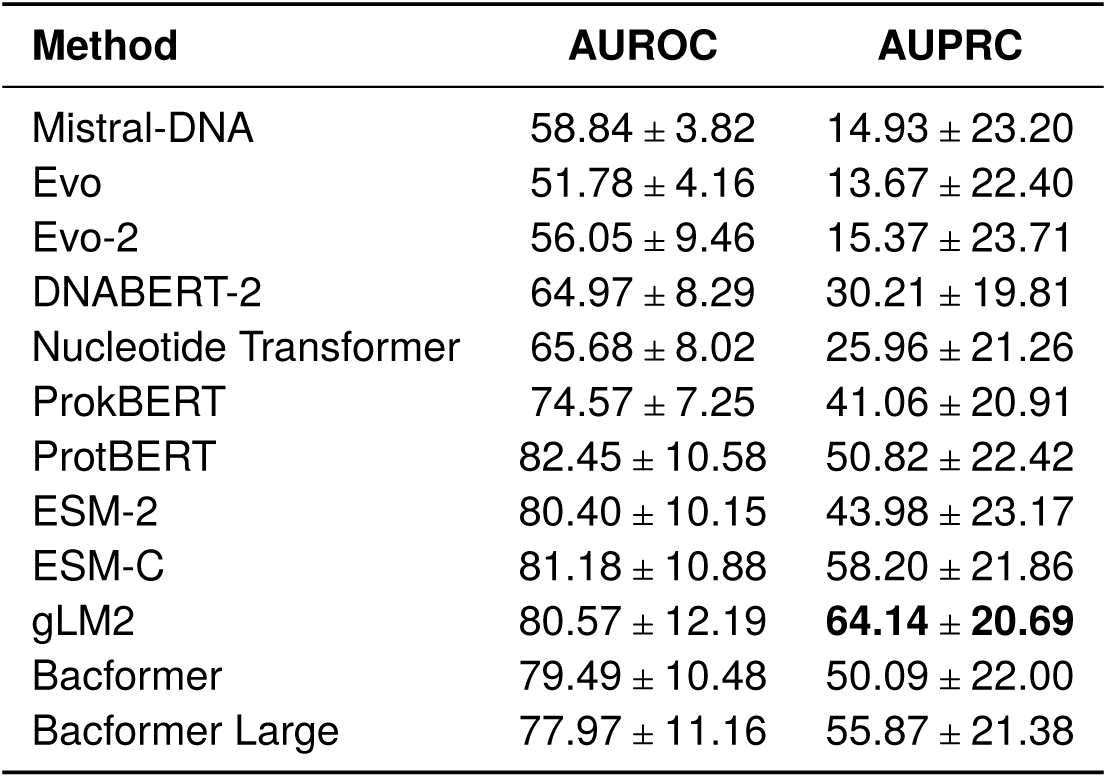
Performance on gene–essentiality prediction. Values are *median* ± *standard deviation* over test genomes; the best score for each metric is highlighted in bold.

| Method | AUROC | AUPRC |
| --- | --- | --- |
| Mistral-DNA | 58.84 ± 3.82 | 14.93 ± 23.20 |
| Evo | 51.78 ± 4.16 | 13.67 ± 22.40 |
| Evo-2 | 56.05 ± 9.46 | 15.37 ± 23.71 |
| DNABERT-2 | 64.97 ± 8.29 | 30.21 ± 19.81 |
| Nucleotide Transformer | 65.68 ± 8.02 | 25.96 ± 21.26 |
| ProkBERT | 74.57 ± 7.25 | 41.06 ± 20.91 |
| ProtBERT | 82.45 ± 10.58 | 50.82 ± 22.42 |
| ESM-2 | 80.40 ± 10.15 | 43.98 ± 23.17 |
| ESM-C | 81.18 ± 10.88 | 58.20 ± 21.86 |
| gLM2 | 80.57 ± 12.19 | **64.14 ± 20.69** |
| Bacformer | 79.49 ± 10.48 | 50.09 ± 22.00 |
| Bacformer Large | 77.97 ± 11.16 | 55.87 ± 21.38 |

**Table S15.** Protein function prediction performance on the held-out test set (higher is better), median ± standard deviation across strains. Best value for each metric is bolded.

| Method | Zero-shot |  | finetuned |  |
| --- | --- | --- | --- | --- |
|  | Macro F1 | Macro accuracy | Macro F1 | Macro accuracy |
| Mistral-DNA | 0.10 ± 7.90 | 0.27 ± 9.07 | 0.70 ± 5.35 | 1.38 ± 6.12 |
| Evo | 1.23 ± 3.66 | 2.03 ± 5.03 | 0.80 ± 4.25 | 1.20 ± 5.75 |
| DNABERT-2 | 2.50 ± 11.66 | 3.78 ± 13.04 | 11.45 ± 14.17 | 16.40 ± 15.00 |
| Nucleotide Transformer | 6.06 ± 11.70 | 8.82 ± 13.22 | 19.73 ± 12.33 | 26.89 ± 12.93 |
| ProkBERT | 11.78 ± 14.12 | 15.71 ± 15.44 | 55.23 ± 16.32 | 64.45 ± 15.36 |
| gLM2 | **90.06 ± 12.04** | **93.66 ± 9.90** | 97.01 ± 3.38 | 98.06 ± 2.21 |
| ProtBERT | 78.26 ± 14.47 | 84.65 ± 12.65 | 97.24 ± 2.95 | 98.31 ± 1.80 |
| ESM-2 | 87.85 ± 9.88 | 92.04 ± 7.58 | 93.48 ± 5.04 | 96.06 ± 3.31 |
| ESM-C | 87.58 ± 16.60 | 91.34 ± 14.52 | **97.35 ± 2.92** | **98.41 ± 1.96** |
| Bacformer | 87.54 ± 9.34 | 92.04 ± 7.15 | 91.92 ± 6.39 | 94.82 ± 4.52 |
| Bacformer Large | 84.55 ± 16.40 | 89.59 ± 14.22 | 91.94 ± 10.34 | 94.50 ± 8.60 |

**Table S16.** Overall phenotypic traits prediction performance across all phenotypes. Values are mean ± standard deviation across 5 seeds. The results were macro-averaged across classes for each phenotype. Best value for each metric is bolded.

| Method | AUROC | AUPRC |
| --- | --- | --- |
| Mistral-DNA | 60.33 ± 9.88 | 39.56 ± 33.16 |
| DNABERT-2 | 63.81 ± 11.15 | 41.93 ± 34.31 |
| Nucleotide Transformer | 65.98 ± 11.54 | 43.05 ± 34.91 |
| ProkBERT | 67.58 ± 11.94 | 44.58 ± 34.78 |
| Evo | 60.19 ± 11.04 | 39.55 ± 33.36 |
| Evo-2 | 55.41 ± 8.85 | 24.96 ± 20.52 |
| ESM-2 | 68.68 ± 12.46 | 45.42 ± 35.54 |
| ESM-C | 69.06 ± 12.07 | 45.36 ± 35.47 |
| ProtBERT | 68.40 ± 12.05 | 45.24 ± 34.98 |
| gLM2 | 65.74 ± 11.47 | 43.77 ± 34.45 |
| Bacformer | 71.35 ± 13.01 | 47.60 ± 36.13 |
| Bacformer Large | **73.33 ± 12.13** | **48.34 ± 36.39** |

**Table S17.** Number of resistant, susceptible, and total labelled genomes for every antibiotic in the binary prediction setting.

| Drug | # Resistant | # Susceptible | # Total |
| --- | --- | --- | --- |
| amikacin | 7353 | 588 | 7941 |
| ampicillin | 10052 | 6211 | 16263 |
| amoxicillin–clavulanic acid | 12458 | 1683 | 14141 |
| azithromycin | 10497 | 2509 | 13006 |
| cefazolin | 1666 | 1897 | 3563 |
| cefepime | 3237 | 1188 | 4425 |
| cefotaxime | 566 | 446 | 1012 |
| cefoxitin | 12463 | 3902 | 16365 |
| ceftazidime | 2362 | 717 | 3079 |
| ceftriaxone | 13385 | 3469 | 16854 |
| ciprofloxacin | 17353 | 4280 | 21633 |
| clindamycin | 4327 | 337 | 4664 |
| ertapenem | 7708 | 373 | 8081 |
| erythromycin | 5605 | 1860 | 7465 |
| fosfomycin | 1186 | 396 | 1582 |
| gentamicin | 18242 | 2951 | 21193 |
| imipenem | 8485 | 409 | 8894 |
| kanamycin | 3768 | 410 | 4178 |
| levofloxacin | 3302 | 1106 | 4408 |
| meropenem | 11320 | 811 | 12131 |
| nalidixic acid | 1262 | 780 | 2042 |
| nitrofurantoin | 10535 | 5316 | 15851 |
| oxacillin | 6929 | 762 | 7691 |
| piperacillin–tazobactam | 3294 | 968 | 4262 |
| rifampin | 611 | 354 | 965 |
| streptomycin | 11965 | 2442 | 14407 |
| sulfamethoxazole | 3770 | 707 | 4477 |
| sulfisoxazole | 1569 | 323 | 1892 |
| tetracycline | 6936 | 3242 | 10178 |
| tigecycline | 662 | 357 | 1019 |
| trimethoprim | 2617 | 582 | 3199 |
| ampicillin–sulbactam | 1632 | 451 | 2083 |
| aztreonam | 2530 | 728 | 3258 |
| ceftaroline | 611 | 152 | 763 |
| chloramphenicol | 5345 | 1016 | 6361 |
| colistin | 572 | 290 | 862 |
| daptomycin | 492 | 131 | 623 |

**Table S18.** Performance (percent, higher is better) on antibiotic-resistance prediction tasks across drugs. Values are mean ± standard deviation across antibiotics; best value for each metric is bolded.

| Method | Binary |  | Regression |  |
| --- | --- | --- | --- | --- |
|  | AUROC | AUPRC | R<sup>2</sup> | Pearson |
| Mistral-DNA | 76.05 ± 9.14 | 49.86 ± 22.73 | 19.71 ± 17.37 | 40.95 ± 21.51 |
| DNABERT-2 | 79.88 ± 7.26 | 52.35 ± 23.55 | 23.61 ± 18.33 | 45.89 ± 19.27 |
| Nucleotide Transformer | 84.22 ± 6.47 | 59.70 ± 22.90 | 27.74 ± 17.80 | 51.19 ± 16.76 |
| ProkBERT | 83.94 ± 6.39 | 58.90 ± 23.67 | 28.00 ± 17.57 | 51.24 ± 16.91 |
| Evo | 68.22 ± 9.32 | 44.06 ± 23.28 | 10.01 ± 10.06 | 29.18 ± 17.26 |
| Evo-2 | 66.27 ± 10.19 | 38.35 ± 23.08 | 03.95 ± 21.04 | 00.53 ± 00.72 |
| ESM-2 | 85.05 ± 6.63 | 62.04 ± 23.64 | 31.18 ± 17.39 | 54.60 ± 16.15 |
| ESM-C | 85.68 ± 6.81 | 63.41 ± 23.43 | 33.15 ± 17.62 | 56.79 ± 15.46 |
| ProtBERT | 84.51 ± 6.31 | 61.79 ± 23.19 | 28.23 ± 18.46 | 51.56 ± 17.61 |
| gLM2 | 82.16 ± 6.75 | 55.80 ± 24.46 | 26.15 ± 18.13 | 49.41 ± 17.14 |
| Bacformer | 87.61 ± 5.68 | 67.97 ± 20.73 | 33.84 ± 18.60 | 57.53 ± 15.43 |
| Bacformer Large | **92.15 ± 4.20** | **76.14 ± 19.59** | **42.59 ± 19.54** | **65.73 ± 12.88** |

